# Moving Yeasts: Resolving the Mystery

**DOI:** 10.1101/2022.03.05.483086

**Authors:** Gulab Puri, Sulbha Chaudhari, Areeb Inamdar, Manish Kohli, Vishal Sangawe, Chandrakant Jadhav, Lakshman Teja, Nitin Adhapure

## Abstract

We have recently discovered a novel dimorphic yeast *Aureobasidium tremulum* sp.nov. The species name tremulum indicates its trembling movement. With keen microscopic observations and various experimentation, we concluded that the movement is of two types, trembling movement, and actual displacement. The movement is because of two reasons; 1. Due to the dynamic movement of intracellular lipid granules, and; 2. Due to the movement of an intracellular bacterial population which is present as an endosymbiont. The mechanism of displacement has been hypothesized. The presence of both lipid granules and bacteria has been confirmed using microscopic analysis (Bright field, Phase contrast, confocal, Cryo-SEM, and HR-TEM). The bacterial presence was confirmed using whole-genome metagenomic analysis. Endosymbiotic bacteria have also been isolated and identified by 16s rRNA gene sequencing. Interestingly, some of them possibly are novel species. Also, the whole genome of *A.tremulum* has been announced. We also report on, restriction of the mode of bacterial mycophagy in presence of an antibiotic.

## Introduction

Since yeast does not have flagella or cilia-like locomotory organs, it is well - accepted truth that yeasts are non-motile^1^. The findings in the present manuscript allow us to rethink about this truth.

We have recently discovered a novel dimorphic yeast based on its trembling behavior it has been allotted with the name *Aureobasidium tremulum sp. nov*. This has been reported recently^2^..Though in one way we had succeeded in reporting a new species still we didn’t get the answer for the trembling behavior of yeast. The recent report also claims observation of nanomotions of yeast^3^. However, the displacement observed was at the nanoscale. In the present study, we mainly focused on proving the movement of yeast by designing suitable experiments and using high throughput analytical techniques. Subsequently, we have also focused on finding the reasons for the yeast movement. Since this is the first report of yeast movement we have used a polyphasic approach to prove the same. Based on extensive light microscopic observations we have hypothesized that the movement of *A.tremulum* is because of two reasons; 1. Due to the movement of endocellular bacteria, and/or 2. Due to the movement of intracellular lipid granules. a Moreover, the whole genome of *A.tremulum* has also been announced.

## Results and Discussion

### About *A.tremulum*

*A.tremulum* colonies on the agar plate are brownish to dark brown colored. The colonies appear to be rough and dry. Each colony is round with a convex elevation from a cross-sectional viewpoint and the edges appear to be undulated. Growth is optimal on Saboraud Dextrose Agar (SDA). Cells are generally oblong-shaped with very few cells assuming an irregular shape occasionally. Budding occurs frequently. The average size of mature, non-budding cells when measured on a slide under a specialized microscope is 6.4 by 2.8 μm. Sexual reproduction was not been observed. Pseudo hyphae formation did not occur. Growth occurred at 25^0^C^2^.

Growth of *A.tremulum* is insignificant and suboptimal at 5, 10, and 15 degrees celsius. The strain shows significant growth only within the temperature range of 20 to 25 degrees celsius. The culture was deposited at NCMR, Pune, India, and has been allotted with accession number MCC 1683^2^.

### Observation of growth pattern of dimorphic yeast on a different medium

Yeast grows rapidly on potato dextrose agar (PDA) medium at 25°C as compared to nutrient agar (NA) and malt extract agar (MEA) medium. White colonies of yeast turn into brownish color (due to the melanin like metabolite production)^4^, after 7 to 8 days of incubation at a constant temperature. Such browning of colonies can be observed on PDA and MEA but not on NA. With increasing incubation time, there is an increase in the number of yeast cells, the number of bacterial cells, and the number of lipid granules, as evident by microscopic observations. With increasing browning of a medium, lipid granules also increase in number.

After three to four days of incubation potatoes, dextrose broth (PDB) becomes highly viscous due to polysaccharide production by *A.tremulum*.

### Slide culture technique

In the slide culture technique, the growth pattern of yeast was observed directly on the agar block under a light microscope at 450x. Convertible form of mycelia into the yeast and arrangement of yeast in bunches was observed (Supplementary Fig. 1. 1).

### Motility assay

Brown-colored growth occurred on soft PDA medium at 25°C after two days of incubation. Yeast colonies appeared to be moving from point of stabbed medium to away from the stabbed point. Microscopically, bacteria and lipids were present within a yeast (Supplementary Fig. 4).

### Observing Yeast, Bacteria, and Lipid Granules

In addition to mycelial and yeast form, circular morphology was also observed (Fig. 3a). The yeast form was found to be surrounded by bacteria^5^. The bacterial motility was easily observed using a light microscope. While the bacteria were moving, all morphological forms of yeast were also found to be dancing. The possibility of movement due to air current and or movement due to water current was eliminated by sealing the coverslip using sticky tape.

**Figure 1.**
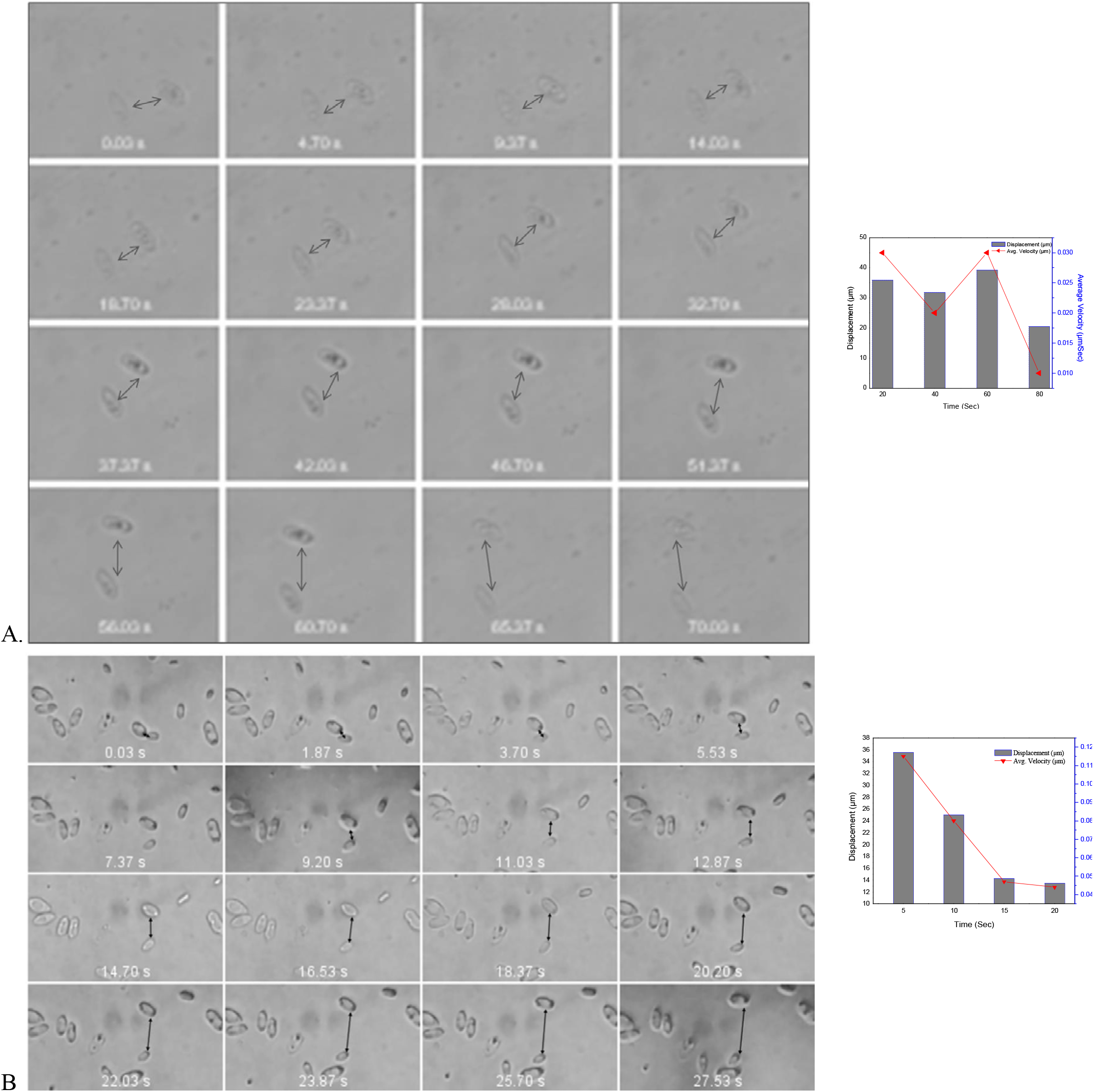
Displacement of yeast cells. Time lapsed images of light microscopic view of *A.tremulum*. Displacement of single yeast cell can be observed, while other yeast cells trembles at same position. Video files with cell tracking, is mentioned as supplementary videos 1 and 2 respectively. The distance travelled and the velocity is mentioned inadjescent graphs and in detail in supplementary files 1 and 2 respectively.

**Figure 2.**
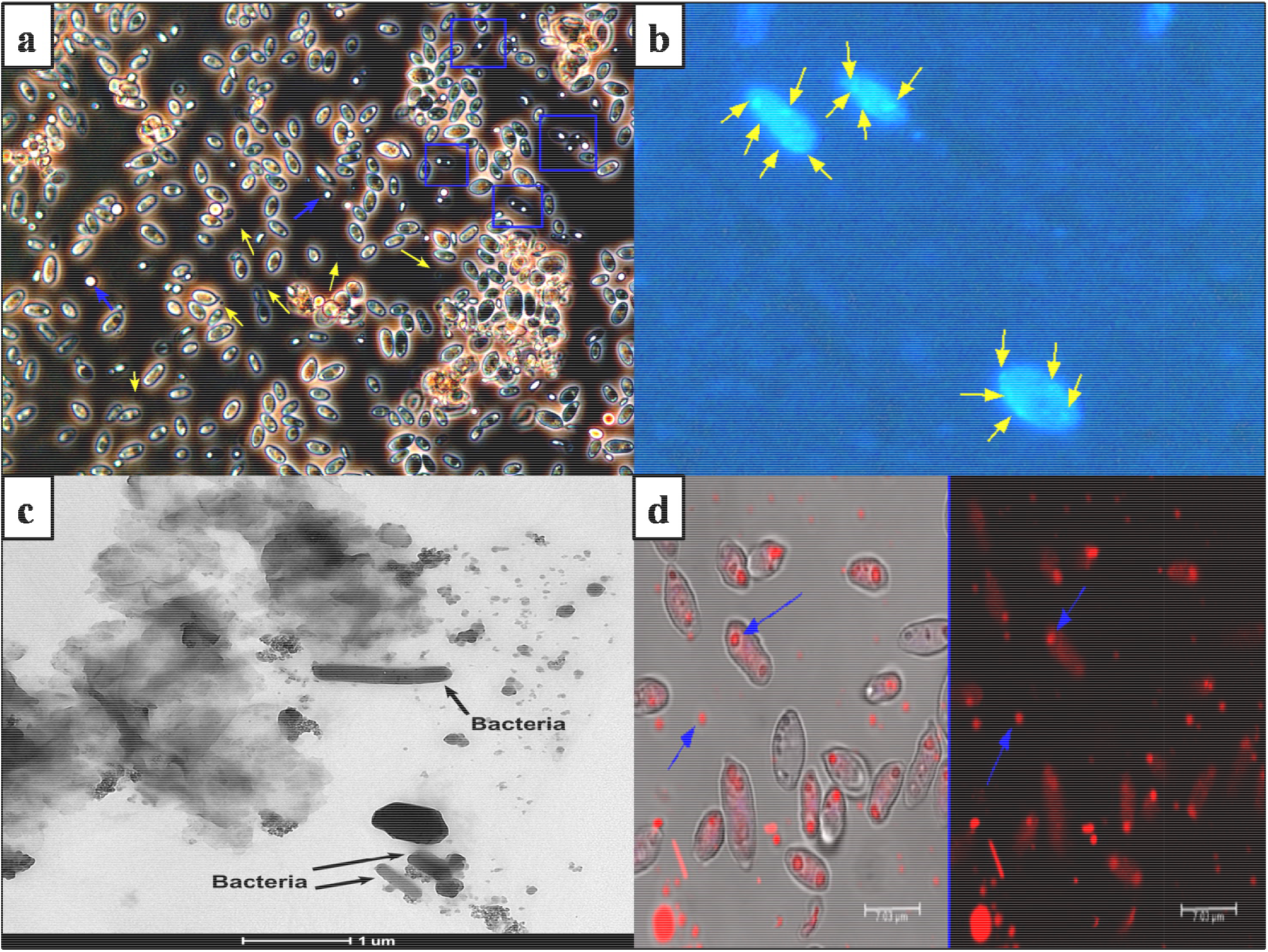
Presence of Bacteria and Lipid droplets in *A. tremulum*. **a.** Phase contrast image wherein Yellow arrow indicates bacteria, Blue arrow indicates extracellular lipid droplets, blue box indicates lipid droplets inside *A.tremulum*. **b.** Fluroscent image using DAPI wherein yellow arrow indicates bacteria inside *A.tremulum*. **c.** HR- TEM image wherein black arrow indicates bacteria released from bursted *A.tremulum* cells. **d.** Confocal microscopic image using Oil red O wherein, blue arrow indicates lipid droplets inside as well as outside *A. tremulum*

**Figure 3.**
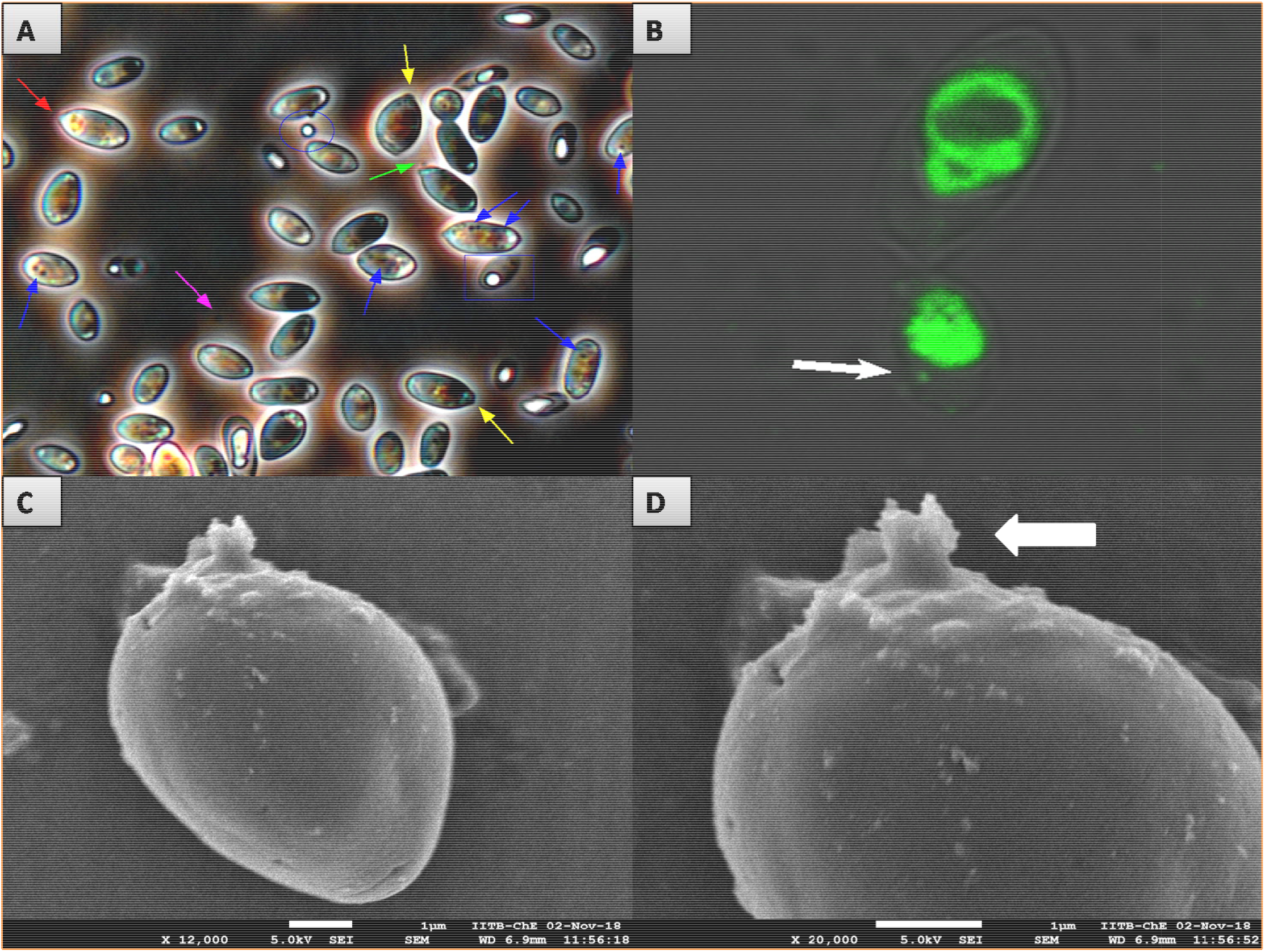
Mechanism of bacterial release from Yeast, **A.** Blue arrow indicates intracellular bacteria, magenta arrow indicates extracellular bacteria, green arrow indicates bacteria just released from yeast cell, yellow arrow indicates bacteria ready to release from yeast and initiation of protrusion formation in yeast, red arrow indicates protrusion of yeast after bacterial release, blue box indicates yeast cell containing lipid droplets and bacteria, blue circle indicates extracellular lipid droplets, **B.** White arrow indicates intracellular bacteria staining with auramine, **C.** Cryo-SEM showing yeast with protrusion at terminal, **D.** Magnified view of yeast protrusion shown by white arrow.

As mentioned above, it has all started with light microscopic observations only. We have been frequently and keenly observing the yeast using our light microscope. It was the dancing movement that makes us attracted to it. What we observed was, there were some bacteria surrounding yeast cells though it was a pure culture of yeast. To nullify the possibility of contamination, we have further streaked it on sterile PDA plates and reobtained the pure culture. After observing this reobtained pure culture under a microscope we have confirmed the bacterial presence surrounding yeast cells. For further confirmation, we have used phase-contrast microscopy, fluorescence microscopy^6–10^, confocal microscopy^11^, and electron microscopy (TEM) which confirms the coexistence of yeast and bacteria. In addition to bacterial cells, we have observed some lipid granules in and out of yeast cells (Fig. 2 and 3)^12–14^. Lipid granules were of varying sizes. In addition to that, since the bacterial and yeast population differ in size, we have performed Flow cytometry^15^. All these techniques gave evidence of the coexistence of yeast and bacteria^5,16^ and also about the presence of lipid granules^17–20^. The permanent and/or transient endosymbiotic relationship between yeast and bacteria is well reported^21–22^. From light microscopic observations, it becomes clear that after 4 days of incubation (in PDB) the color of yeast changes from colorless to dark brown.

### Movement of Yeast

Microscopic observation of *A.tremulum* was done by using a bright field microscope at 450x magnification. As mentioned above, there was a trembling movement of yeast. However, the displacement was also observed and has been mentioned in Fig. 1A and 1B and supplementary videos 1 and 2. It can be observed from the figure and videos that, some yeasts are trembling at their place, while some are displaced. This also nullifies any water/air current-born displacement. The total distance traveled with average velocity is 127.53 μm and 0.0265 μm/sec for figure 1A and 98.78 μm and 0.0577 μm/sec for Fig. 1B, respectively. The detailed displacement and velocity measurements are mentioned in supplementary data 1. Recently nanomotions of yeast have been reported by Willaert et al., (2020) ^3^. As per this report, the cellular activities of yeast are responsible for such nanomotions. However, the displacement observed in our studies is far more than as observed by Willaert et al., (2020) ^3^. We found that, the displacement and velocity of yeast decrease with time (Fig. 1). Such decreased displacement and velocity with time was also reported by Willaert et al., (2020) ^3^.

### Mechanism of Yeast Movement

As mentioned above, some yeast are trembling/twitching at their place while some are considerably displaced. This difference can be explained as follows. The dynamic movement of lipid droplets (Fig. 5 and supplementary video 6 and 7) and the movement of intracellular bacteria (supplementary video 2) are responsible for the trembling behavior of yeast. Whenever the intracellular bacteria has to move from the cytoplasm to the outside of a yeast cell, it has to cross the cell membrane. We observed that there is a thick exopolysaccharide (EPS) layer on the surface of the yeast cell (Fig. 4 A, B, and C). Besides, the cells are connected (tightly attached) to each other through this network (Fig. 4A). So, while moving from inside to the outside of a yeast cell, the endocellular bacteria has to cross the barrier of EPS. The struggle of bacteria to get released from the EPS layer leads to indirect movement/dragging of the yeast cell. But as mentioned, the yeast is attached to the EPS network, so the host yeast cannot move considerably by bacterial dragging (supplementary video 3a), hence tremble at a place. Some yeasts with less/weakened EPS network (Fig. 4 D) can be dragged away by bacteria and hence actually get displaced (supplementary video 3b).

**Figure 4.**
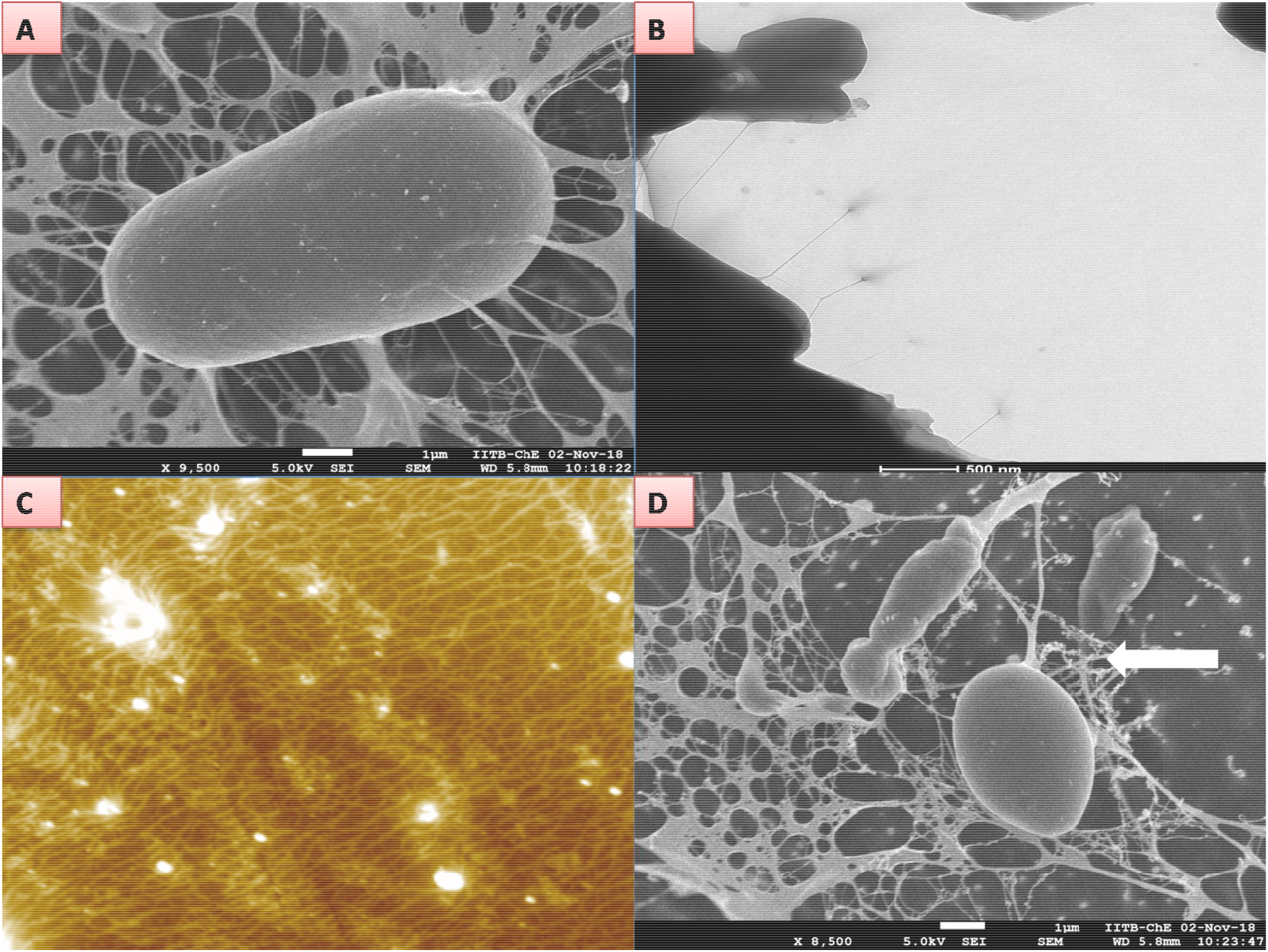
Exopolysaccharide of *A.tremulum*, A. Cryo-SEM image showing strong network of EPS, B. HR- TEM image showing EPS, C. AFM image showing EPS network, D. Cryo-SEM showing weakened network of EPS for a cell shown by white arrow.

**Figure 5.**
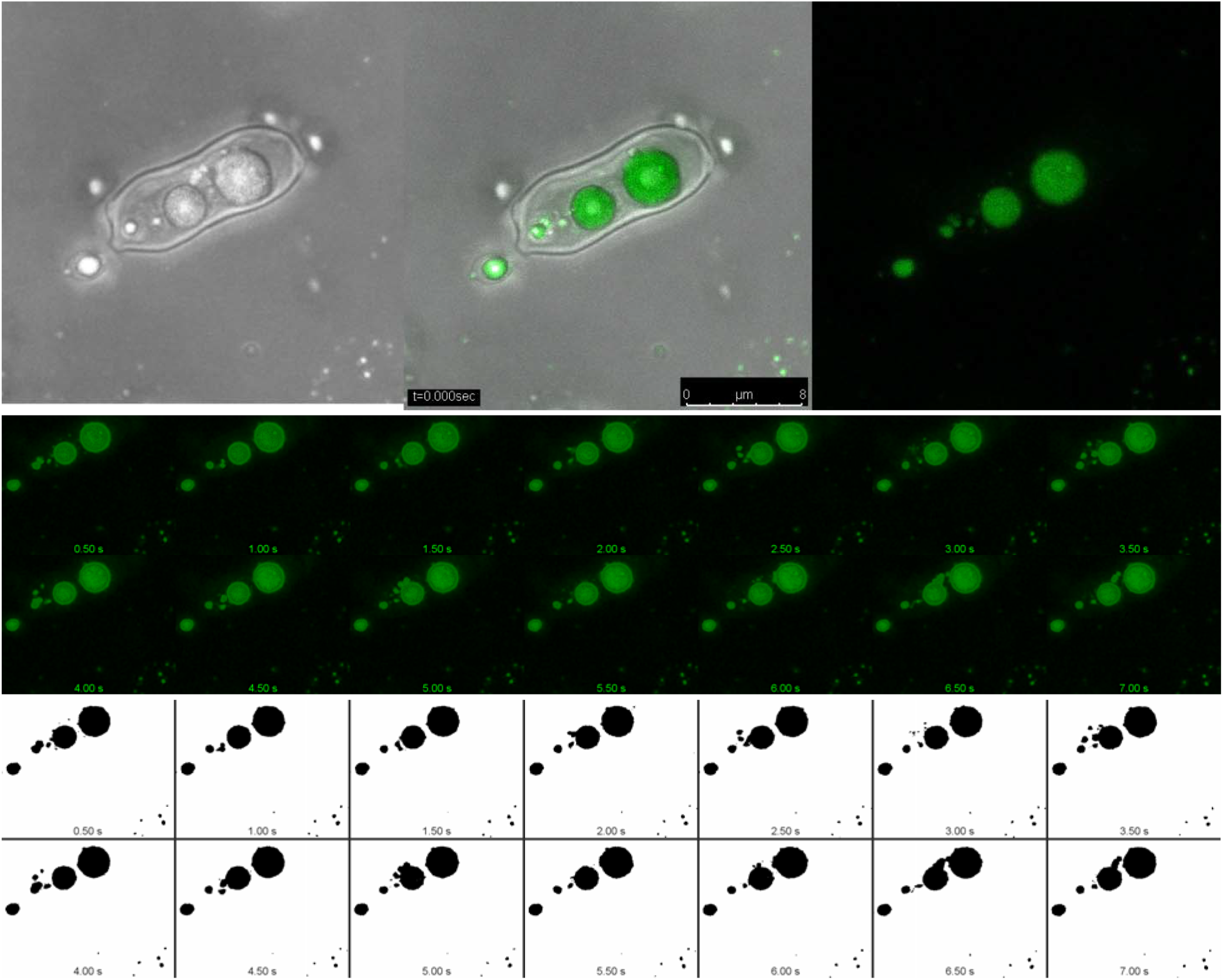
Movement within Yeast cell. Confocal analysis of yeast cell shows intracellular movement of lipid droplets and intra as well as extracellular movement of bacteria, Detailed movement can be observed in supplementary video 6

To confirm the bacteria-driven movement of yeast we have used Nutrient Agar grown yeast where lipid droplets are absent. We have observed yeast movement in these samples. In the case of yeast grown on PDA, the presence of lipid droplets increases the yeast movement for sure. However, the absence of lipid droplets does not affect basic yeast movement. The mere presence of lipid droplets is not responsible for yeast movement. This supports the bacteria-driven movement of yeast. A comparative video showing the movement of yeast in the presence and absence of lipid droplets has been mentioned as supplementary video 8. The details can be found in the respective captions of supplementary videos.

### Mechanism of Bacterial Release

Bacterial release from yeast can be observed in supplementary video 3b at 04:20 minutes frame. In the case of yeast with a thin exopolysaccharide layer on the surface, the bacterial release is easily facilitated. However, in the case of thick EPS on the yeast surface, the bacteria cannot get released easily. The bacterial release is facilitated by protrusion formation (Fig. 3A, C, and D. This protrusion can be observed in supplementary video 3b indicated by the black arrow at 4:19 minutes onwards. The formation of protrusion is due to the bacterial struggle for release.

### Bacterial Presence Vs Yeast Movement

It was observed that during the initial growth phase the bacterial number is quite less. The yeast movement was also less during this phase. The movement was found to be increasing with increasing ecto-endobacterial numbers. This suggests the importance of bacterial presence for yeast movement. Vigorously moving bacteria inside yeast cells can be observed in supplementary video 2 (shown as a red-colored box).

Extracellular bacteria have been tracked and mentioned in figure 5 (supplementary video 6). Three bacteria were labeled as A, B, and C. The detailed displacement (μm) and velocity (μm per sec) observed was 5.901 and 0.175 for bacteria A; 2.959 and 0.0699 for B; and 6.568, 0.198 for C, respectively (supplementary data 2).

The bacterial presence was also confirmed using FACS analysis (Supplementary Fig. 14 and 15).

### Lipid droplets in yeast Movement

Lipid droplets have been observed in *A.tremulum* when grown on PDA and MEA but not on NA. Lipid droplets were found inside as well as outside yeast cells. The intracellular dynamic movement of lipid droplets has been captured and is mentioned in Figure 5 and supplementary video 6. The distance traveled and velocity of the droplet have been determined. It was found that a single droplet has traveled a distance of 41.73 μm with an average velocity of 1.45 μm/sec. Detailed displacement and velocity measurements are mentioned in supplementary file 2. The dynamic movement of lipid droplets has been well-reviewed by Welte (2009)^23^. This movement of lipid droplets is facilitated by either microtubules or actin filaments. Martin and Parton (2006)^24^ have also tracked the movement of lipid droplets. The factors responsible for lipid movement could be nutrient transfer, biogenesis, and breakdown of lipid droplets and exchange with other cellular compartments^23^. When *A. tremulum* was grown on PDA and MEA, it was found that, movement of *A.tremulum* increases with an increasing number of lipid droplets. However, when grown on NA, *A.tremulum* shows movement but no lipid droplets. This proofs that, the movement of *A. tremulum* is because of the bacteria and not because of the lipid droplets.

### Confirmation of Bacterial Presence using Metagenomic Studies

The bacterial presence was confirmed by using metagenomic studies. We have analyzed the pure culture of yeast with whole-genome metagenomic analysis. This was performed at Medgenome Pvt. Ltd. Bangalore, India. The metagenomic analysis shows clear evidence of the presence of bacteria with *A.tremulum* (Supplementary data 3).

The whole-genome Metagenome sequencing generated around 3.42 GB of raw reads. The FASTQC of these reads shows the Q30 score is above 88.78 % and the average GC percentage is around 49%.

### Microbial Diversity

Metagenome analysis indicates that most of the reads (99.889 % mapped to the Eukaryota domain. Therefore it is evident that the Eukaryota kingdom is the most abundant and followed by the bacterial kingdom. Similarly, the analysis shows Phylum Ascomycota is the most abundant one followed by phylum proteobacteria. At the genus level, Genus *Aureobasidium* is the most abundant genus. In the case of prokaryotic diversity, the presence of the following bacteria has been detected, *Agrococcus*, *Microbacterium, Micrococcus luteus, Propionibacterium, Psuedonocardiaceae, Bradyrhizobium japonicum, Deinococcus, Bacteroides, Ralstonia, Enterobacteriaceae, Acinetobacter junii, Pseudomonas, Xanthomonadaceae* (Supplementary data 4).

### Whole Genome Analysis of *A. tremulum*

This study reports the first complete genome sequence of *A. tremulum*. The annotated genome size is 24.13Mbp in length and there are no gaps in the genome sequence which is successfully submitted to NCBI (Accession Number JACXRX000000000). The genome has been compared with known species of *Aureobasidium* species. *A. tremulum* genome is relatively smaller in size than other *Aureobasidium* species such as *A. subglaciale* (25.79 Mbp), *A. namibiae* (25.42 Mbp), *A. pullulans* (25.79 Mbp)^25^.

The GC content of *A. tremulum* is 48.6% which is higher than the *A. pullulans* (41.5%) but approximately similar to other Aureobasidium species such as *A. subglaciale* (48.11%), A*. namibiae* (49.8%)^25^.

*A. pullulans* isthe most closely related species among other *Aureobasidium* species. Since this is a first genome sequence of *A. tremulum* therefore we are considering this as a novel sequence.

### Isolation of Ecto-endo bacteria from *A.tremulum*

Since the bacterial presence was observed inside as well as outside the yeast during microscopic observations, it was decided to isolate these bacteria. It was hypothesized that the same bacteria are moving in and out of the yeast cell depending on the physiological conditions. This nature of bacteria has been well-reviewed by Leveau and Preston 2008^26^. Considering this, a simple approach to isolation of Ecto-endo bacteria was used. The plates of nutrient agar + Fluconazole (1.5mg/ml) were used for isolation. Experiments were repeated six times and bacteria have been isolated. Bacteria have been observed microscopically as well as by foldscope. Molecular characterization has been done at the National Center for Microbial Resource (NCMR) Pune. Isolated bacteria were identified by 16s rRNA gene sequencing and were found to belong to the genus *Micrococcus and Pseudomonas*. The obtained results (Data not shown) had to lead to the conclusion that, some of these bacteria could be a novel bacterial species.

### Bacterial Mycophagy in Dimorphic Yeast

To prove the bacteria-driven movement of yeast, it was decided to obtain bacteria-free yeast (which would be probably non-moving). To achieve this, four approaches were used namely, separation using centrifugal force, separation using low pH medium, separation using an antibiotic treatment. None of these methods succeeded in obtaining bacteria-free yeast. Rather it was observed that, when antibiotic containing solid medium was inoculated with yeast culture, no growth of yeast was obtained. This shows an integral/obligate requirement of bacterial presence for the survival of the yeast. As a part of obtaining bacteria-free yeast, a liquid medium with antibiotics was also used, this gave interesting results. The effect of antibiotics on the bacterial population and subsequently on yeast survival was demonstrated. This was experimented by putting the yeast culture in PDB containing ciprofloxacin (5mg/ml)^27^. The growth and movement of yeast as well as bacteria were observed in a timely manner. It was found that, during initial hours, yeast cell remains intact but there is a decrease in bacterial number surrounding yeast. After four days of incubation, the cell wall of yeast starts disappearing as evident with light microscopic observations. The yeast cells appear to be bursting. For confirmation, the sample was observed under a cryo scanning electron microscope (Cryo -SEM)^28^. The cryo-SEM analysis clearly shows the bursting of yeast cells and the release of numbers of bacteria (Fig. 6). As per the images, the bursting is from within. So there would be an obvious question that, how such bursting from within the yeast cell has occurred. This phenomenon can be hypothesized in the following manner.

**Figure 6.**
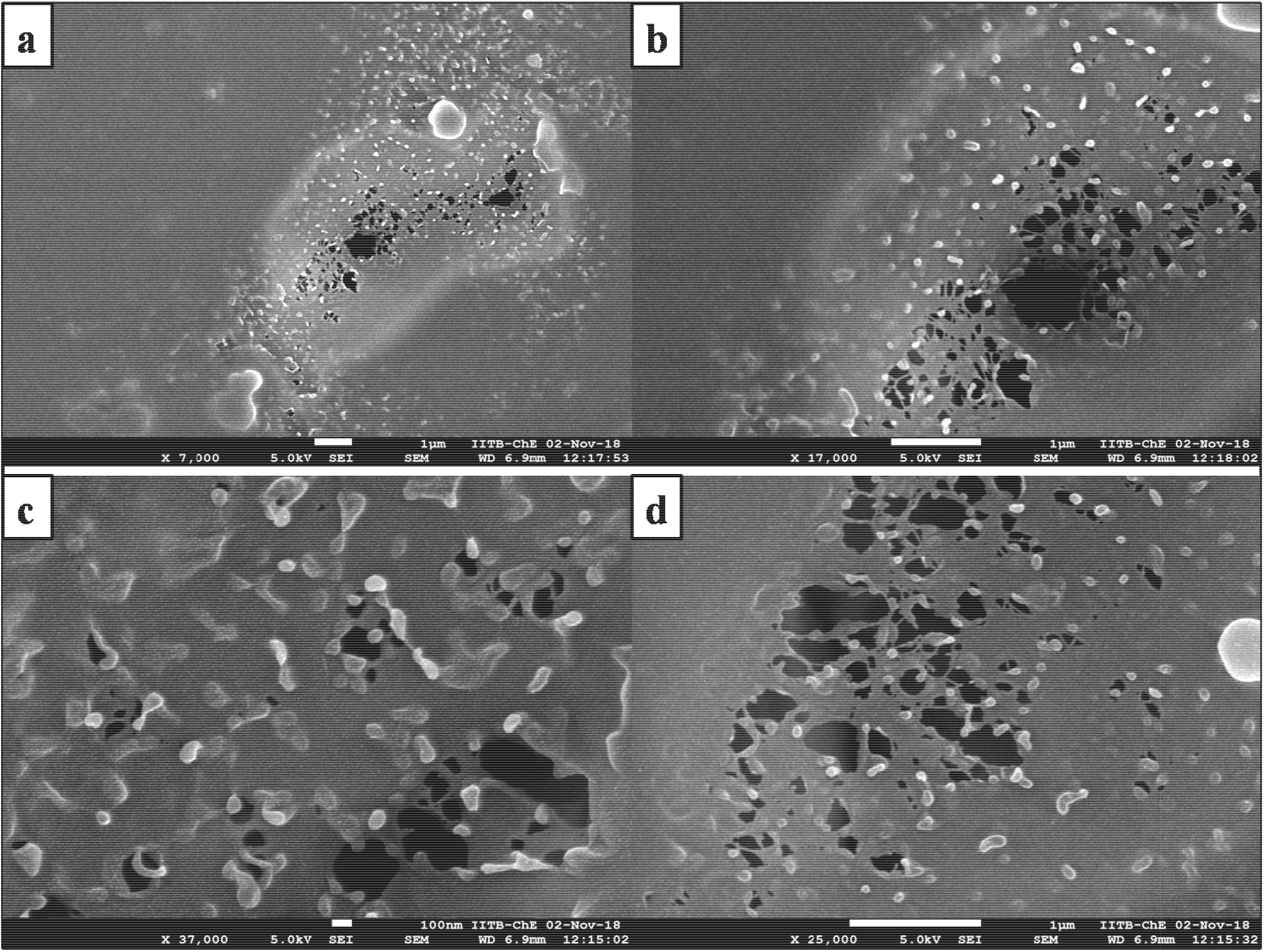
Cryo-SEM images showing Bursting of *A.tremulum* and release of endocellular bacterial population.

As we know, the yeast is carrying some bacteria with it even in pure culture. This means, somehow the yeast is availing nutrients to the bacteria for their survival. When growing under normal conditions, the bacteria can consume the nutrients present in the cytoplasm of yeast cells as well as the exudates secreted outside of yeast cells. This means the bacteria can freely move in and out of yeast cells (both intra and extracellular presence of bacteria have been located by microscopic observations) (Fig. 2, 3, 5, and 6) (supplementary videos 2-5). Now let us consider the drastic/unfavorable conditions (here presence of ciprofloxacin) in the outer environment of yeast. We have observed a drastic decrease in ectobacterial population in presence of ciprofloxacin. Here, the decrease in ectobacterial population can be either due to the killing of ectobacteria or due to the movement of ectobacteria to yeast cytoplasm. We believe in later reason. We hypothesize that in presence of ciprofloxacin ectobacteria move inside yeast cells to protect themselves. Here comes the question that, how they would be living inside yeast? Some type of interactions is well reported where fungal cells provide intracellular metabolites as food to endosymbionts. Interestingly, the bacteria not only survive but multiplies as well, inside yeast. The inner bacterial number increases in such a way that, it creates a turgor pressure on the cell membrane and finally bursts. In short, we hypothesize that the phenomenon involved is a change in the mode of bacterial mycophagy^26^. In presence of antibiotics, bacteria shift from extracellular biotrophy to endocellular biotrophy. However, with long-term exposure to the antibiotic, the bacterial shifting from endocellular biotrophy to extracellular biotrophy is restricted and results in multiplication and subsequent crowding of bacterial cells inside yeast. This finally led to increased internal turgor pressure on the cell membrane and eventually results in yeast cell bursting. This proposed phenomenon can be observed in cryo-SEM images (Figure 6) and the HR-TEM image (Figure 2c). Now as incubation time increases, the yeast cells had already busted and released the bacteria. The released bacteria get killed by the ciprofloxacin already present in the medium. This medium, if streaked on nutrient or potato dextrose agar, results in the growth of neither yeast nor bacteria. The bacterial mycophagy has never been reported for dimorphic yeast.

A question may come to our mind that, why not in normal (without antibiotic) conditions the bursting is observed. This can be answered as, in normal conditions, the bacterial cells can come out of yeast, so no bursting occurs. This means there is no restriction on extracellular biotrophy. So, both extra and endocellular biotrophy works in normal conditions.

### Nature of Bacteria: Obligate ecto/endo cellular or Facultative

The extracellular and endocellular bacterial presence was observed^5^ using light microscopic (supplementary video 2 and 3), phase contrast (Fig. 2A and 3A, Supplementary video 4), fluorescence microscopy (Figure 2B), confocal microscopy (Fig. 3B and 5 supplementary video 6), cryo-SEM (Fig. 6) and HR TEM (Fig. 2c). This indicates that bacteria can move in and out of yeast cells. This suggests the facultative nature of bacteria.

In the mycelial form of yeast, no significant presence of ectobacteria was observed as per light microscopic observations. In contrast, ectobacterial numbers sharply and rapidly increased in yeast form. The possible mechanism could be the release of the bacterial population from mycelia during the conversion of mycelial to yeast form.

The possibility of obligate extracellular coexistence of bacteria with yeast can be ruled out by obtaining a pure culture of yeast in repetitive experiments. Thus proving that the pure culture of yeast carries the bacteria which are present in yeast cytoplasm, suggesting the facultative nature of bacteria.

Bacteria were isolated from yeast. These bacteria were supposed to be ecto-endocellular bacteria of yeast.

### Bacteria- Yeast Association: Obligate or Facultative?

When yeast was inoculated on NA+ Fluconazole, the presence of bacteria was evident with well-isolated colony formation. On contrary, when NA+Antibiotics were used, no growth of yeast was evident.

5mg/mL concentration of Trimethoprim + sulphamethoxazole has been assessed for its effect on yeast and its endosymbionts. After incubation of 36 days, the culture medium was streak inoculated on nutrient and potato dextrose agar and resulted in the growth of yeast as well as bacteria. This means there is no effect of trimethoprim+ sulphamethoxazole. Yeast cell appears to grow normally with motile bacteria & lipid droplets.

Yeast cells were found to be susceptible to ampicillin in some manner. This is because the movement was found to be decreased and some cytosolic material appears to be coming out of the cell. Bacterial number not affected by ampicillin. Both bacteria and yeast have been observed to be unaffected by tetracycline (5mg/mL). Chloramphenicol (5mg/ml) is lowering bacterial number (no complete removal) but not necessarily affecting yeast growth.

As mentioned above, the endocellular bacteria from yeast were isolated and were found to be able to grow without yeast. However, when various combinations of antibiotics were used, the yeast was found to be unable to grow without bacteria as evident from the results of antibiotic susceptibility.

This indicates that bacteria are not dependant on yeast for survival but yeast is dependent on bacteria for survival. In short, the bacteria –yeast association is obligate from yeast’s point of view and is facultative from a bacterial point of view.

## Conclusion

Bacteria-driven movement of yeast has given a new perspective towards eukaryote-prokaryote association. Though the presence of endocellular/endohyphal bacteria has been known since the last decade, the phenomenon mentioned in the article is novel.

Endohyphal bacteria from dimorphic yeast have been microscopically observed. Such bacteria have been isolated and identified by molecular methods. The presence of bacteria has been proven by metagenomic studies and also by electron microscopic analysis. The experiments with antibiotics suggest that, though the bacteria can grow without yeast, yeast cannot grow without bacteria. This leads to showing the obligate symbiotic relationship of endosymbionts with yeast. Restriction on shifting of bacterial mycophagy in presence of antibiotics has been demonstrated.

This is the first report of the presence of endocellular bacteria in dimorphic yeast. There is a lot of scope for future explorations like the role of such endocellular bacteria in the conversion of mycelial to yeast form, role in the survival of yeast, and metabolite production. As reported earlier, many endosymbiotic bacteria from various macro and microorganisms are found to be responsible for anticancer agents which were previously considered to be secreted by their respective host in this context, we should not wonder if these endosymbionts would be found to be the actual producer of metabolites (polysaccharide, melanin) which are considered to be produced by *A.tremulum* itself.

The indifferent role of bacteria, in providing movement to its host has opened new doors of research.

## Materials and Methods

### Phase Contrast Microscopy

Phase-contrast microscopy is an optical microscopy technique that converts phase shifts in the light passing through a transparent specimen to brightness changes in the image. Phase shifts themselves are invisible but become visible when shown as brightness variations. One drop of distilled water with a loopful of yeast culture was taken on a clean slide and mix by using a sterile nichrome loop, this placed the coverslip onto yeast culture and was observed under phase-contrast microscopy at 40x (OLYMPUS BX53F V-DA).

### Fluorescence Microscopy

Is an optical microscope that uses fluorescence and phosphorescence instead of, or in addition to, scattering, reflection, and attenuation or absorption, to study the properties of organic and inorganic substances. 100μl distilled water with a loopful of yeast culture was taken on a clean slide and mix with 10μl of DAPI fluorescent dye and incubated in dark for 10 minutes. After incubation sample was observed under a fluorescence Microscope (OLYMPUS BX53F V-DA) at 40x.

### Confocal Microscopy

*A. tremulum* culture was stained with Auramine O and examined by using the Leica TCS SP 5 Confocal Laser Scanning Microscope (Leica Microsystems, Wetzlar, Germany) in the Laboratory of Bioimaging facility NCCS, Pune of Maharashtra province, India.

The LAS X version 3.4.2 183668 software (Leica Microsystems, Wetzlar, Germany) and ImageJ 1.53a version were used for further analysis. The fluorescence excitation of stained cells was induced using Argon visible (Leica Microsystems, Wetzlar, Germany) while the detection was recorded by PMT 2 (495nm-640nm) with 109 gain. Moreover, a PMT trans channel was used to visualize the cells in transmitted light with 354 gain. The sample was scanned in ‘xyzt’ mode to depth 7.41 μm while using HCX PL APO CS 100.00× 1.40 (Leica Microsystems, Wetzlar, Germany) Oil objective^11^.

### Cryo-Scanning Electron Microscopy (Cryo- FEG- SEM)

Cryo SEM was performed at IIT, Mumbai, India (Cryo-FEG-SEM JEOL JSM-7600F). Cryogenic Scanning electron microscopy is a form of electron microscopy where a hydrated but cryogenically fixed sample is imaged on a scanning electron microscope’s cold stage in a cryogenic chamber. The cooling is usually achieved with liquid nitrogen.

The sample was placed (200-500microliter) on carbon tape and freezing was carried out at −2000C under liquid nitrogen by slush freeze followed by sublimation at −85°C for 10 minutes. The sample was sputtered with platinum and imaging was performed at different resolutions.

### High Resolution-Transmission Electron Microscopy (HR-TEM)

HR-TEM was performed with FEI-Tecnai G^2^ 20 transmission electron microscope facility in IIT Delhi using a double tilt sample holder, 200-500 microliter sample suspension was placed on the carbon-coated copper grid by micropipette and examined.

### Flow Cytometry

To prove the coexistence of yeast and bacteria we have used flow cytometry. Flow cytometry is a technique used to detect and measure the physical and chemical characteristics of a population of cells or particles. Brown-colored pigmented yeast colony was scraped from potato dextrose agar plate and mix into 10 ml sterile saline (0.85%) and filtered using Whatman filter paper no 1. The filtrate was added into a clean small-size test tube and the probe of the flow cytometer was deep into this suspension.

### Preparation of bacteria-free yeast

To understand the interactions between yeast and bacteria it was essential to know whether yeast can survive in absence of bacteria or not. To assess this, four approaches were used.

### Separation of yeast from bacteria using centrifugation

Enrichment of Yeast culture: 1mL of yeast culture was inoculated in sterile 100 ml of potato dextrose broth & incubated at 25°C for 2 days. This results in the enrichment of yeast culture.

Separation of Yeast and Bacteria by applied centrifugal force: 2ml of enriched yeast culture was aseptically collected and centrifuged at different rpm such as 4000, 6000, 8000, 10,000, 12,000, 14,000, 16,000, 18,000, 20,000 for constant time i.e. 5 minutes & supernatant was collected in another sterile tube separately.

Analysis of supernatant and pellet for the presence of bacteria and yeast respectively - Collected supernatant from the respective tube was streaked on sterile Nutrient agar plates separately & incubated at 25°C for 2 days.

### Separation of yeast from bacteria by Lowering the pH of culture medium

100 ml of potato dextrose agar having pH 3.0 was prepared. The sterile medium was poured into sterile Petri plates. After solidification active yeast culture was streaked on plates containing agar medium plates and then incubated at 25ºC for 6 days.

### Separation of yeast from bacteria using Antibiotics (Agar ditch method)

100 ml of potato dextrose agar was prepared and poured into sterile Petri plates. After solidification, two parallel lines were drawn by marking a ditch at one end of the Petri dish. On the Marked portion, some portion of agar was removed using a sterile nichrome wire loop to make a ditch in agar. Yeast culture was streaked perpendicular to the ditch at equidistance. The ditch was filled with the help of a sterile pipette with antibiotic solution ciprofloxacin, trimethoprim, and sulphamethoxazole 5mg/ml separately. Plates were then incubated at 25ºC for 2 days.

### Assessing the susceptibility of yeast to varying concentrations of antibiotics in solid medium

Ciprofloxacin, ampicillin, Tetracycline & Trimethoprim + Sulphamethaxazole solutions were prepared to have a concentration of 5mg/mL. Sterile filter paper discs (diameter 4 mm) were dipped in each of above mentioned antibiotic solutions and placed on PDA plates previously inoculated with yeast culture. The plates were then incubated at 25ºC.

### Assessing the effect of antibiotic concentration on the growth of yeast in solid medium

Ciprofloxacin & Chloramphenicol antibiotic solution were prepared with concentrations (mg/ml) 5, 10, 15, 20 and 25. Sterile Filter paper discs (diameter 4 mm) were dipped in each of above mentioned antibiotic solutions and placed on PDA plates previously inoculated with yeast culture. The plates were then incubated at 25ºC.

### Separation of yeast from bacteria using Antibiotics / Assessment of susceptibility of yeast to the antibiotics in liquid culture

#### Ciprofloxacin

1mL of yeast culture was inoculated in 100ml of sterile potatoes dextrose broth & incubated at 25°C for 24 hours. After incubation ciprofloxacin (Cipodac 500mg) as per concentration 5mg/ml was added into overnight grown yeast culture & again incubated at 25°C. After and during incubation of 4 days the culture was observed in wet mount preparations under a light microscope at 450X magnification. This culture was also inoculated on sterile plates of nutrient and potato dextrose agar for the possible growth of yeast.

#### Trimethoprim + Sulphamethaxazole

1mL of yeast culture was inoculated in 100mL of sterile potatoes dextrose broth & incubated at 25°C for 24 hours. After incubation, Trimethoprim + Sulphamethaxazole (Cotrimoxazole 500mg) as per concentration 5mg/mL was added into overnight grown yeast culture & again incubated at 25°C. After and during incubation of 4 days the culture was observed in wet mount preparations under a light microscope at 450X magnification. This culture was also inoculated on sterile plates of nutrient and potato dextrose agar for the possible growth of yeast.

#### Ampicillin

1mL of yeast culture was inoculated in 100mL of sterile potatoes dextrose broth & incubated at 25°C for 24 hours. After incubation Ampicillin (Bacipen 500mg) as per concentration 5mg/ml was added into overnight grown yeast culture & again incubated at 25°C. After and during incubation of 4 days the culture was observed in wet mount preparations under a light microscope at 450X magnification. This culture was also inoculated on sterile plates of nutrient and potato dextrose agar for the possible growth of yeast.

#### Oxytetracycline hydrochloride

1mL of yeast culture was inoculated in 100ml of sterile potatoes dextrose broth & incubated at 25°C for 24 hours. After incubation, Oxytetracycline hydrochloride (TERRAMYCIN 500mg) as per concentration 5mg/mL was added into overnight grown yeast culture & again incubated at 25°C. After and during incubation of 4 days the culture was observed in wet mount preparations under a light microscope at 450X magnification. This culture was also inoculated on sterile plates of nutrient and potato dextrose agar for the possible growth of yeast.

#### Chloramphenicol

1mL of yeast culture was inoculated in 100mL of sterile potatoes dextrose broth & incubated at 25°C for 24 hours. After incubation chloramphenicol (5mg/ml) was added into overnight grown yeast culture & again incubated at 25°C for 7 days. After 7 incubation 10 mL of chloramphenicol (5mg/mL) containing yeast culture was added into fresh sterile 100mL potatoes dextrose broth& incubated at 25ºC for 16 days. After 16 days of incubation chloramphenicol containing yeast culture was added into a sterile test tube & chloramphenicol as per concentration 5mg/mL was added into the same test tube & incubated at 25ºC for 24 hours. After and during incubation of 4 days the culture was observed in wet mount preparations under a light microscope at 450X magnification. This culture was also inoculated on sterile plates of nutrient and potato dextrose agar for the possible growth of yeast.

### DNA extraction, Library Preparation & Whole Genome Metagenome sequencing

High molecular weight DNA was extracted from yeast culture/ mixed culture using Qubit BR DNA kit (Invitrogen) genomic DNA kit following the manufacturer’s instructions at geneOmbio technologies Pvt ltd. company Pune. DNA quantification and size assessment were conducted using geneOmbio technologies Pvt ltd. company Pune.

To prove the bacterial presence with yeast, we have used the whole-genome Metagenome approach to sequence the complete fungal genome and to identify the Microbiome associated with it. The genomic DNA was fragmented to 350 base pairs using Covaris M220 Focused ultrasonicator. Following manufacturer instructions, pair-end libraries were obtained using NEBNext Ultra DNA Library Preparation Kit. The Quality and quantity of libraries were checked using a tape station (Agilent Technologies, Santa Clara, California, USA) before sequencing them on the Illumina HiSeqX platform (Illumina Inc., San Diego, California, USA) with 2X150 bp sequencing chemistry at Medgenome Pvt. Ltd. Bangalore.

### Metagenome Analysis

The basic analysis includes quality checks and adapter removal of the raw Fastq files. This is followed by the removal of any human DNA contamination by aligning the reads to the human reference genome (hg19/GRCh37) using BWA MEM aligner and only the unaligned reads were taken for further processing^29^. We also aligned the reads to the bacterial (11400 genomes), fungal (280 genomes), viral genome (7854) databases using BWA MEM aligner^29^. Further, the de novo assembly was carried out using metaSPAdes assembler (v3.11.1)^30^. Assembly is performed with k-mer size 55 using the de-Bruijn graph method. The assembled scaffolds were used for gene prediction and used as input to prodigal (v2.6.3) for the prediction of open reading frames (ORFs)^31^. The obtained ORFs are filtered using in-house Perl scripts. ORFs of length below 100 bp is being filtered. Supplementary data 3 depicts the ORF length distribution. The predicted ORFs are searched against the non-redundant NCBI-Nr protein database [O’Leary et al., 2016] using DIAMOND (v0.7.9.58)^32^.

The alignment file along with the filtered ORFs are used as input to the functional annotation using MEGAN5 (MEtaGenome ANalyzer) software^33^. Since the abundance was only 0-0.1% therefore, annotations are not obtained for viruses and Archae. The species with a relative abundance of more than 1%are taken for further analysis.

### Denovo genome assembly and annotation of *A. tremulum*

Denovo assembly was built from the obtained whole-genome metagenome reads using publically available MetaWRAP^34^ and MEGAHIT^35^ pipelines. Subsequently, to achieve the *A. tremulum genome* sequence the obtained reads were mapped back to the reference genome *A.pullulans* (NW_021941021.1) using the BWA pipeline^29^. For genome annotation, we used GenSAS in which mainly Augustus, PASA, and Diamond were used^36^. However, we used an in-house script and BLASTn for the classification of the novel or unannotated genome sequences^37^ The *A tremulum* genome was submitted to the NCBI using the genome submission portal.

### Isolation and Identification of Ecto-endobacteria from yeast

To have supportive proof we have decided to isolate bacteria from this ‘Pure Culture’ of yeast. 100 ml of Nutrient agar was prepared and after autoclaving, it was then added with fluconazole to get a final concentration of 1.5mg/mL. The obtained sterile medium was then poured into sterile Petri dishes. After solidification, the pure culture of yeast was streak inoculated on these plates and then incubated at room temperature for 4 days. The experiment was done in 5 sets and repeated for 4 times. The obtained isolates were then identified using 16s rRNA gene sequencing.

## Supporting information

Supplementary Videos with captions

## Acknowledgment

Part of the present research work has been done using funds received from the Department of Biotechnology, Government of India for the project entitled “Foldscope as a research Tool” sanctioned to Dr. Adhapure Nitin N. during 2018-19 (No. BT/IN/Indo-US/Foldscope/39/2015). We are thankful to Dr. Rohit Sharma, Scientist ‘C’, National Center for Microbial Resource, Pune, for useful discussions.

## Author Contributions

GP and SC done basic laboratory work, MK did antibiotic-related experimentation, AI and VS did image analysis and referencing part, CJ and LT did the bioinformatics part, and NA was responsible for ideation, designing of experiments, initial laboratory work, and manuscript writing.

## References

1. Flor-Parra, I., Bernal, M., Zhurinsky, J., & Daga, R. R.. Cell migration and division in amoeboid-like fission yeast. Biology open, 3(1), 108–115. https://doi.org/10.1242/bio.20136783 (2014)

2. Crous, P. W., Carnegie, A. J., Wingfield, M. J., Sharma, R., Mughini, G., Noordeloos, M. E., Santini, A., Shouche, Y. S., Bezerra, J., Dima, B., Guarnaccia, V., Imrefi, I., Jurjević, Ž., Knapp, D. G., Kovács, G. M., Magistà, D., Perrone, G., Rämä, T., Rebriev, Y. A., Shivas, R. G., … Groenewald, J. Z.. Fungal Planet description sheets: 868-950. Persoonia, 42, 291–473. https://doi.org/10.3767/persoonia.2019.42.11\ (2019)

3. Willaert, R. G., Vanden Boer, P., Malovichko, A., Alioscha-Perez, M., Radotić, K., Bartolić, D., Kalauzi, A., Villalba, M. I., Sanglard, D., Dietler, G., Sahli, H., & Kasas, S.. Single yeast cell nanomotions correlate with cellular activity. Science advances, 6(26), eaba3139. https://doi.org/10.1126/sciadv.aba3139 (2020)

4. Ben Tahar, I., Kus-Liśkiewicz, M., Lara, Y., Javaux, E., & Fickers, P.. Characterization of a nontoxic pyomelanin pigment produced by the yeast Yarrowia lipolytica. Biotechnology progress, 36(2), e2912. https://doi.org/10.1002/btpr.2912 (2020)

5. Heydari, S., Siavoshi, F., Ebrahimi, H., Sarrafnejad, A., & Sharifi, A. H.. Excision of endosymbiotic bacteria from yeast under aging and starvation stresses. Infection, genetics and evolution : journal of molecular epidemiology and evolutionary genetics in infectious diseases, 78, 104141. https://doi.org/10.1016/j.meegid.2019.104141 (2020)

6. de Jonge N, Peckys DB. Live Cell Electron Microscopy Is Probably Impossible. ACS Nano.;10(10):9061–9063. doi:10.1021/acsnano.6b02809 (2016)

7. Salmanian, A. H., Siavoshi, F., Akbari, F., Afshari, A., & Malekzadeh, R.. Yeast of the oral cavity is the reservoir of Heliobacter pylori. Journal of oral pathology & medicine : official publication of the International Association of Oral Pathologists and the American Academy of Oral Pathology, 37(6), 324–328. https://doi.org/10.1111/j.1600-0714.2007.00632.x (2008).

8. Salmanian, A.H., Siavoshi, F., Beyrami, Z., LatifiLNavid, S., Tavakolian, A. and Sadjadi, A., Foodborne yeasts serve as reservoirs of *Helicobacter pylori*. Journal of Food Safety, 32: 152–160. doi:10.1111/j.1745-4565.2011.00362.x (2012)

9. Siavoshi F, Nourali-Ahari F, Zeinali S, Hashemi-Dogaheh M, Malekzadeh R, Massarrat S. Yeasts protects Helicobacter pylori against the enviromental stress. Arc Iran Med. ;1:2–8. (1998)

10. Siavoshi, F., Salmanian, A. H., Akbari, F., Malekzadeh, R., & Massarrat, S.. Detection of Helicobacter pylori-specific genes in the oral yeast. Helicobacter, 10(4), 318–322. https://doi.org/10.1111/j.1523-5378.2005.00319.x (2005)

11. Rajkowska, K., Nowicka-Krawczyk, P., & Kunicka-Styczyńska, A.. Effect of Clove and Thyme Essential Oils on *Candida* Biofilm Formation and the Oil Distribution in Yeast Cells. Molecules (Basel, Switzerland), 24(10), 1954. https://doi.org/10.3390/molecules24101954 (2019)

12. Thoms, S., Debelyy, M. O., Connerth, M., Daum, G., & Erdmann, R.. The putative Saccharomyces cerevisiae hydrolase Ldh1p is localized to lipid droplets. Eukaryotic cell, 10(6), 770–775. https://doi.org/10.1128/EC.05038-11 (2011)

13. Adeyo, O., Horn, P. J., Lee, S., Binns, D. D., Chandrahas, A., Chapman, K. D., & Goodman, J. M.. The yeast lipin orthologue Pah1p is important for biogenesis of lipid droplets. The Journal of cell biology, 192(6), 1043–1055. https://doi.org/10.1083/jcb.201010111 (2011)

14. Wolinski, H., Kolb, D., Hermann, S., Koning, R. I., & Kohlwein, S. D.. A role for seipin in lipid droplet dynamics and inheritance in yeast. Journal of cell science, 124(Pt 22), 3894–3904. https://doi.org/10.1242/jcs.091454 (2011)

15. Rosebrock AP. Analysis of the Budding Yeast Cell Cycle by Flow Cytometry. Cold Spring Harb Protoc. 2017(1):10.1101/pdb.prot088740. Published Jan 3. doi:10.1101/pdb.prot088740 (2017)

16. Haiko, J., Saeedi, B., Bagger, G., Karpati, F., & Özenci, V.. Coexistence of Candida species and bacteria in patients with cystic fibrosis. European journal of clinical microbiology & infectious diseases : official publication of the European Society of Clinical Microbiology, 38(6), 1071–1077. https://doi.org/10.1007/s10096-019-03493-3 (2019)

17. van Zutphen, T., Todde, V., de Boer, R., Kreim, M., Hofbauer, H. F., Wolinski, H., Veenhuis, M., van der Klei, I. J., & Kohlwein, S. D.. Lipid droplet autophagy in the yeast Saccharomyces cerevisiae. Molecular biology of the cell, 25(2), 290–301. https://doi.org/10.1091/mbc.E13-08-0448 (2014)

18. Welte, M. A., & Gould, A. P.. Lipid droplet functions beyond energy storage. Biochimica et biophysica acta. Molecular and cell biology of lipids, 1862(10 Pt B), 1260–1272. https://doi.org/10.1016/j.bbalip.2017.07.006 (2017)

19. Graef M. (2018). Lipid droplet-mediated lipid and protein homeostasis in budding yeast. FEBS letters, 592(8), 1291–1303. https://doi.org/10.1002/1873-3468.12996

20. Vevea, J. D., Garcia, E. J., Chan, R. B., Zhou, B., Schultz, M., Di Paolo, G., McCaffery, J. M., & Pon, L. A.. Role for Lipid Droplet Biogenesis and Microlipophagy in Adaptation to Lipid Imbalance in Yeast. Developmental cell, 35(5), 584–599. https://doi.org/10.1016/j.devcel.2015.11.010 (2015)

21. McFall-Ngai M.. Are biologists in ‘future shock’? Symbiosis integrates biology across domains. Nature reviews. Microbiology, 6(10), 789–792. https://doi.org/10.1038/nrmicro1982 (2008)

22. Ruiz-Lozano, J. M., & Bonfante, P.. Identification of a putative P-transporter operon in the genome of a Burkholderia strain living inside the arbuscular mycorrhizal fungus Gigaspora margarita. Journal of bacteriology, 181(13), 4106–4109. https://doi.org/10.1128/JB.181.13.4106-4109.1999 (1999)

23. Welte M. A.. Fat on the move: intracellular motion of lipid droplets. Biochemical Society transactions, 37(Pt 5), 991–996. https://doi.org/10.1042/BST0370991 (2009)

24. Martin S, Parton RG. Lipid droplets: a unified view of a dynamic organelle. Nat Rev Mol Cell Biol.7(5):373–8. doi: 10.1038/nrm1912. PMID: 16550215. 2006

25. Chan, G. F., Bamadhaj, H. M., Gan, H. M., & Rashid, N. A.. Genome sequence of *Aureobasidium pullulans* AY4, an emerging opportunistic fungal pathogen with diverse biotechnological potential. Eukaryotic cell, 11(11), 1419–1420. https://doi.org/10.1128/EC.00245-12. (2012)

26. Leveau, J. H., & Preston, G. M.. Bacterial mycophagy: definition and diagnosis of a unique bacterial-fungal interaction. The New phytologist, 177(4), 859–876. https://doi.org/10.1111/j.1469-8137.2007.02325.x (2008)

27. Trenschel, R., Peceny, R., Runde, V., Elmaagacli, A., Dermoumi, H., Heintschel von Heinegg, E., Müller, K. D., Schaefer, U. W., & Beelen, D. W.. Fungal colonization and invasive fungal infections following allogeneic BMT using metronidazole, ciprofloxacin and fluconazole or ciprofloxacin and fluconazole as intestinal decontamination. Bone marrow transplantation, 26(9), 993–997. https://doi.org/10.1038/sj.bmt.1702655 (2000)

28. Szotkowski, M., Byrtusova, D., Haronikova, A., Vysoka, M., Rapta, M., Shapaval, V., & Marova, I.. Study of Metabolic Adaptation of Red Yeasts to Waste Animal Fat Substrate. Microorganisms, 7(11), 578. https://doi.org/10.3390/microorganisms7110578 (2019)

29. Li, H., & Durbin, R.. Fast and accurate short read alignment with Burrows-Wheeler transform. Bioinformatics (Oxford, England), 25(14), 1754–1760. https://doi.org/10.1093/bioinformatics/btp324 (2009)

30. Nurk, S., Meleshko, D., Korobeynikov, A., & Pevzner, P. A.. metaSPAdes: a new versatile metagenomic assembler. Genome research, 27(5), 824–834. https://doi.org/10.1101/gr.213959.116 (2017)

31. Hyatt, D., Chen, G. L., Locascio, P. F., Land, M. L., Larimer, F. W., & Hauser, L. J.. Prodigal: prokaryotic gene recognition and translation initiation site identification. BMC bioinformatics, 11, 119. https://doi.org/10.1186/1471-2105-11-119 (2010)

32. Buchfink, B., Chao Xie & Daniel H. Huson, Fast and Sensitive Protein Alignment usingDIAMOND, Nature Methods, 12, 59–60 doi:10.1038/nmeth.3176. (2015)

33. Huson, D. H., Auch, A. F., Qi, J., & Schuster, S. C.. MEGAN analysis of metagenomic data. Genome research, 17(3), 377–386. https://doi.org/10.1101/gr.5969107 (2007)

34. Uritskiy, G.V., DiRuggiero, J. & Taylor, J. MetaWRAP—a flexible pipeline for genome-resolved metagenomic data analysis. Microbiome 6, 158. https://doi.org/10.1186/s40168-018-0541-1 (2018)

35. Li D, Liu CM, Luo R, Sadakane K, Lam TW. MEGAHIT: an ultra-fast single-node solution for large and complex metagenomics assembly via succinct de Bruijn graph. Bioinformatics.;31(10):1674–1676. doi:10.1093/bioinformatics/btv033 (2015)

36. Humann, L; T Lee, S Ficklin, D Main. “Structural and Functional Annotation of Eukaryotic Genomes with GenSAS.” Gene Prediction: Methods and Protocols. Ed. Martin Kollmar. New York: Humana Press,. 29–51. DOI: 10.1007/978-1-4939-9173-0_3 (2019)

37. Altschul SF, Gish W, Miller W, Myers EW, Lipman DJ. Basic local alignment search tool. J Mol Biol.; 215(3):403–410. doi:10.1016/S0022-2836(05)80360-2 (1990)

