## Supplementary Figures for "Moving Yeasts: Resolving the Mystery"

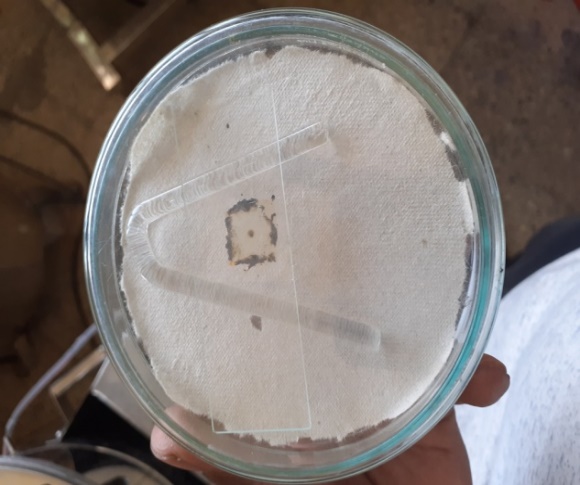

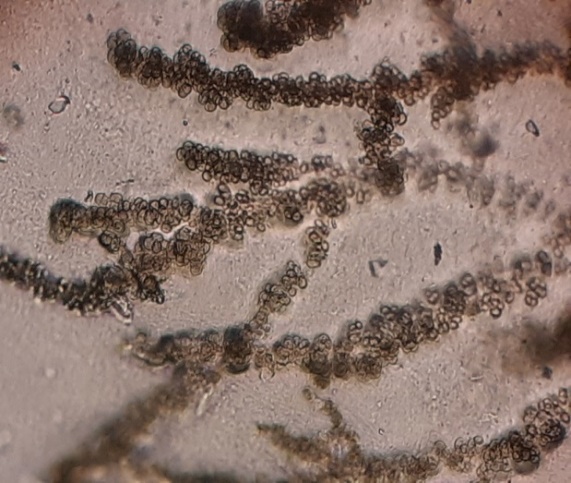

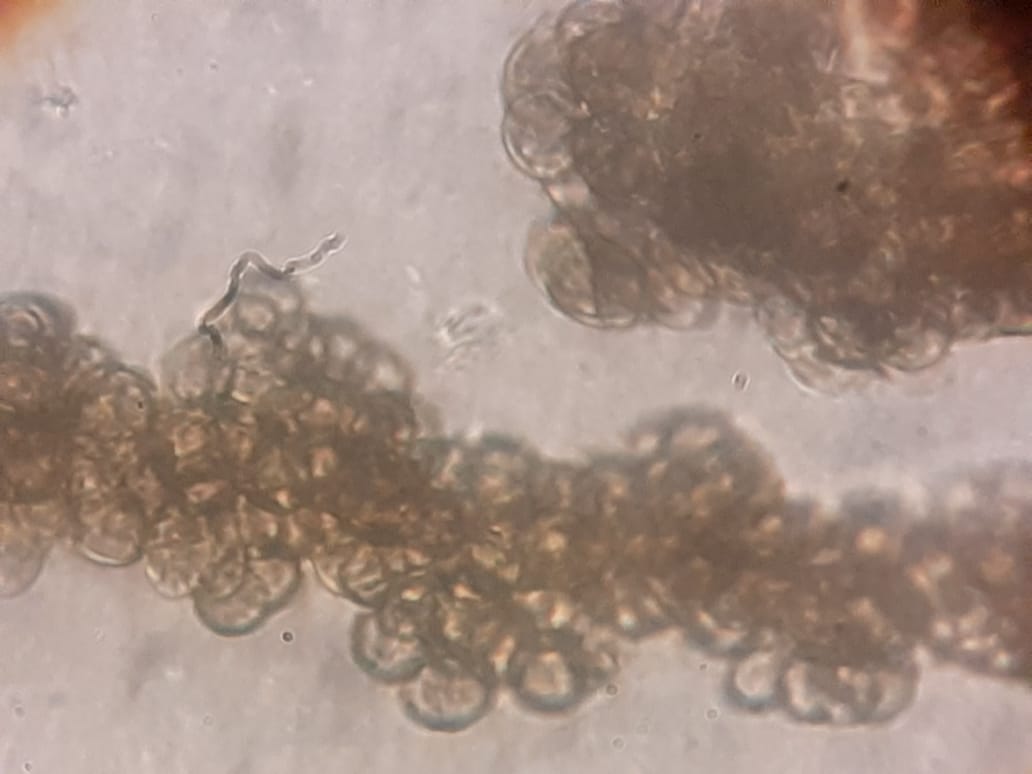

**A	B	C**

**Figure. 1 A:** Slide culture technique after ten days incubation, **B:** Microscopic image at 10x (A). **C:** Microscopic image at 45x (A).

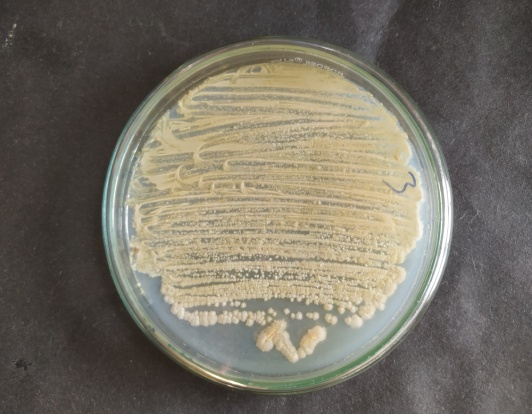

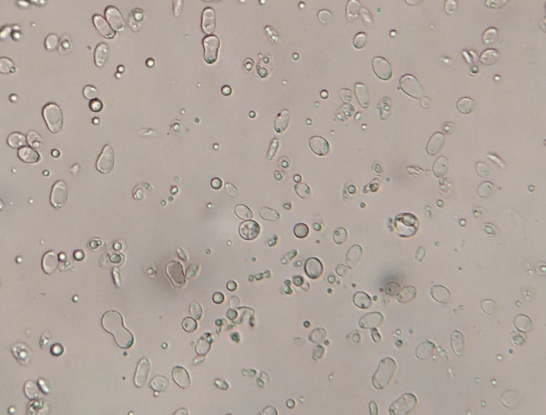

**A	B**

**Figure. 2 A.** Growth of A.tremulum on PDA after 6 days of incubation at pH 3.0. **B.** Microscopic view of yeast at 450x.

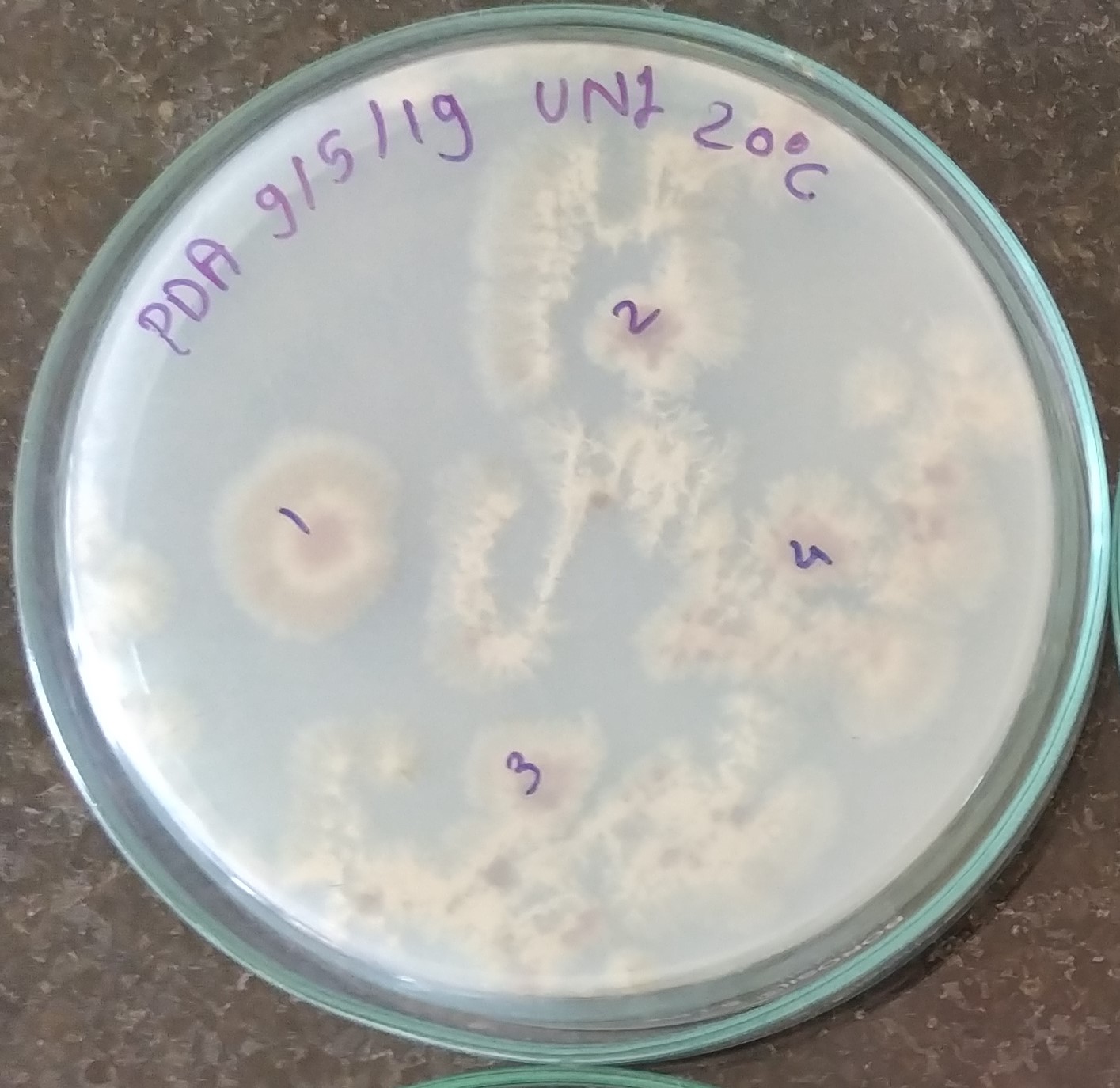

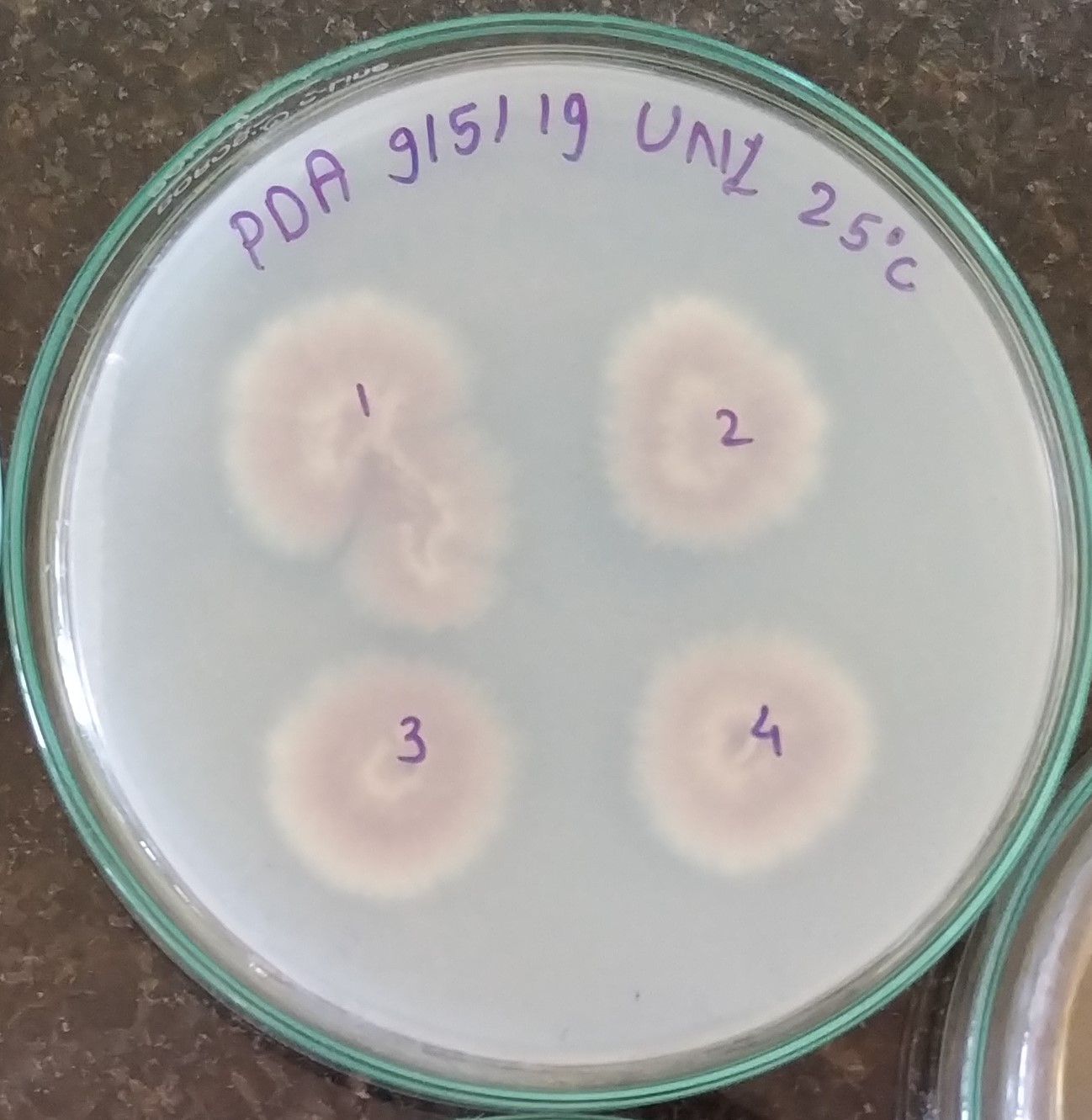

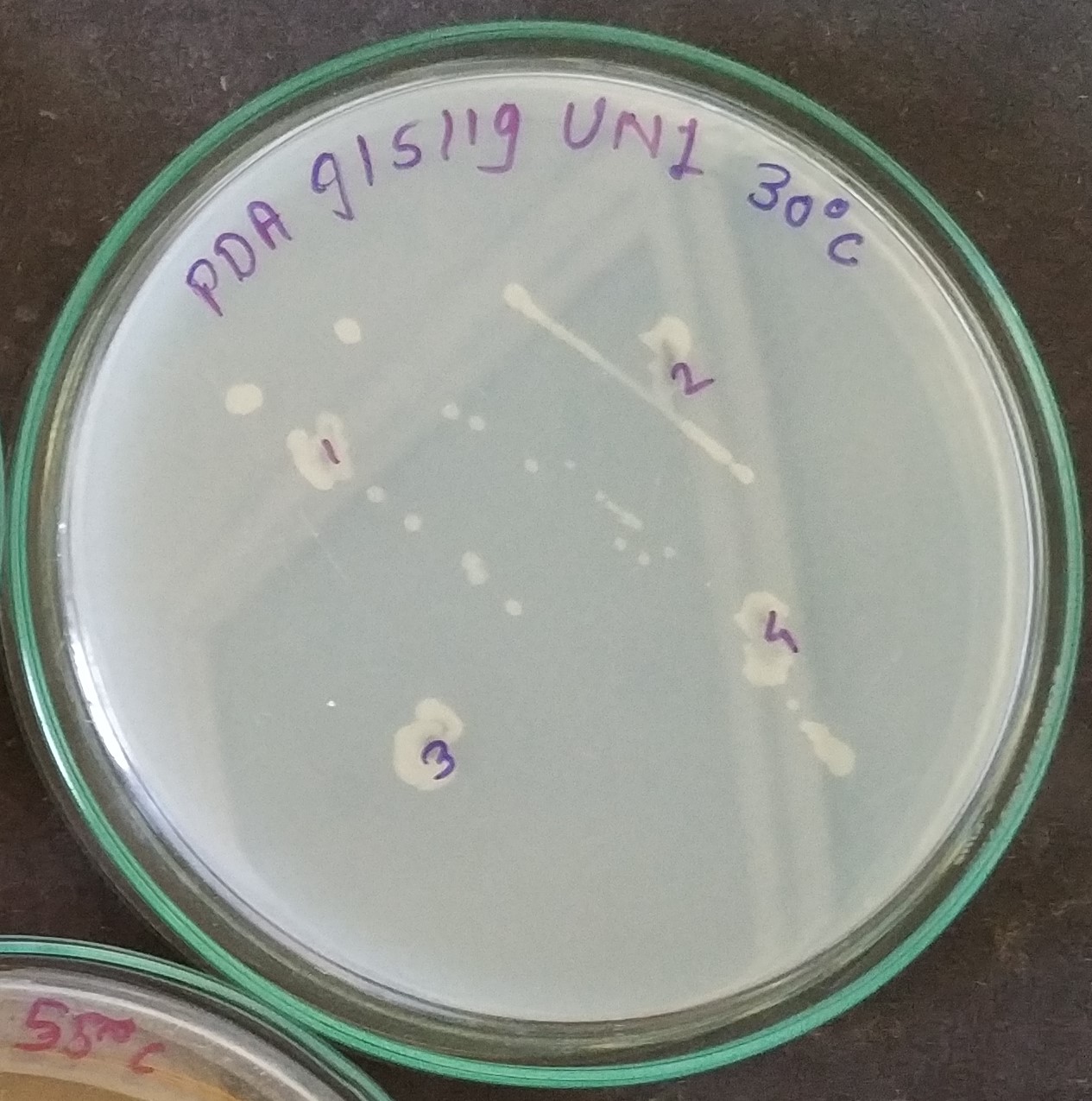

**A	B	C**

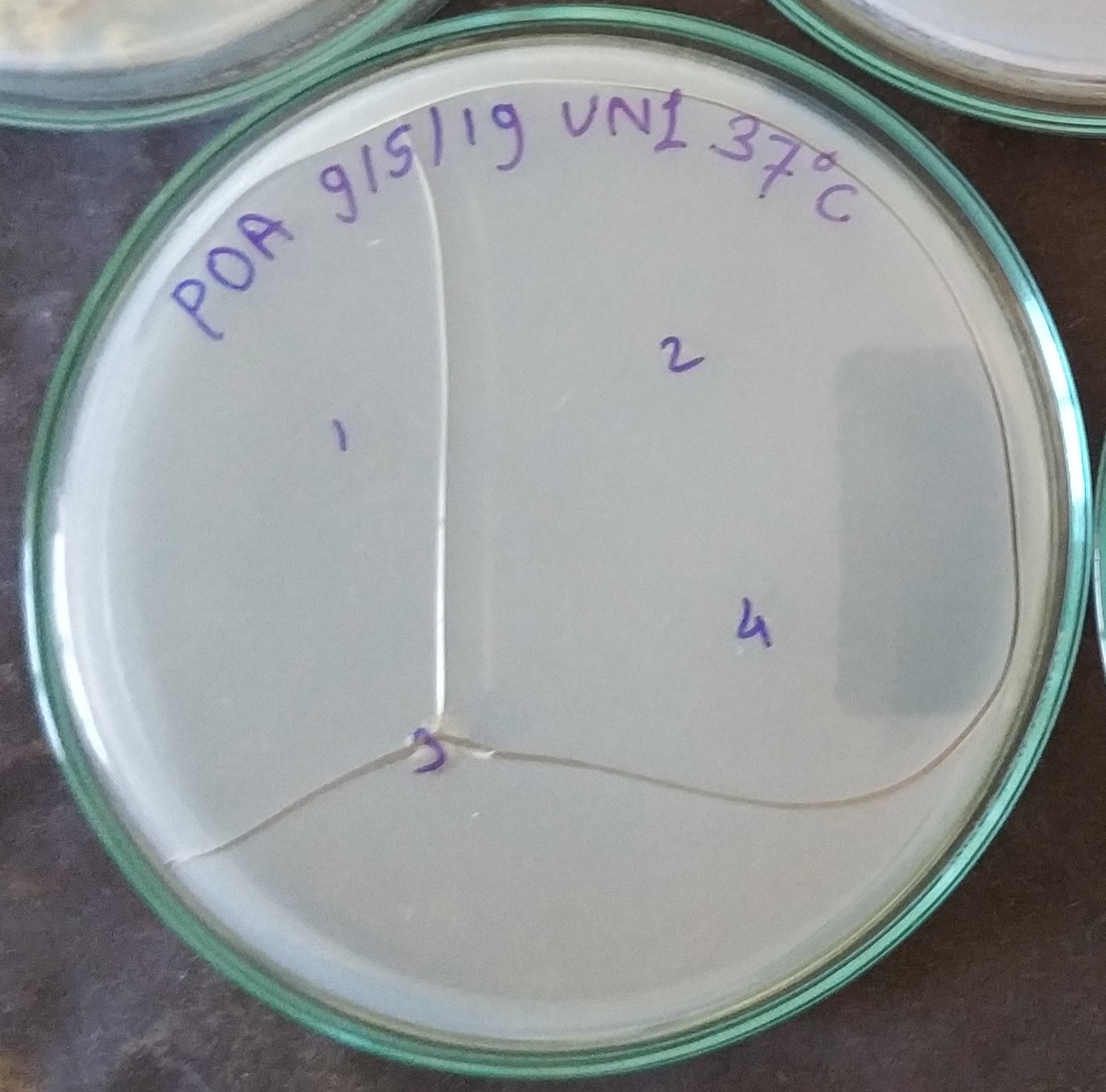

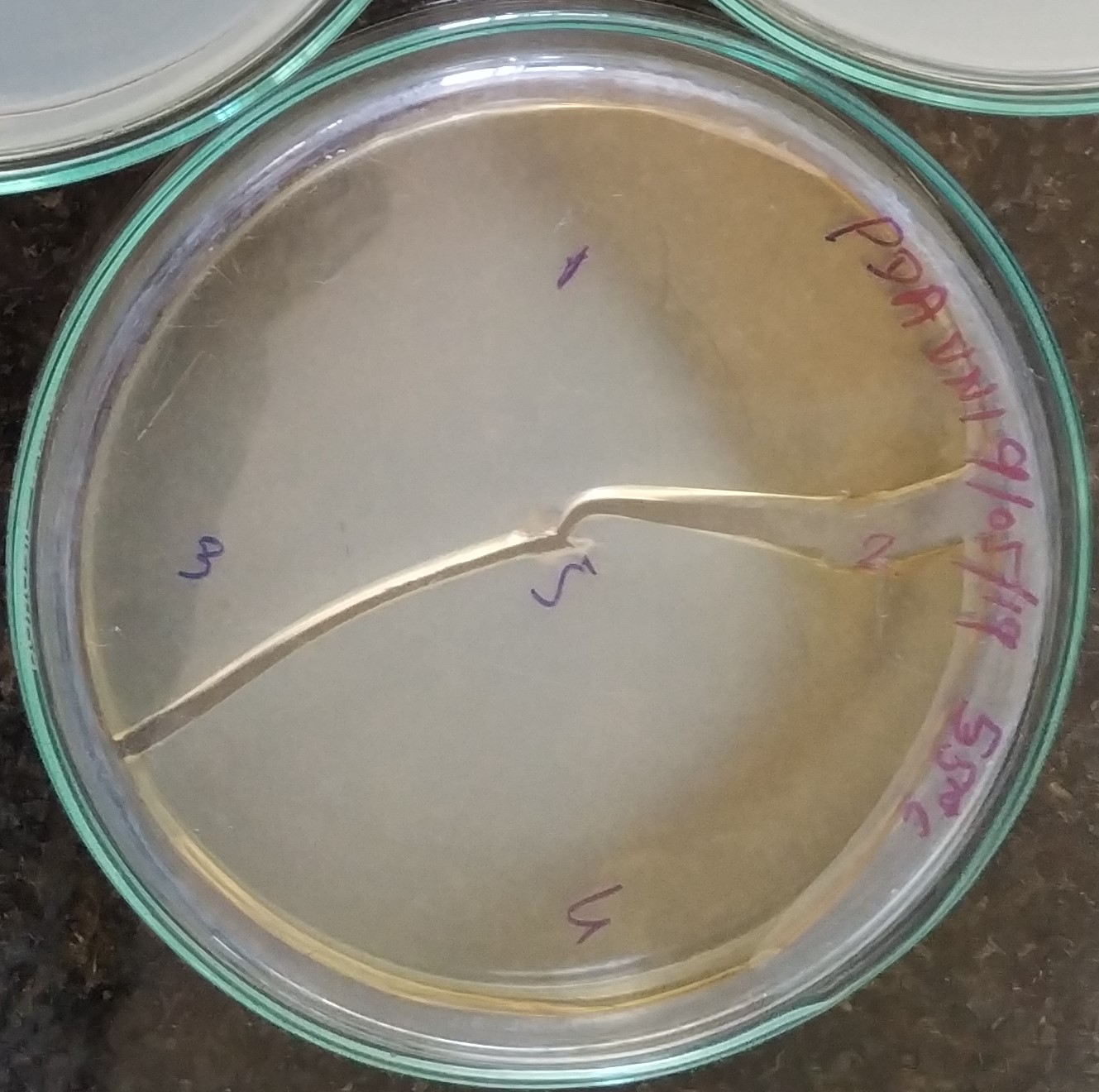

**D	E**

**Figure. 3** Growth observed after four days incubation on Potatoes dextrose agar medium at, **A.**20°C, **B.** 25°C, **C.** 30°C, **D.** 37°C and **E.** 55°C.

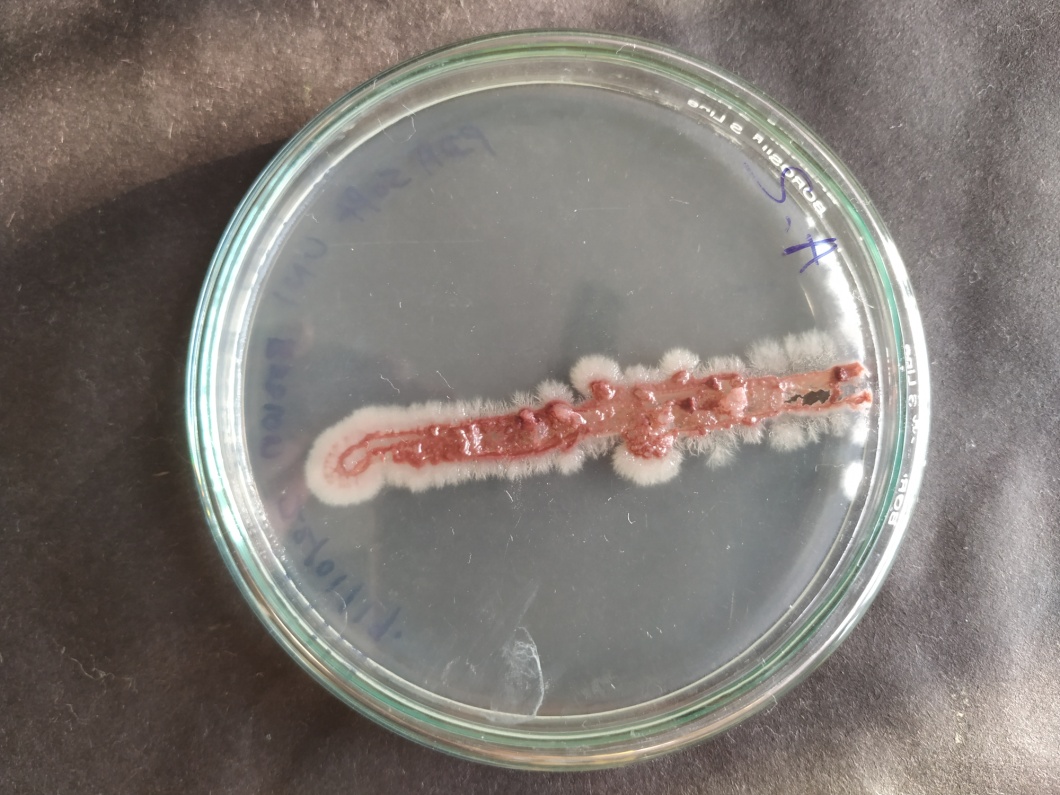

**Figure.4** Motility Assay

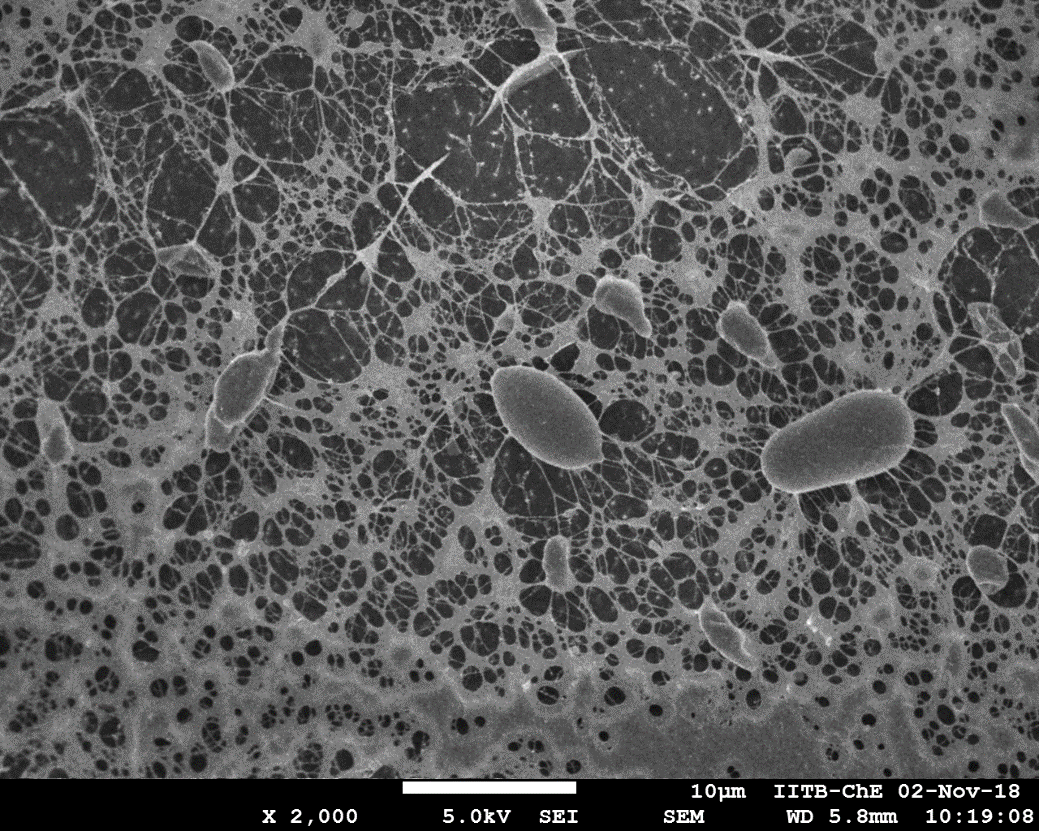

**Figure 5.** Cryo SEM images showing Cells trapped in Exopolysaccharide Network

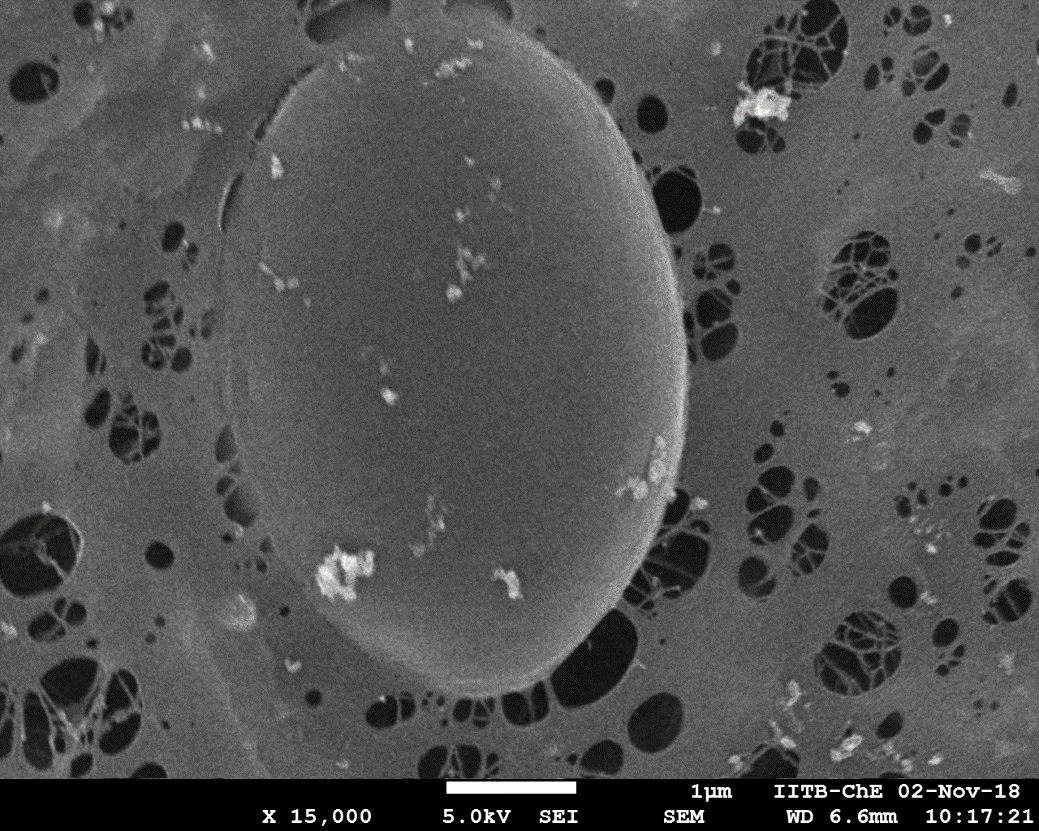

**Figure 6.** Cryo SEM images showing Yeast Cell partially embedded in Exopolysaccharide Network

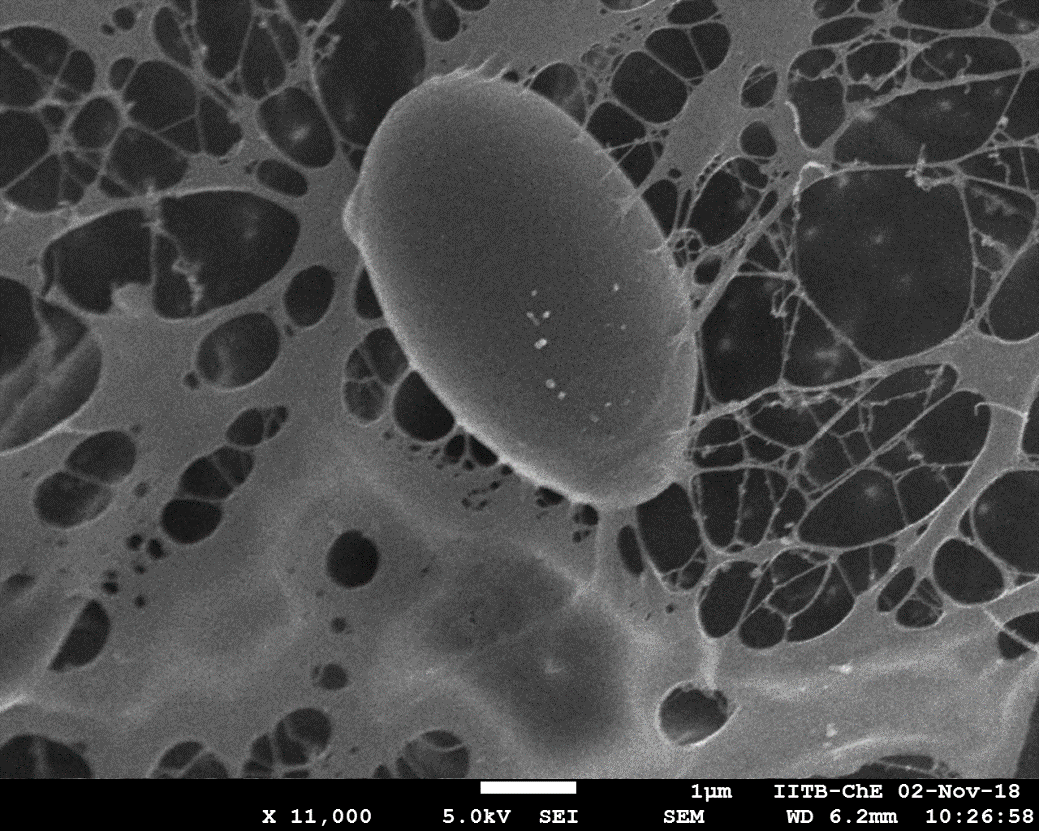

**Figure 7.** Cryo SEM images showing Cryo SEM images showing Strong Binding of Yeast cell with Exopolysaccharide Network

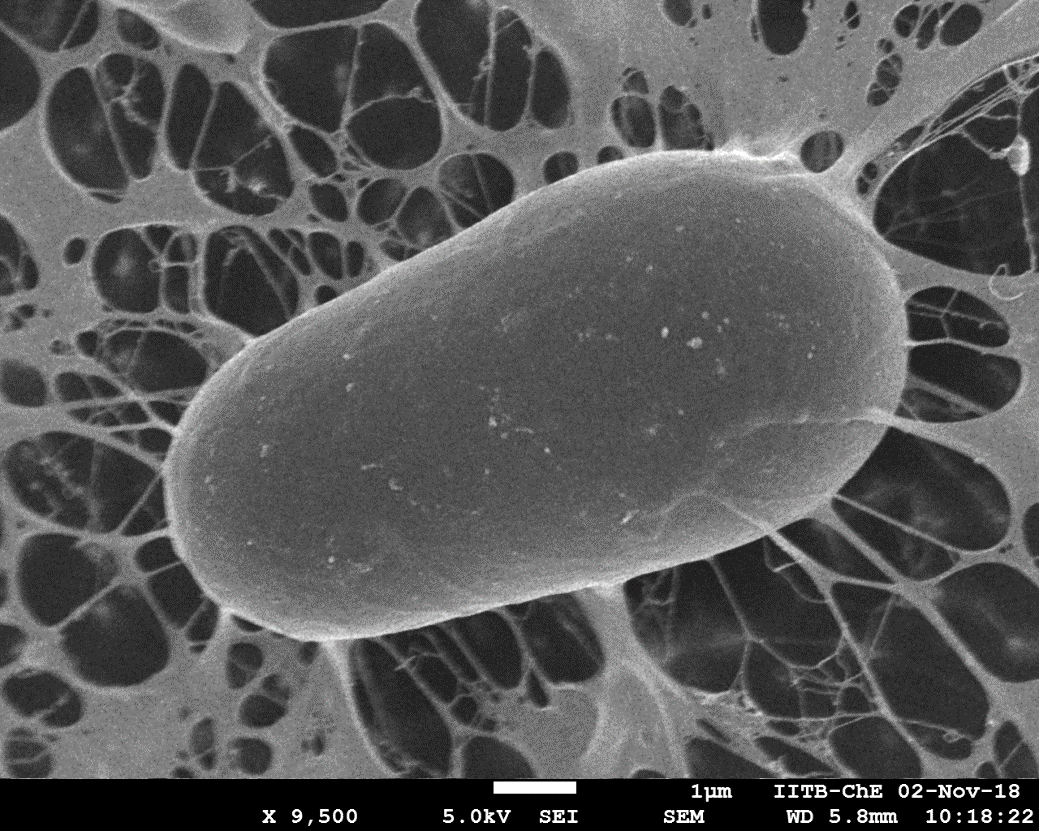

**Figure 8.** Cryo SEM images showing Cryo SEM images showing Strong Binding of Yeast cell with Exopolysaccharide Network

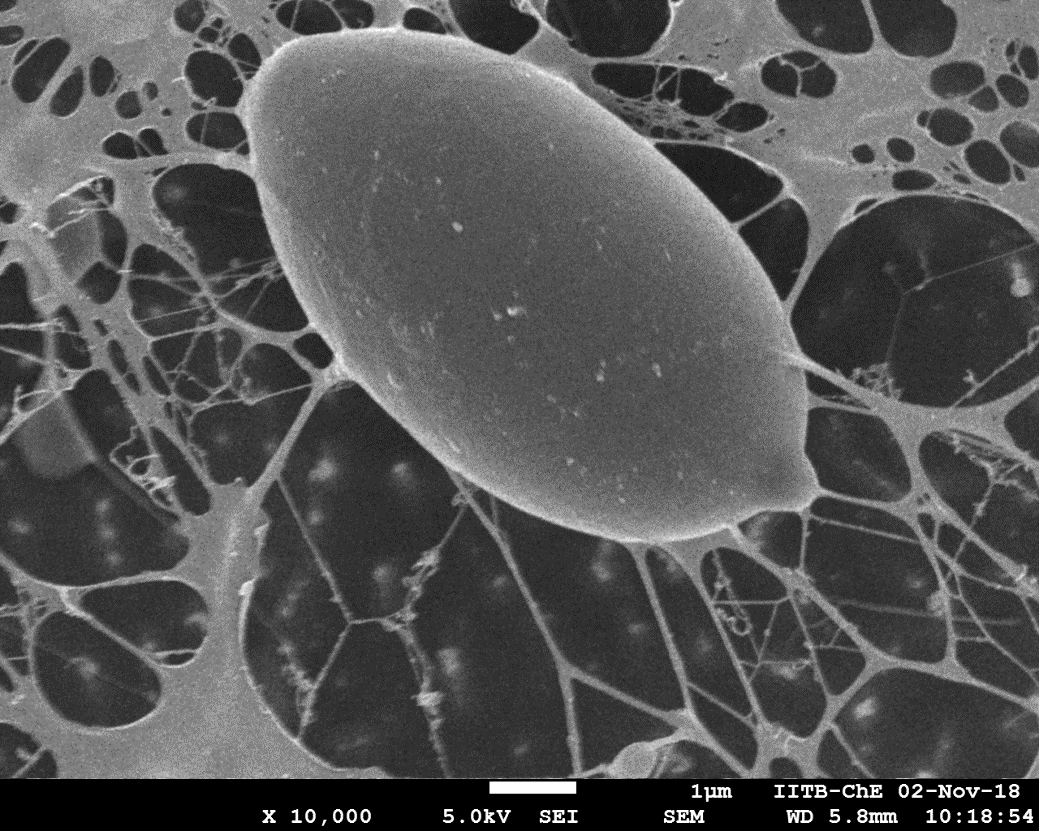

**Figure 9.** Cryo SEM images showing Strong Binding of Yeast cell with Exopolysaccharide Network

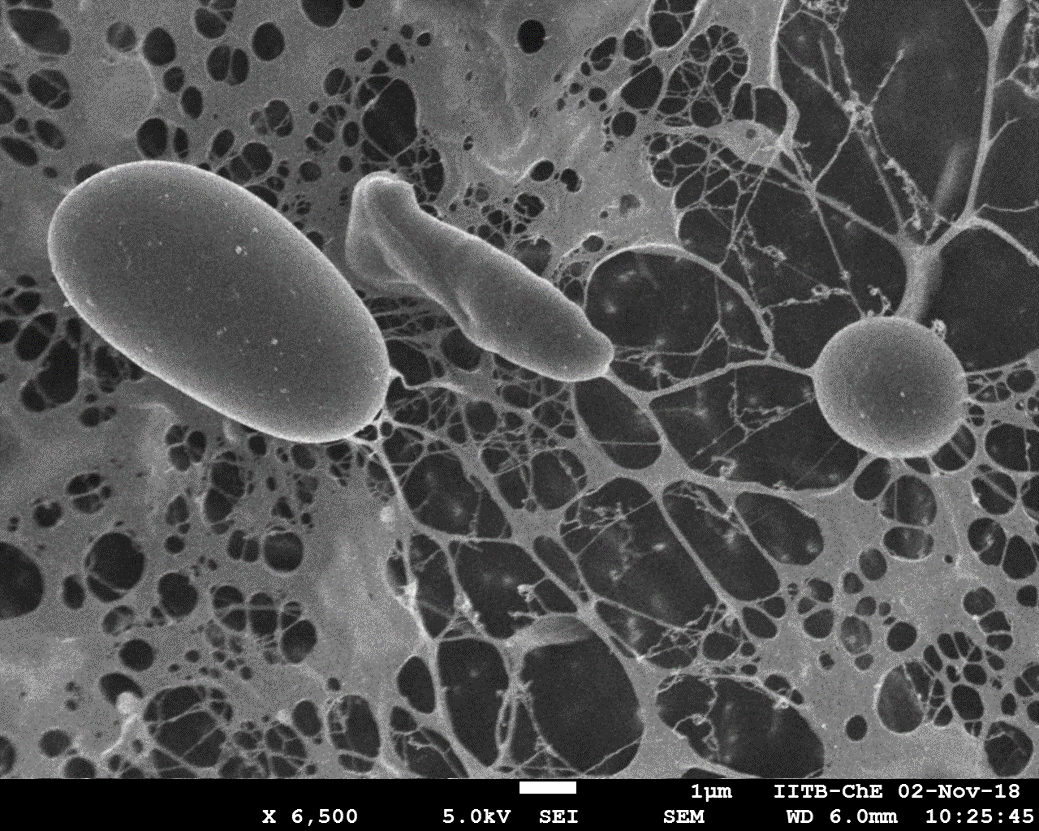

**Figure 10.** Cryo SEM images showing Weakened Exopolysaccharide Network and cell escaping

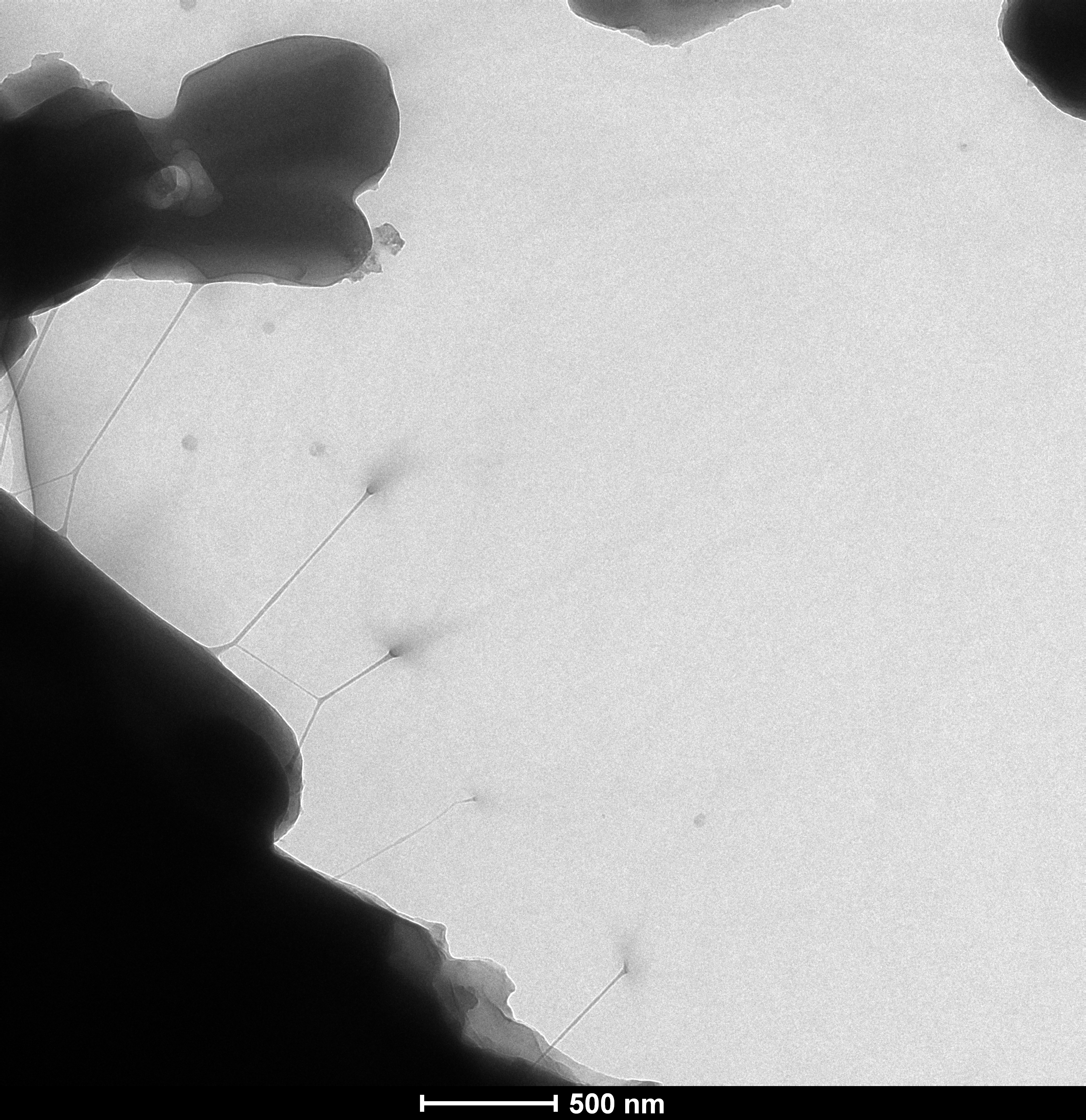

**Figure 11.** HR-TEM images showing Exopolysaccharide Network

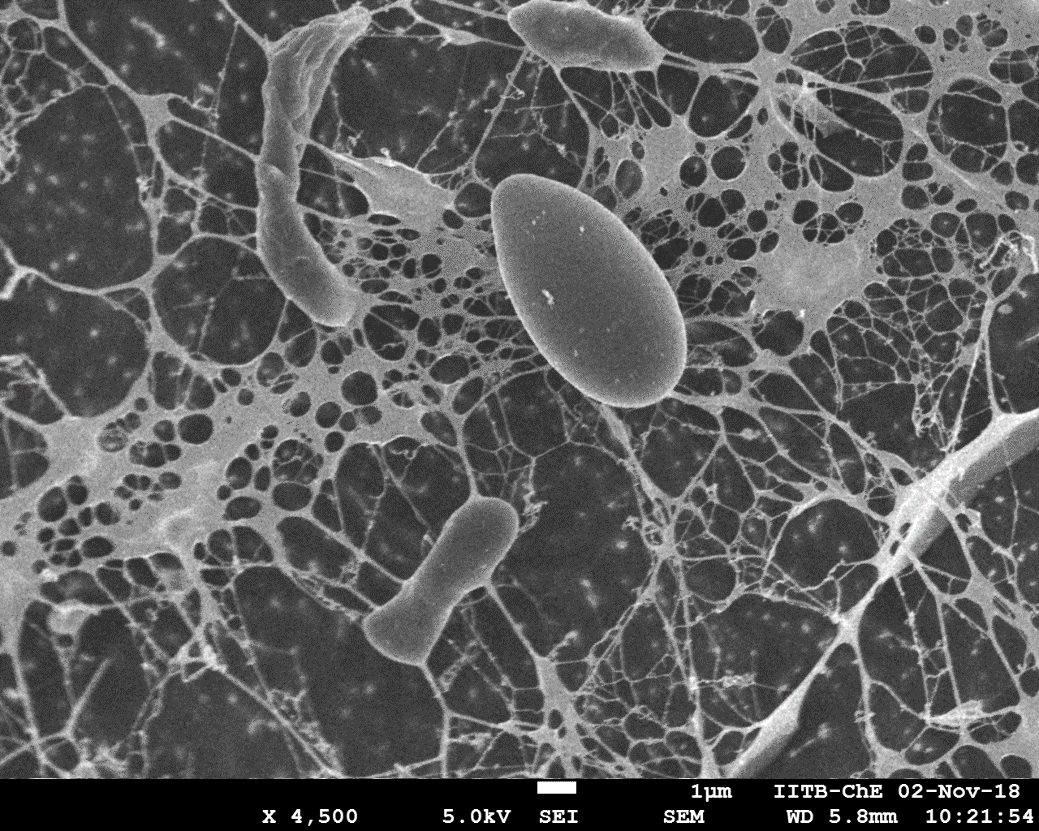

**Figure 12.** Cryo SEM images showing Escape of Yeast cell from Exopolysaccharide Network

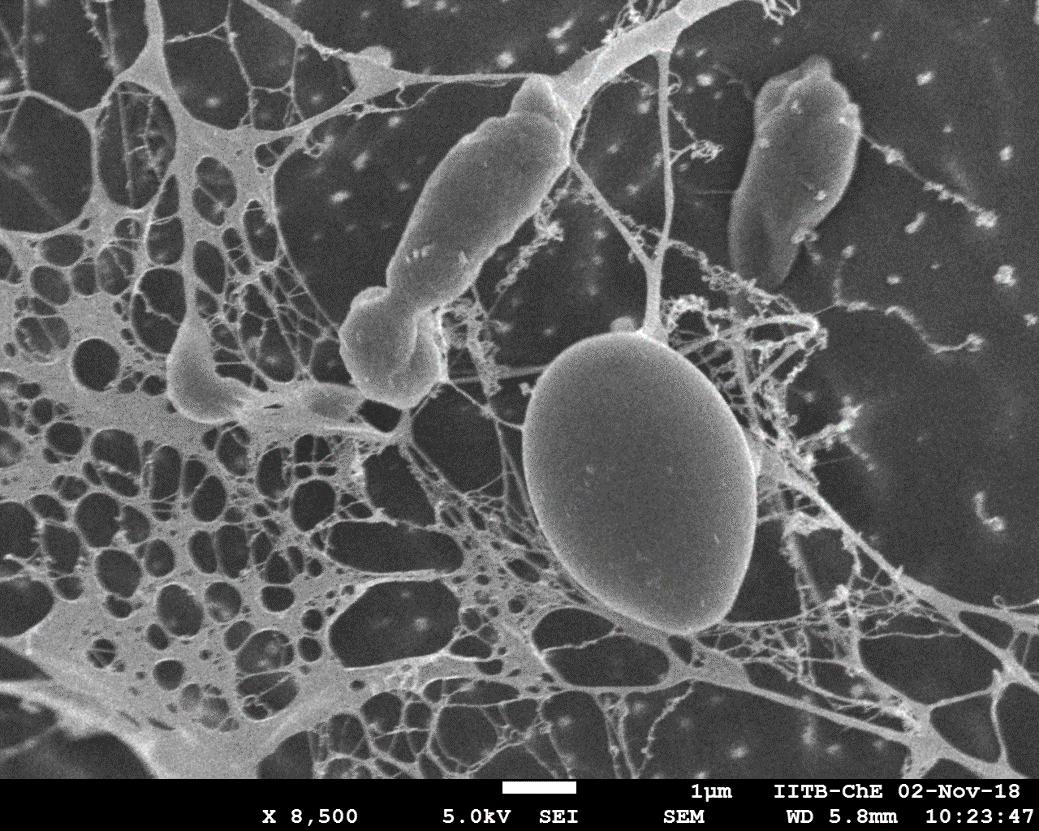

**Figure 13.** Cryo SEM images showing Escape of Yeast cell from Exopolysaccharide Network

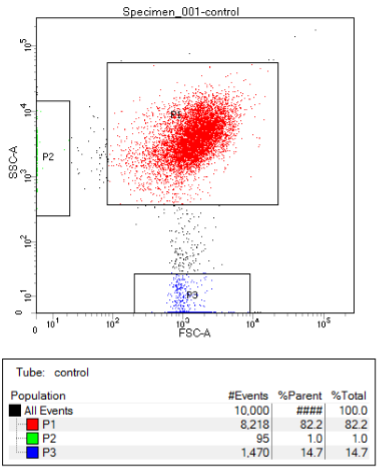

**Figure 14.** FACS analysis of intact Yeast cell

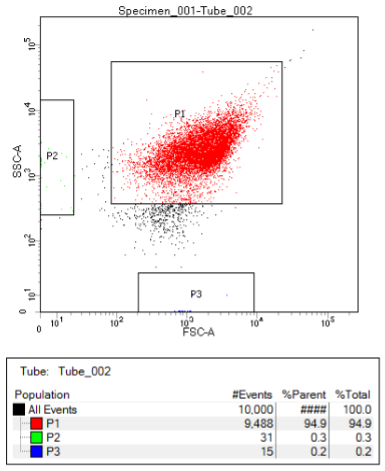

**Figure 15.** FACS analysis of Lysed Yeast cell
