## Supplementary Data 1A for "Moving Yeasts: Resolving the Mystery"

| Track n° | Slice n° | X | Y | Distance | Velocity | Pixel Value |
|---|---|---|---|---|---|---|
| 1 | 1 | 146 | 134 | -1 | -1 | 160 |
| 1 | 2 | 142 | 103 | 4.032 | 2.016 | 154 |
| 1 | 3 | 142 | 103 | 0 | 0 | 154 |
| 1 | 4 | 144 | 101 | 0.365 | 0.182 | 138 |
| 1 | 5 | 142 | 101 | 0.258 | 0.129 | 135 |
| 1 | 6 | 142 | 101 | 0 | 0 | 135 |
| 1 | 7 | 142 | 104 | 0.387 | 0.194 | 155 |
| 1 | 8 | 140 | 106 | 0.365 | 0.182 | 162 |
| 1 | 9 | 140 | 103 | 0.387 | 0.194 | 145 |
| 1 | 10 | 141 | 103 | 0.129 | 0.065 | 149 |
| 1 | 11 | 140 | 104 | 0.182 | 0.091 | 146 |
| 1 | 12 | 141 | 100 | 0.532 | 0.266 | 140 |
| 1 | 13 | 140 | 104 | 0.532 | 0.266 | 144 |
| 1 | 14 | 140 | 106 | 0.258 | 0.129 | 166 |
| 1 | 15 | 141 | 106 | 0.129 | 0.065 | 177 |
| 1 | 16 | 141 | 102 | 0.516 | 0.258 | 142 |
| 1 | 17 | 143 | 102 | 0.258 | 0.129 | 145 |
| 1 | 18 | 140 | 104 | 0.465 | 0.233 | 145 |
| 1 | 19 | 141 | 101 | 0.408 | 0.204 | 142 |
| 1 | 20 | 141 | 101 | 0 | 0 | 146 |
| 1 | 21 | 141 | 101 | 0 | 0 | 147 |
| 1 | 22 | 141 | 101 | 0 | 0 | 144 |
| 1 | 23 | 141 | 101 | 0 | 0 | 136 |
| 1 | 24 | 141 | 101 | 0 | 0 | 136 |
| 1 | 25 | 141 | 101 | 0 | 0 | 135 |
| 1 | 26 | 141 | 98 | 0.387 | 0.194 | 146 |
| 1 | 27 | 141 | 98 | 0 | 0 | 144 |
| 1 | 28 | 141 | 99 | 0.129 | 0.065 | 136 |
| 1 | 29 | 142 | 99 | 0.129 | 0.065 | 135 |
| 1 | 30 | 142 | 99 | 0 | 0 | 135 |
| 1 | 31 | 142 | 99 | 0 | 0 | 139 |
| 1 | 32 | 141 | 99 | 0.129 | 0.065 | 136 |
| 1 | 33 | 141 | 99 | 0 | 0 | 135 |
| 1 | 34 | 141 | 99 | 0 | 0 | 138 |
| 1 | 35 | 140 | 100 | 0.182 | 0.091 | 137 |
| 1 | 36 | 140 | 100 | 0 | 0 | 147 |
| 1 | 37 | 140 | 100 | 0 | 0 | 141 |
| 1 | 38 | 142 | 95 | 0.695 | 0.347 | 155 |
| 1 | 39 | 142 | 95 | 0 | 0 | 162 |
| 1 | 40 | 142 | 95 | 0 | 0 | 163 |
| 1 | 41 | 142 | 95 | 0 | 0 | 166 |
| 1 | 42 | 143 | 95 | 0.129 | 0.065 | 159 |
| 1 | 43 | 143 | 95 | 0 | 0 | 143 |
| 1 | 44 | 143 | 95 | 0 | 0 | 140 |
| 1 | 45 | 143 | 95 | 0 | 0 | 147 |

| | | | | | | |
|---|---|---|---|---|---|---|
| 1 | 46 | 142 | 97 | 0.288 | 0.144 | 135 |
| 1 | 47 | 142 | 97 | 0 | 0 | 137 |
| 1 | 48 | 142 | 97 | 0 | 0 | 135 |
| 1 | 49 | 142 | 97 | 0 | 0 | 136 |
| 1 | 50 | 141 | 98 | 0.182 | 0.091 | 144 |
| 1 | 51 | 139 | 98 | 0.258 | 0.129 | 138 |
| 1 | 52 | 139 | 96 | 0.258 | 0.129 | 146 |
| 1 | 53 | 140 | 99 | 0.408 | 0.204 | 160 |
| 1 | 54 | 140 | 97 | 0.258 | 0.129 | 135 |
| 1 | 55 | 140 | 96 | 0.129 | 0.065 | 135 |
| 1 | 56 | 138 | 97 | 0.288 | 0.144 | 137 |
| 1 | 57 | 138 | 97 | 0 | 0 | 138 |
| 1 | 58 | 138 | 97 | 0 | 0 | 143 |
| 1 | 59 | 138 | 97 | 0 | 0 | 142 |
| 1 | 60 | 138 | 97 | 0 | 0 | 139 |
| 1 | 61 | 138 | 97 | 0 | 0 | 139 |
| 1 | 62 | 138 | 97 | 0 | 0 | 141 |
| 1 | 63 | 137 | 97 | 0.129 | 0.065 | 137 |
| 1 | 64 | 137 | 97 | 0 | 0 | 139 |
| 1 | 65 | 137 | 97 | 0 | 0 | 137 |
| 1 | 66 | 137 | 97 | 0 | 0 | 137 |
| 1 | 67 | 137 | 97 | 0 | 0 | 141 |
| 1 | 68 | 137 | 97 | 0 | 0 | 138 |
| 1 | 69 | 137 | 97 | 0 | 0 | 137 |
| 1 | 70 | 137 | 97 | 0 | 0 | 140 |
| 1 | 71 | 137 | 97 | 0 | 0 | 136 |
| 1 | 72 | 137 | 97 | 0 | 0 | 138 |
| 1 | 73 | 137 | 97 | 0 | 0 | 140 |
| 1 | 74 | 137 | 97 | 0 | 0 | 145 |
| 1 | 75 | 137 | 97 | 0 | 0 | 157 |
| 1 | 76 | 137 | 97 | 0 | 0 | 139 |
| 1 | 77 | 137 | 97 | 0 | 0 | 139 |
| 1 | 78 | 137 | 97 | 0 | 0 | 148 |
| 1 | 79 | 137 | 97 | 0 | 0 | 141 |
| 1 | 80 | 137 | 97 | 0 | 0 | 142 |
| 1 | 81 | 137 | 97 | 0 | 0 | 144 |
| 1 | 82 | 137 | 97 | 0 | 0 | 144 |
| 1 | 83 | 137 | 97 | 0 | 0 | 146 |
| 1 | 84 | 137 | 97 | 0 | 0 | 141 |
| 1 | 85 | 137 | 97 | 0 | 0 | 151 |
| 1 | 86 | 137 | 97 | 0 | 0 | 144 |
| 1 | 87 | 137 | 97 | 0 | 0 | 147 |
| 1 | 88 | 137 | 97 | 0 | 0 | 158 |
| 1 | 89 | 137 | 97 | 0 | 0 | 158 |
| 1 | 90 | 137 | 97 | 0 | 0 | 156 |
| 1 | 91 | 137 | 97 | 0 | 0 | 145 |
| 1 | 92 | 137 | 97 | 0 | 0 | 151 |

| | | | | | | |
|---|---|---|---|---|---|---|
| 1 | 93 | 137 | 97 | 0 | 0 | 135 |
| 1 | 94 | 137 | 97 | 0 | 0 | 145 |
| 1 | 95 | 137 | 97 | 0 | 0 | 137 |
| 1 | 96 | 137 | 97 | 0 | 0 | 139 |
| 1 | 97 | 137 | 97 | 0 | 0 | 146 |
| 1 | 98 | 137 | 97 | 0 | 0 | 148 |
| 1 | 99 | 137 | 97 | 0 | 0 | 144 |
| 1 | 100 | 137 | 97 | 0 | 0 | 139 |
| 1 | 101 | 137 | 97 | 0 | 0 | 147 |
| 1 | 102 | 137 | 97 | 0 | 0 | 146 |
| 1 | 103 | 137 | 97 | 0 | 0 | 137 |
| 1 | 104 | 137 | 97 | 0 | 0 | 140 |
| 1 | 105 | 137 | 97 | 0 | 0 | 146 |
| 1 | 106 | 137 | 97 | 0 | 0 | 155 |
| 1 | 107 | 137 | 97 | 0 | 0 | 160 |
| 1 | 108 | 137 | 97 | 0 | 0 | 158 |
| 1 | 109 | 137 | 97 | 0 | 0 | 154 |
| 1 | 110 | 137 | 98 | 0.129 | 0.065 | 139 |
| 1 | 111 | 135 | 98 | 0.258 | 0.129 | 140 |
| 1 | 112 | 132 | 98 | 0.387 | 0.194 | 165 |
| 1 | 113 | 132 | 99 | 0.129 | 0.065 | 160 |
| 1 | 114 | 132 | 99 | 0 | 0 | 141 |
| 1 | 115 | 132 | 99 | 0 | 0 | 150 |
| 1 | 116 | 131 | 99 | 0.129 | 0.065 | 164 |
| 1 | 117 | 131 | 99 | 0 | 0 | 162 |
| 1 | 118 | 131 | 99 | 0 | 0 | 164 |
| 1 | 119 | 131 | 99 | 0 | 0 | 163 |
| 1 | 120 | 131 | 99 | 0 | 0 | 163 |
| 1 | 121 | 131 | 99 | 0 | 0 | 163 |
| 1 | 122 | 131 | 97 | 0.258 | 0.129 | 163 |
| 1 | 123 | 131 | 96 | 0.129 | 0.065 | 165 |
| 1 | 124 | 131 | 96 | 0 | 0 | 166 |
| 1 | 125 | 131 | 96 | 0 | 0 | 168 |
| 1 | 126 | 131 | 96 | 0 | 0 | 163 |
| 1 | 127 | 131 | 96 | 0 | 0 | 163 |
| 1 | 128 | 132 | 96 | 0.129 | 0.065 | 167 |
| 1 | 129 | 132 | 96 | 0 | 0 | 166 |
| 1 | 130 | 132 | 96 | 0 | 0 | 168 |
| 1 | 131 | 132 | 96 | 0 | 0 | 171 |
| 1 | 132 | 132 | 96 | 0 | 0 | 170 |
| 1 | 133 | 132 | 96 | 0 | 0 | 168 |
| 1 | 134 | 133 | 96 | 0.129 | 0.065 | 158 |
| 1 | 135 | 135 | 96 | 0.258 | 0.129 | 155 |
| 1 | 136 | 135 | 96 | 0 | 0 | 164 |
| 1 | 137 | 135 | 96 | 0 | 0 | 146 |
| 1 | 138 | 135 | 96 | 0 | 0 | 148 |
| 1 | 139 | 135 | 96 | 0 | 0 | 156 |

| | | | | | | |
|---|---|---|---|---|---|---|
| 1 | 140 | 136 | 97 | 0.182 | 0.091 | 154 |
| 1 | 141 | 136 | 97 | 0 | 0 | 137 |
| 1 | 142 | 136 | 97 | 0 | 0 | 139 |
| 1 | 143 | 136 | 97 | 0 | 0 | 144 |
| 1 | 144 | 136 | 97 | 0 | 0 | 137 |
| 1 | 145 | 136 | 97 | 0 | 0 | 136 |
| 1 | 146 | 136 | 97 | 0 | 0 | 137 |
| 1 | 147 | 136 | 97 | 0 | 0 | 154 |
| 1 | 148 | 136 | 97 | 0 | 0 | 153 |
| 1 | 149 | 136 | 97 | 0 | 0 | 136 |
| 1 | 150 | 127 | 100 | 1.224 | 0.612 | 166 |
| 1 | 151 | 129 | 100 | 0.258 | 0.129 | 145 |
| 1 | 152 | 129 | 100 | 0 | 0 | 135 |
| 1 | 153 | 129 | 99 | 0.129 | 0.065 | 156 |
| 1 | 154 | 129 | 98 | 0.129 | 0.065 | 166 |
| 1 | 155 | 129 | 98 | 0 | 0 | 169 |
| 1 | 156 | 128 | 98 | 0.129 | 0.065 | 164 |
| 1 | 157 | 128 | 98 | 0 | 0 | 166 |
| 1 | 158 | 128 | 100 | 0.258 | 0.129 | 168 |
| 1 | 159 | 128 | 100 | 0 | 0 | 168 |
| 1 | 160 | 128 | 100 | 0 | 0 | 168 |
| 1 | 161 | 130 | 98 | 0.365 | 0.182 | 164 |
| 1 | 162 | 130 | 98 | 0 | 0 | 165 |
| 1 | 163 | 130 | 98 | 0 | 0 | 165 |
| 1 | 164 | 130 | 97 | 0.129 | 0.065 | 167 |
| 1 | 165 | 130 | 97 | 0 | 0 | 170 |
| 1 | 166 | 130 | 97 | 0 | 0 | 170 |
| 1 | 167 | 130 | 97 | 0 | 0 | 168 |
| 1 | 168 | 130 | 97 | 0 | 0 | 159 |
| 1 | 169 | 130 | 96 | 0.129 | 0.065 | 167 |
| 1 | 170 | 130 | 96 | 0 | 0 | 163 |
| 1 | 171 | 130 | 96 | 0 | 0 | 159 |
| 1 | 172 | 130 | 95 | 0.129 | 0.065 | 167 |
| 1 | 173 | 131 | 95 | 0.129 | 0.065 | 152 |
| 1 | 174 | 131 | 95 | 0 | 0 | 166 |
| 1 | 175 | 132 | 95 | 0.129 | 0.065 | 155 |
| 1 | 176 | 132 | 95 | 0 | 0 | 152 |
| 1 | 177 | 132 | 95 | 0 | 0 | 142 |
| 1 | 178 | 132 | 92 | 0.387 | 0.194 | 165 |
| 1 | 179 | 131 | 95 | 0.408 | 0.204 | 151 |
| 1 | 180 | 131 | 96 | 0.129 | 0.065 | 133 |
| 1 | 181 | 131 | 96 | 0 | 0 | 142 |
| 1 | 182 | 131 | 96 | 0 | 0 | 138 |
| 1 | 183 | 131 | 96 | 0 | 0 | 140 |
| 1 | 184 | 131 | 96 | 0 | 0 | 135 |
| 1 | 185 | 131 | 96 | 0 | 0 | 137 |
| 1 | 186 | 128 | 95 | 0.408 | 0.204 | 166 |

| | | | | | | |
|---|---|---|---|---|---|---|
| 1 | 187 | 128 | 95 | 0 | 0 | 158 |
| 1 | 188 | 128 | 95 | 0 | 0 | 161 |
| 1 | 189 | 128 | 95 | 0 | 0 | 168 |
| 1 | 190 | 128 | 94 | 0.129 | 0.065 | 163 |
| 1 | 191 | 128 | 94 | 0 | 0 | 167 |
| 1 | 192 | 128 | 94 | 0 | 0 | 163 |
| 1 | 193 | 128 | 94 | 0 | 0 | 167 |
| 1 | 194 | 127 | 94 | 0.129 | 0.065 | 165 |
| 1 | 195 | 127 | 94 | 0 | 0 | 163 |
| 1 | 196 | 127 | 96 | 0.258 | 0.129 | 162 |
| 1 | 197 | 127 | 96 | 0 | 0 | 161 |
| 1 | 198 | 127 | 94 | 0.258 | 0.129 | 166 |
| 1 | 199 | 127 | 94 | 0 | 0 | 165 |
| 1 | 200 | 127 | 94 | 0 | 0 | 166 |
| 1 | 201 | 129 | 95 | 0.288 | 0.144 | 160 |
| 1 | 202 | 128 | 95 | 0.129 | 0.065 | 166 |
| 1 | 203 | 128 | 95 | 0 | 0 | 166 |
| 1 | 204 | 128 | 95 | 0 | 0 | 166 |
| 1 | 205 | 128 | 95 | 0 | 0 | 158 |
| 1 | 206 | 128 | 95 | 0 | 0 | 162 |
| 1 | 207 | 129 | 95 | 0.129 | 0.065 | 146 |
| 1 | 208 | 129 | 95 | 0 | 0 | 155 |
| 1 | 209 | 129 | 95 | 0 | 0 | 144 |
| 1 | 210 | 129 | 95 | 0 | 0 | 139 |
| 1 | 211 | 129 | 95 | 0 | 0 | 138 |
| 1 | 212 | 129 | 95 | 0 | 0 | 151 |
| 1 | 213 | 129 | 95 | 0 | 0 | 153 |
| 1 | 214 | 129 | 95 | 0 | 0 | 153 |
| 1 | 215 | 129 | 95 | 0 | 0 | 150 |
| 1 | 216 | 129 | 95 | 0 | 0 | 152 |
| 1 | 217 | 129 | 95 | 0 | 0 | 165 |
| 1 | 218 | 128 | 96 | 0.182 | 0.091 | 155 |
| 1 | 219 | 128 | 96 | 0 | 0 | 161 |
| 1 | 220 | 128 | 96 | 0 | 0 | 167 |
| 1 | 221 | 128 | 96 | 0 | 0 | 167 |
| 1 | 222 | 128 | 96 | 0 | 0 | 170 |
| 1 | 223 | 128 | 96 | 0 | 0 | 167 |
| 1 | 224 | 127 | 96 | 0.129 | 0.065 | 167 |
| 1 | 225 | 127 | 96 | 0 | 0 | 160 |
| 1 | 226 | 127 | 96 | 0 | 0 | 162 |
| 1 | 227 | 127 | 96 | 0 | 0 | 162 |
| 1 | 228 | 127 | 96 | 0 | 0 | 158 |
| 1 | 229 | 127 | 96 | 0 | 0 | 153 |
| 1 | 230 | 127 | 96 | 0 | 0 | 164 |
| 1 | 231 | 127 | 96 | 0 | 0 | 160 |
| 1 | 232 | 126 | 97 | 0.182 | 0.091 | 157 |
| 1 | 233 | 126 | 95 | 0.258 | 0.129 | 165 |

| | | | | | | |
|---|---|---|---|---|---|---|
| 1 | 234 | 126 | 95 | 0 | 0 | 163 |
| 1 | 235 | 126 | 95 | 0 | 0 | 165 |
| 1 | 236 | 126 | 95 | 0 | 0 | 158 |
| 1 | 237 | 126 | 95 | 0 | 0 | 161 |
| 1 | 238 | 126 | 94 | 0.129 | 0.065 | 166 |
| 1 | 239 | 126 | 94 | 0 | 0 | 165 |
| 1 | 240 | 125 | 94 | 0.129 | 0.065 | 160 |
| 1 | 241 | 125 | 94 | 0 | 0 | 165 |
| 1 | 242 | 125 | 94 | 0 | 0 | 164 |
| 1 | 243 | 125 | 94 | 0 | 0 | 163 |
| 1 | 244 | 125 | 94 | 0 | 0 | 165 |
| 1 | 245 | 125 | 94 | 0 | 0 | 165 |
| 1 | 246 | 125 | 94 | 0 | 0 | 165 |
| 1 | 247 | 125 | 94 | 0 | 0 | 165 |
| 1 | 248 | 125 | 94 | 0 | 0 | 165 |
| 1 | 249 | 125 | 94 | 0 | 0 | 166 |
| 1 | 250 | 125 | 92 | 0.258 | 0.129 | 164 |
| 1 | 251 | 125 | 92 | 0 | 0 | 165 |
| 1 | 252 | 125 | 92 | 0 | 0 | 165 |
| 1 | 253 | 125 | 92 | 0 | 0 | 165 |
| 1 | 254 | 125 | 92 | 0 | 0 | 166 |
| 1 | 255 | 125 | 92 | 0 | 0 | 165 |
| 1 | 256 | 125 | 92 | 0 | 0 | 166 |
| 1 | 257 | 125 | 92 | 0 | 0 | 166 |
| 1 | 258 | 125 | 92 | 0 | 0 | 167 |
| 1 | 259 | 125 | 92 | 0 | 0 | 166 |
| 1 | 260 | 125 | 92 | 0 | 0 | 165 |
| 1 | 261 | 125 | 92 | 0 | 0 | 161 |
| 1 | 262 | 125 | 92 | 0 | 0 | 165 |
| 1 | 263 | 125 | 92 | 0 | 0 | 164 |
| 1 | 264 | 125 | 92 | 0 | 0 | 164 |
| 1 | 265 | 125 | 92 | 0 | 0 | 159 |
| 1 | 266 | 125 | 92 | 0 | 0 | 163 |
| 1 | 267 | 125 | 92 | 0 | 0 | 164 |
| 1 | 268 | 125 | 92 | 0 | 0 | 163 |
| 1 | 269 | 125 | 92 | 0 | 0 | 163 |
| 1 | 270 | 125 | 92 | 0 | 0 | 164 |
| 1 | 271 | 125 | 92 | 0 | 0 | 163 |
| 1 | 272 | 125 | 91 | 0.129 | 0.065 | 161 |
| 1 | 273 | 125 | 91 | 0 | 0 | 160 |
| 1 | 274 | 125 | 91 | 0 | 0 | 160 |
| 1 | 275 | 125 | 91 | 0 | 0 | 161 |
| 1 | 276 | 125 | 91 | 0 | 0 | 163 |
| 1 | 277 | 125 | 91 | 0 | 0 | 164 |
| 1 | 278 | 125 | 91 | 0 | 0 | 165 |
| 1 | 279 | 125 | 91 | 0 | 0 | 163 |
| 1 | 280 | 125 | 91 | 0 | 0 | 162 |

| | | | | | | |
|---|---|---|---|---|---|---|
| 1 | 281 | 123 | 95 | 0.577 | 0.288 | 165 |
| 1 | 282 | 123 | 95 | 0 | 0 | 157 |
| 1 | 283 | 124 | 94 | 0.182 | 0.091 | 154 |
| 1 | 284 | 124 | 94 | 0 | 0 | 145 |
| 1 | 285 | 124 | 94 | 0 | 0 | 139 |
| 1 | 286 | 124 | 94 | 0 | 0 | 137 |
| 1 | 287 | 124 | 94 | 0 | 0 | 140 |
| 1 | 288 | 124 | 93 | 0.129 | 0.065 | 152 |
| 1 | 289 | 124 | 93 | 0 | 0 | 144 |
| 1 | 290 | 124 | 93 | 0 | 0 | 144 |
| 1 | 291 | 124 | 93 | 0 | 0 | 140 |
| 1 | 292 | 124 | 93 | 0 | 0 | 139 |
| 1 | 293 | 124 | 93 | 0 | 0 | 143 |
| 1 | 294 | 124 | 93 | 0 | 0 | 144 |
| 1 | 295 | 124 | 93 | 0 | 0 | 149 |
| 1 | 296 | 124 | 93 | 0 | 0 | 146 |
| 1 | 297 | 124 | 93 | 0 | 0 | 150 |
| 1 | 298 | 124 | 93 | 0 | 0 | 164 |
| 1 | 299 | 123 | 93 | 0.129 | 0.065 | 159 |
| 1 | 300 | 123 | 93 | 0 | 0 | 156 |
| 1 | 301 | 123 | 93 | 0 | 0 | 153 |
| 1 | 302 | 123 | 93 | 0 | 0 | 149 |
| 1 | 303 | 123 | 93 | 0 | 0 | 149 |
| 1 | 304 | 123 | 93 | 0 | 0 | 149 |
| 1 | 305 | 123 | 93 | 0 | 0 | 149 |
| 1 | 306 | 123 | 93 | 0 | 0 | 152 |
| 1 | 307 | 123 | 93 | 0 | 0 | 146 |
| 1 | 308 | 123 | 93 | 0 | 0 | 149 |
| 1 | 309 | 123 | 93 | 0 | 0 | 155 |
| 1 | 310 | 123 | 93 | 0 | 0 | 145 |
| 1 | 311 | 123 | 93 | 0 | 0 | 144 |
| 1 | 312 | 123 | 93 | 0 | 0 | 144 |
| 1 | 313 | 122 | 93 | 0.129 | 0.065 | 161 |
| 1 | 314 | 122 | 93 | 0 | 0 | 156 |
| 1 | 315 | 122 | 93 | 0 | 0 | 148 |
| 1 | 316 | 122 | 93 | 0 | 0 | 145 |
| 1 | 317 | 122 | 93 | 0 | 0 | 151 |
| 1 | 318 | 122 | 93 | 0 | 0 | 139 |
| 1 | 319 | 122 | 93 | 0 | 0 | 137 |
| 1 | 320 | 122 | 93 | 0 | 0 | 138 |
| 1 | 321 | 122 | 93 | 0 | 0 | 140 |
| 1 | 322 | 122 | 93 | 0 | 0 | 142 |
| 1 | 323 | 122 | 92 | 0.129 | 0.065 | 148 |
| 1 | 324 | 122 | 92 | 0 | 0 | 139 |
| 1 | 325 | 124 | 91 | 0.288 | 0.144 | 137 |
| 1 | 326 | 125 | 90 | 0.182 | 0.091 | 139 |
| 1 | 327 | 122 | 90 | 0.387 | 0.194 | 156 |

| | | | | | | |
|---|---|---|---|---|---|---|
| 1 | 328 | 122 | 90 | 0 | 0 | 152 |
| 1 | 329 | 122 | 90 | 0 | 0 | 151 |
| 1 | 330 | 122 | 90 | 0 | 0 | 154 |
| 1 | 331 | 122 | 90 | 0 | 0 | 146 |
| 1 | 332 | 122 | 90 | 0 | 0 | 151 |
| 1 | 333 | 122 | 90 | 0 | 0 | 139 |
| 1 | 334 | 122 | 90 | 0 | 0 | 139 |
| 1 | 335 | 123 | 87 | 0.408 | 0.204 | 155 |
| 1 | 336 | 123 | 87 | 0 | 0 | 155 |
| 1 | 337 | 123 | 87 | 0 | 0 | 153 |
| 1 | 338 | 123 | 87 | 0 | 0 | 149 |
| 1 | 339 | 123 | 87 | 0 | 0 | 157 |
| 1 | 340 | 123 | 87 | 0 | 0 | 149 |
| 1 | 341 | 123 | 87 | 0 | 0 | 149 |
| 1 | 342 | 120 | 87 | 0.387 | 0.194 | 159 |
| 1 | 343 | 120 | 87 | 0 | 0 | 160 |
| 1 | 344 | 120 | 87 | 0 | 0 | 158 |
| 1 | 345 | 120 | 87 | 0 | 0 | 159 |
| 1 | 346 | 120 | 87 | 0 | 0 | 157 |
| 1 | 347 | 120 | 87 | 0 | 0 | 157 |
| 1 | 348 | 120 | 87 | 0 | 0 | 152 |
| 1 | 349 | 120 | 87 | 0 | 0 | 160 |
| 1 | 350 | 120 | 87 | 0 | 0 | 156 |
| 1 | 351 | 120 | 87 | 0 | 0 | 162 |
| 1 | 352 | 120 | 87 | 0 | 0 | 154 |
| 1 | 353 | 120 | 87 | 0 | 0 | 148 |
| 1 | 354 | 117 | 85 | 0.465 | 0.233 | 163 |
| 1 | 355 | 117 | 85 | 0 | 0 | 162 |
| 1 | 356 | 121 | 84 | 0.532 | 0.266 | 161 |
| 1 | 357 | 121 | 84 | 0 | 0 | 163 |
| 1 | 358 | 121 | 84 | 0 | 0 | 165 |
| 1 | 359 | 121 | 84 | 0 | 0 | 158 |
| 1 | 360 | 123 | 84 | 0.258 | 0.129 | 162 |
| 1 | 361 | 122 | 83 | 0.182 | 0.091 | 167 |
| 1 | 362 | 119 | 83 | 0.387 | 0.194 | 165 |
| 1 | 363 | 119 | 83 | 0 | 0 | 165 |
| 1 | 364 | 119 | 83 | 0 | 0 | 163 |
| 1 | 365 | 119 | 83 | 0 | 0 | 163 |
| 1 | 366 | 119 | 83 | 0 | 0 | 163 |
| 1 | 367 | 119 | 83 | 0 | 0 | 165 |
| 1 | 368 | 119 | 83 | 0 | 0 | 165 |
| 1 | 369 | 119 | 83 | 0 | 0 | 166 |
| 1 | 370 | 119 | 83 | 0 | 0 | 163 |
| 1 | 371 | 119 | 83 | 0 | 0 | 162 |
| 1 | 372 | 119 | 83 | 0 | 0 | 166 |
| 1 | 373 | 119 | 83 | 0 | 0 | 165 |
| 1 | 374 | 119 | 83 | 0 | 0 | 165 |

| | | | | | | |
|---|---|---|---|---|---|---|
| 1 | 375 | 119 | 86 | 0.387 | 0.194 | 156 |
| 1 | 376 | 119 | 86 | 0 | 0 | 155 |
| 1 | 377 | 119 | 86 | 0 | 0 | 158 |
| 1 | 378 | 119 | 86 | 0 | 0 | 161 |
| 1 | 379 | 119 | 86 | 0 | 0 | 142 |
| 1 | 380 | 119 | 86 | 0 | 0 | 151 |
| 1 | 381 | 120 | 84 | 0.288 | 0.144 | 167 |
| 1 | 382 | 120 | 83 | 0.129 | 0.065 | 165 |
| 1 | 383 | 120 | 87 | 0.516 | 0.258 | 135 |
| 1 | 384 | 120 | 87 | 0 | 0 | 133 |
| 1 | 385 | 120 | 87 | 0 | 0 | 147 |
| 1 | 386 | 120 | 87 | 0 | 0 | 136 |
| 1 | 387 | 119 | 88 | 0.182 | 0.091 | 144 |
| 1 | 388 | 119 | 88 | 0 | 0 | 138 |
| 1 | 389 | 119 | 88 | 0 | 0 | 146 |
| 1 | 390 | 119 | 88 | 0 | 0 | 137 |
| 1 | 391 | 117 | 86 | 0.365 | 0.182 | 159 |
| 1 | 392 | 117 | 86 | 0 | 0 | 162 |
| 1 | 393 | 117 | 86 | 0 | 0 | 151 |
| 1 | 394 | 117 | 86 | 0 | 0 | 156 |
| 1 | 395 | 117 | 86 | 0 | 0 | 163 |
| 1 | 396 | 117 | 86 | 0 | 0 | 161 |
| 1 | 397 | 117 | 86 | 0 | 0 | 149 |
| 1 | 398 | 117 | 86 | 0 | 0 | 154 |
| 1 | 399 | 117 | 86 | 0 | 0 | 145 |
| 1 | 400 | 117 | 86 | 0 | 0 | 159 |
| 1 | 401 | 117 | 86 | 0 | 0 | 158 |
| 1 | 402 | 117 | 86 | 0 | 0 | 161 |
| 1 | 403 | 117 | 86 | 0 | 0 | 162 |
| 1 | 404 | 114 | 87 | 0.408 | 0.204 | 162 |
| 1 | 405 | 114 | 87 | 0 | 0 | 158 |
| 1 | 406 | 114 | 87 | 0 | 0 | 162 |
| 1 | 407 | 116 | 87 | 0.258 | 0.129 | 146 |
| 1 | 408 | 116 | 87 | 0 | 0 | 148 |
| 1 | 409 | 116 | 87 | 0 | 0 | 134 |
| 1 | 410 | 116 | 87 | 0 | 0 | 134 |
| 1 | 411 | 116 | 87 | 0 | 0 | 131 |
| 1 | 412 | 116 | 87 | 0 | 0 | 131 |
| 1 | 413 | 116 | 87 | 0 | 0 | 132 |
| 1 | 414 | 116 | 87 | 0 | 0 | 132 |
| 1 | 415 | 116 | 87 | 0 | 0 | 129 |
| 1 | 416 | 116 | 87 | 0 | 0 | 132 |
| 1 | 417 | 116 | 87 | 0 | 0 | 135 |
| 1 | 418 | 116 | 87 | 0 | 0 | 131 |
| 1 | 419 | 114 | 87 | 0.258 | 0.129 | 131 |
| 1 | 420 | 114 | 87 | 0 | 0 | 133 |
| 1 | 421 | 114 | 87 | 0 | 0 | 135 |

| | | | | | | |
|---|---|---|---|---|---|---|
| 1 | 422 | 114 | 87 | 0 | 0 | 133 |
| 1 | 423 | 114 | 87 | 0 | 0 | 134 |
| 1 | 424 | 114 | 87 | 0 | 0 | 135 |
| 1 | 425 | 114 | 87 | 0 | 0 | 134 |
| 1 | 426 | 114 | 87 | 0 | 0 | 134 |
| 1 | 427 | 114 | 87 | 0 | 0 | 134 |
| 1 | 428 | 114 | 87 | 0 | 0 | 137 |
| 1 | 429 | 113 | 86 | 0.182 | 0.091 | 149 |
| 1 | 430 | 113 | 86 | 0 | 0 | 147 |
| 1 | 431 | 113 | 86 | 0 | 0 | 141 |
| 1 | 432 | 113 | 86 | 0 | 0 | 141 |
| 1 | 433 | 113 | 86 | 0 | 0 | 139 |
| 1 | 434 | 113 | 86 | 0 | 0 | 137 |
| 1 | 435 | 113 | 86 | 0 | 0 | 140 |
| 1 | 436 | 113 | 86 | 0 | 0 | 138 |
| 1 | 437 | 113 | 86 | 0 | 0 | 139 |
| 1 | 438 | 113 | 86 | 0 | 0 | 141 |
| 1 | 439 | 113 | 86 | 0 | 0 | 137 |
| 1 | 440 | 113 | 86 | 0 | 0 | 138 |
| 1 | 441 | 113 | 86 | 0 | 0 | 135 |
| 1 | 442 | 113 | 86 | 0 | 0 | 138 |
| 1 | 443 | 113 | 86 | 0 | 0 | 138 |
| 1 | 444 | 113 | 86 | 0 | 0 | 137 |
| 1 | 445 | 113 | 86 | 0 | 0 | 135 |
| 1 | 446 | 113 | 86 | 0 | 0 | 133 |
| 1 | 447 | 113 | 86 | 0 | 0 | 135 |
| 1 | 448 | 113 | 86 | 0 | 0 | 131 |
| 1 | 449 | 113 | 86 | 0 | 0 | 131 |
| 1 | 450 | 113 | 85 | 0.129 | 0.065 | 137 |
| 1 | 451 | 113 | 85 | 0 | 0 | 132 |
| 1 | 452 | 113 | 85 | 0 | 0 | 132 |
| 1 | 453 | 113 | 85 | 0 | 0 | 132 |
| 1 | 454 | 113 | 85 | 0 | 0 | 134 |
| 1 | 455 | 113 | 85 | 0 | 0 | 138 |
| 1 | 456 | 113 | 83 | 0.258 | 0.129 | 139 |
| 1 | 457 | 113 | 83 | 0 | 0 | 140 |
| 1 | 458 | 113 | 83 | 0 | 0 | 139 |
| 1 | 459 | 113 | 83 | 0 | 0 | 135 |
| 1 | 460 | 113 | 83 | 0 | 0 | 135 |
| 1 | 461 | 113 | 83 | 0 | 0 | 145 |
| 1 | 462 | 113 | 83 | 0 | 0 | 144 |
| 1 | 463 | 113 | 83 | 0 | 0 | 147 |
| 1 | 464 | 113 | 83 | 0 | 0 | 169 |
| 1 | 465 | 113 | 83 | 0 | 0 | 167 |
| 1 | 466 | 113 | 83 | 0 | 0 | 165 |
| 1 | 467 | 112 | 84 | 0.182 | 0.091 | 139 |
| 1 | 468 | 112 | 84 | 0 | 0 | 139 |

| | | | | | | |
|---|---|---|---|---|---|---|
| 1 | 469 | 112 | 84 | 0 | 0 | 142 |
| 1 | 470 | 112 | 84 | 0 | 0 | 140 |
| 1 | 471 | 113 | 84 | 0.129 | 0.065 | 149 |
| 1 | 472 | 113 | 84 | 0 | 0 | 154 |
| 1 | 473 | 113 | 84 | 0 | 0 | 149 |
| 1 | 474 | 109 | 80 | 0.73 | 0.365 | 167 |
| 1 | 475 | 109 | 80 | 0 | 0 | 166 |
| 1 | 476 | 109 | 80 | 0 | 0 | 168 |
| 1 | 477 | 109 | 80 | 0 | 0 | 167 |
| 1 | 478 | 111 | 80 | 0.258 | 0.129 | 167 |
| 1 | 479 | 112 | 80 | 0.129 | 0.065 | 160 |
| 1 | 480 | 112 | 80 | 0 | 0 | 154 |
| 1 | 481 | 112 | 80 | 0 | 0 | 152 |
| 1 | 482 | 111 | 79 | 0.182 | 0.091 | 170 |
| 1 | 483 | 111 | 79 | 0 | 0 | 170 |
| 1 | 484 | 110 | 79 | 0.129 | 0.065 | 169 |
| 1 | 485 | 110 | 79 | 0 | 0 | 162 |
| 1 | 486 | 110 | 82 | 0.387 | 0.194 | 168 |
| 1 | 487 | 110 | 82 | 0 | 0 | 168 |
| 1 | 488 | 110 | 82 | 0 | 0 | 166 |
| 1 | 489 | 110 | 82 | 0 | 0 | 167 |
| 1 | 490 | 110 | 82 | 0 | 0 | 141 |
| 1 | 491 | 110 | 82 | 0 | 0 | 138 |
| 1 | 492 | 110 | 82 | 0 | 0 | 140 |
| 1 | 493 | 110 | 81 | 0.129 | 0.065 | 143 |
| 1 | 494 | 110 | 81 | 0 | 0 | 142 |
| 1 | 495 | 110 | 81 | 0 | 0 | 140 |
| 1 | 496 | 110 | 81 | 0 | 0 | 144 |
| 1 | 497 | 110 | 81 | 0 | 0 | 143 |
| 1 | 498 | 110 | 81 | 0 | 0 | 145 |
| 1 | 499 | 110 | 81 | 0 | 0 | 145 |
| 1 | 500 | 110 | 81 | 0 | 0 | 145 |
| 1 | 501 | 110 | 80 | 0.129 | 0.065 | 144 |
| 1 | 502 | 110 | 80 | 0 | 0 | 147 |
| 1 | 503 | 110 | 80 | 0 | 0 | 141 |
| 1 | 504 | 110 | 80 | 0 | 0 | 145 |
| 1 | 505 | 110 | 80 | 0 | 0 | 139 |
| 1 | 506 | 110 | 80 | 0 | 0 | 137 |
| 1 | 507 | 110 | 80 | 0 | 0 | 138 |
| 1 | 508 | 110 | 80 | 0 | 0 | 143 |
| 1 | 509 | 109 | 80 | 0.129 | 0.065 | 138 |
| 1 | 510 | 107 | 77 | 0.465 | 0.233 | 165 |
| 1 | 511 | 107 | 77 | 0 | 0 | 163 |
| 1 | 512 | 107 | 77 | 0 | 0 | 165 |
| 1 | 513 | 107 | 77 | 0 | 0 | 162 |
| 1 | 514 | 107 | 77 | 0 | 0 | 161 |
| 1 | 515 | 107 | 77 | 0 | 0 | 162 |

| | | | | | | |
|---|---|---|---|---|---|---|
| 1 | 516 | 107 | 77 | 0 | 0 | 163 |
| 1 | 517 | 108 | 78 | 0.182 | 0.091 | 163 |
| 1 | 518 | 108 | 78 | 0 | 0 | 159 |
| 1 | 519 | 108 | 78 | 0 | 0 | 139 |
| 1 | 520 | 108 | 77 | 0.129 | 0.065 | 140 |
| 1 | 521 | 108 | 78 | 0.129 | 0.065 | 161 |
| 1 | 522 | 108 | 78 | 0 | 0 | 156 |
| 1 | 523 | 109 | 77 | 0.182 | 0.091 | 151 |
| 1 | 524 | 109 | 76 | 0.129 | 0.065 | 156 |
| 1 | 525 | 109 | 76 | 0 | 0 | 165 |
| 1 | 526 | 109 | 76 | 0 | 0 | 164 |
| 1 | 527 | 109 | 75 | 0.129 | 0.065 | 163 |
| 1 | 528 | 107 | 77 | 0.365 | 0.182 | 166 |
| 1 | 529 | 107 | 77 | 0 | 0 | 168 |
| 1 | 530 | 107 | 77 | 0 | 0 | 165 |
| 1 | 531 | 107 | 77 | 0 | 0 | 165 |
| 1 | 532 | 107 | 77 | 0 | 0 | 161 |
| 1 | 533 | 107 | 77 | 0 | 0 | 161 |
| 1 | 534 | 107 | 77 | 0 | 0 | 161 |
| 1 | 535 | 105 | 80 | 0.465 | 0.233 | 161 |
| 1 | 536 | 105 | 80 | 0 | 0 | 165 |
| 1 | 537 | 105 | 80 | 0 | 0 | 159 |
| 1 | 538 | 105 | 80 | 0 | 0 | 140 |
| 1 | 539 | 105 | 80 | 0 | 0 | 158 |
| 1 | 540 | 106 | 78 | 0.288 | 0.144 | 153 |
| 1 | 541 | 106 | 78 | 0 | 0 | 161 |
| 1 | 542 | 106 | 78 | 0 | 0 | 152 |
| 1 | 543 | 106 | 78 | 0 | 0 | 155 |
| 1 | 544 | 105 | 75 | 0.408 | 0.204 | 159 |
| 1 | 545 | 105 | 75 | 0 | 0 | 160 |
| 1 | 546 | 105 | 75 | 0 | 0 | 160 |
| 1 | 547 | 105 | 75 | 0 | 0 | 164 |
| 1 | 548 | 105 | 75 | 0 | 0 | 161 |
| 1 | 549 | 105 | 75 | 0 | 0 | 164 |
| 1 | 550 | 105 | 75 | 0 | 0 | 158 |
| 1 | 551 | 107 | 76 | 0.288 | 0.144 | 140 |
| 1 | 552 | 107 | 76 | 0 | 0 | 145 |
| 1 | 553 | 107 | 76 | 0 | 0 | 146 |
| 1 | 554 | 107 | 75 | 0.129 | 0.065 | 149 |
| 1 | 555 | 107 | 75 | 0 | 0 | 142 |
| 1 | 556 | 107 | 75 | 0 | 0 | 143 |
| 1 | 557 | 107 | 75 | 0 | 0 | 142 |
| 1 | 558 | 107 | 75 | 0 | 0 | 140 |
| 1 | 559 | 107 | 75 | 0 | 0 | 142 |
| 1 | 560 | 107 | 75 | 0 | 0 | 144 |
| 1 | 561 | 107 | 75 | 0 | 0 | 140 |
| 1 | 562 | 107 | 75 | 0 | 0 | 134 |

| | | | | | | |
|---|---|---|---|---|---|---|
| 1 | 563 | 107 | 75 | 0 | 0 | 134 |
| 1 | 564 | 107 | 75 | 0 | 0 | 135 |
| 1 | 565 | 104 | 73 | 0.465 | 0.233 | 160 |
| 1 | 566 | 104 | 73 | 0 | 0 | 160 |
| 1 | 567 | 105 | 73 | 0.129 | 0.065 | 160 |
| 1 | 568 | 105 | 73 | 0 | 0 | 161 |
| 1 | 569 | 105 | 73 | 0 | 0 | 161 |
| 1 | 570 | 105 | 73 | 0 | 0 | 160 |
| 1 | 571 | 105 | 73 | 0 | 0 | 158 |
| 1 | 572 | 105 | 79 | 0.774 | 0.387 | 145 |
| 1 | 573 | 105 | 79 | 0 | 0 | 141 |
| 1 | 574 | 105 | 79 | 0 | 0 | 143 |
| 1 | 575 | 105 | 79 | 0 | 0 | 140 |
| 1 | 576 | 105 | 79 | 0 | 0 | 140 |
| 1 | 577 | 106 | 79 | 0.129 | 0.065 | 144 |
| 1 | 578 | 106 | 79 | 0 | 0 | 139 |
| 1 | 579 | 106 | 79 | 0 | 0 | 159 |
| 1 | 580 | 106 | 79 | 0 | 0 | 153 |
| 1 | 581 | 103 | 75 | 0.645 | 0.323 | 163 |
| 1 | 582 | 103 | 75 | 0 | 0 | 160 |
| 1 | 583 | 103 | 75 | 0 | 0 | 160 |
| 1 | 584 | 103 | 75 | 0 | 0 | 155 |
| 1 | 585 | 102 | 75 | 0.129 | 0.065 | 157 |
| 1 | 586 | 102 | 75 | 0 | 0 | 154 |
| 1 | 587 | 102 | 75 | 0 | 0 | 144 |
| 1 | 588 | 104 | 70 | 0.695 | 0.347 | 158 |
| 1 | 589 | 104 | 70 | 0 | 0 | 165 |
| 1 | 590 | 104 | 70 | 0 | 0 | 169 |
| 1 | 591 | 104 | 70 | 0 | 0 | 172 |
| 1 | 592 | 104 | 70 | 0 | 0 | 167 |
| 1 | 593 | 104 | 70 | 0 | 0 | 169 |
| 1 | 594 | 104 | 71 | 0.129 | 0.065 | 155 |
| 1 | 595 | 104 | 71 | 0 | 0 | 164 |
| 1 | 596 | 104 | 71 | 0 | 0 | 162 |
| 1 | 597 | 104 | 71 | 0 | 0 | 163 |
| 1 | 598 | 104 | 74 | 0.387 | 0.194 | 142 |
| 1 | 599 | 104 | 74 | 0 | 0 | 146 |
| 1 | 600 | 104 | 74 | 0 | 0 | 160 |
| 1 | 601 | 104 | 74 | 0 | 0 | 165 |
| 1 | 602 | 104 | 74 | 0 | 0 | 160 |
| 1 | 603 | 104 | 74 | 0 | 0 | 157 |
| 1 | 604 | 104 | 74 | 0 | 0 | 160 |
| 1 | 605 | 101 | 80 | 0.865 | 0.433 | 147 |
| 1 | 606 | 101 | 80 | 0 | 0 | 148 |
| 1 | 607 | 102 | 80 | 0.129 | 0.065 | 145 |
| 1 | 608 | 102 | 80 | 0 | 0 | 136 |
| 1 | 609 | 102 | 80 | 0 | 0 | 145 |

| | | | | | | |
|---|---|---|---|---|---|---|
| 1 | 610 | 102 | 80 | 0 | 0 | 144 |
| 1 | 611 | 103 | 76 | 0.532 | 0.266 | 158 |
| 1 | 612 | 103 | 76 | 0 | 0 | 145 |
| 1 | 613 | 103 | 76 | 0 | 0 | 152 |
| 1 | 614 | 102 | 76 | 0.129 | 0.065 | 153 |
| 1 | 615 | 101 | 78 | 0.288 | 0.144 | 145 |
| 1 | 616 | 100 | 74 | 0.532 | 0.266 | 166 |
| 1 | 617 | 100 | 74 | 0 | 0 | 165 |
| 1 | 618 | 100 | 74 | 0 | 0 | 164 |
| 1 | 619 | 100 | 74 | 0 | 0 | 165 |
| 1 | 620 | 100 | 74 | 0 | 0 | 163 |
| 1 | 621 | 100 | 74 | 0 | 0 | 166 |
| 1 | 622 | 100 | 74 | 0 | 0 | 163 |
| 1 | 623 | 100 | 74 | 0 | 0 | 166 |
| 1 | 624 | 100 | 75 | 0.129 | 0.065 | 162 |
| 1 | 625 | 100 | 75 | 0 | 0 | 163 |
| 1 | 626 | 100 | 75 | 0 | 0 | 162 |
| 1 | 627 | 100 | 75 | 0 | 0 | 161 |
| 1 | 628 | 100 | 75 | 0 | 0 | 158 |
| 1 | 629 | 101 | 75 | 0.129 | 0.065 | 151 |
| 1 | 630 | 100 | 75 | 0.129 | 0.065 | 163 |
| 1 | 631 | 100 | 75 | 0 | 0 | 163 |
| 1 | 632 | 102 | 75 | 0.258 | 0.129 | 154 |
| 1 | 633 | 103 | 73 | 0.288 | 0.144 | 161 |
| 1 | 634 | 103 | 73 | 0 | 0 | 161 |
| 1 | 635 | 103 | 73 | 0 | 0 | 162 |
| 1 | 636 | 102 | 74 | 0.182 | 0.091 | 164 |
| 1 | 637 | 102 | 74 | 0 | 0 | 156 |
| 1 | 638 | 102 | 74 | 0 | 0 | 156 |
| 1 | 639 | 102 | 75 | 0.129 | 0.065 | 155 |
| 1 | 640 | 102 | 75 | 0 | 0 | 145 |
| 1 | 641 | 102 | 75 | 0 | 0 | 142 |
| 1 | 642 | 102 | 75 | 0 | 0 | 144 |
| 1 | 643 | 102 | 75 | 0 | 0 | 140 |
| 1 | 644 | 102 | 75 | 0 | 0 | 142 |
| 1 | 645 | 102 | 75 | 0 | 0 | 141 |
| 1 | 646 | 102 | 75 | 0 | 0 | 143 |
| 1 | 647 | 102 | 75 | 0 | 0 | 141 |
| 1 | 648 | 102 | 75 | 0 | 0 | 138 |
| 1 | 649 | 102 | 75 | 0 | 0 | 144 |
| 1 | 650 | 102 | 75 | 0 | 0 | 141 |
| 1 | 651 | 102 | 75 | 0 | 0 | 143 |
| 1 | 652 | 105 | 67 | 1.102 | 0.551 | 167 |
| 1 | 653 | 105 | 67 | 0 | 0 | 165 |
| 1 | 654 | 103 | 72 | 0.695 | 0.347 | 158 |
| 1 | 655 | 104 | 70 | 0.288 | 0.144 | 155 |
| 1 | 656 | 104 | 70 | 0 | 0 | 139 |

| | | | | | | |
|---|---|---|---|---|---|---|
| 1 | 657 | 104 | 70 | 0 | 0 | 161 |
| 1 | 658 | 104 | 70 | 0 | 0 | 144 |
| 1 | 659 | 104 | 70 | 0 | 0 | 132 |
| 1 | 660 | 104 | 70 | 0 | 0 | 132 |
| 1 | 661 | 104 | 70 | 0 | 0 | 134 |
| 1 | 662 | 104 | 70 | 0 | 0 | 132 |
| 1 | 663 | 104 | 69 | 0.129 | 0.065 | 136 |
| 1 | 664 | 104 | 69 | 0 | 0 | 151 |
| 1 | 665 | 104 | 69 | 0 | 0 | 148 |
| 1 | 666 | 103 | 69 | 0.129 | 0.065 | 143 |
| 1 | 667 | 102 | 69 | 0.129 | 0.065 | 151 |
| 1 | 668 | 102 | 69 | 0 | 0 | 164 |
| 1 | 669 | 100 | 71 | 0.365 | 0.182 | 162 |
| 1 | 670 | 100 | 71 | 0 | 0 | 164 |
| 1 | 671 | 100 | 71 | 0 | 0 | 162 |
| 1 | 672 | 100 | 70 | 0.129 | 0.065 | 160 |
| 1 | 673 | 100 | 69 | 0.129 | 0.065 | 161 |
| 1 | 674 | 100 | 69 | 0 | 0 | 161 |
| 1 | 675 | 100 | 69 | 0 | 0 | 162 |
| 1 | 676 | 100 | 69 | 0 | 0 | 164 |
| 1 | 677 | 100 | 69 | 0 | 0 | 163 |
| 1 | 678 | 100 | 69 | 0 | 0 | 162 |
| 1 | 679 | 100 | 69 | 0 | 0 | 162 |
| 1 | 680 | 100 | 69 | 0 | 0 | 163 |
| 1 | 681 | 100 | 69 | 0 | 0 | 161 |
| 1 | 682 | 100 | 69 | 0 | 0 | 165 |
| 1 | 683 | 100 | 69 | 0 | 0 | 166 |
| 1 | 684 | 100 | 69 | 0 | 0 | 163 |
| 1 | 685 | 100 | 69 | 0 | 0 | 165 |
| 1 | 686 | 100 | 69 | 0 | 0 | 163 |
| 1 | 687 | 100 | 69 | 0 | 0 | 161 |
| 1 | 688 | 99 | 69 | 0.129 | 0.065 | 163 |
| 1 | 689 | 99 | 69 | 0 | 0 | 159 |
| 1 | 690 | 99 | 69 | 0 | 0 | 159 |
| 1 | 691 | 99 | 69 | 0 | 0 | 160 |
| 1 | 692 | 99 | 69 | 0 | 0 | 160 |
| 1 | 693 | 99 | 69 | 0 | 0 | 161 |
| 1 | 694 | 99 | 69 | 0 | 0 | 158 |
| 1 | 695 | 99 | 69 | 0 | 0 | 160 |
| 1 | 696 | 99 | 69 | 0 | 0 | 161 |
| 1 | 697 | 99 | 69 | 0 | 0 | 163 |
| 1 | 698 | 99 | 69 | 0 | 0 | 164 |
| 1 | 699 | 99 | 68 | 0.129 | 0.065 | 163 |
| 1 | 700 | 99 | 68 | 0 | 0 | 163 |
| 1 | 701 | 99 | 68 | 0 | 0 | 163 |
| 1 | 702 | 99 | 68 | 0 | 0 | 164 |
| 1 | 703 | 99 | 68 | 0 | 0 | 165 |

| | | | | | | |
|---|---|---|---|---|---|---|
| 1 | 704 | 99 | 68 | 0 | 0 | 163 |
| 1 | 705 | 99 | 68 | 0 | 0 | 164 |
| 1 | 706 | 99 | 68 | 0 | 0 | 163 |
| 1 | 707 | 99 | 68 | 0 | 0 | 164 |
| 1 | 708 | 99 | 68 | 0 | 0 | 163 |
| 1 | 709 | 99 | 68 | 0 | 0 | 164 |
| 1 | 710 | 96 | 70 | 0.465 | 0.233 | 161 |
| 1 | 711 | 96 | 70 | 0 | 0 | 161 |
| 1 | 712 | 96 | 70 | 0 | 0 | 161 |
| 1 | 713 | 96 | 70 | 0 | 0 | 161 |
| 1 | 714 | 96 | 70 | 0 | 0 | 161 |
| 1 | 715 | 96 | 70 | 0 | 0 | 161 |
| 1 | 716 | 96 | 70 | 0 | 0 | 161 |
| 1 | 717 | 96 | 70 | 0 | 0 | 161 |
| 1 | 718 | 96 | 70 | 0 | 0 | 159 |
| 1 | 719 | 96 | 71 | 0.129 | 0.065 | 160 |
| 1 | 720 | 96 | 71 | 0 | 0 | 163 |
| 1 | 721 | 96 | 71 | 0 | 0 | 161 |
| 1 | 722 | 96 | 71 | 0 | 0 | 160 |
| 1 | 723 | 96 | 71 | 0 | 0 | 154 |
| 1 | 724 | 96 | 71 | 0 | 0 | 159 |
| 1 | 725 | 96 | 71 | 0 | 0 | 159 |
| 1 | 726 | 96 | 71 | 0 | 0 | 160 |
| 1 | 727 | 96 | 71 | 0 | 0 | 161 |
| 1 | 728 | 96 | 71 | 0 | 0 | 161 |
| 1 | 729 | 96 | 71 | 0 | 0 | 163 |
| 1 | 730 | 96 | 71 | 0 | 0 | 161 |
| 1 | 731 | 96 | 71 | 0 | 0 | 163 |
| 1 | 732 | 96 | 71 | 0 | 0 | 161 |
| 1 | 733 | 96 | 71 | 0 | 0 | 164 |
| 1 | 734 | 96 | 71 | 0 | 0 | 161 |
| 1 | 735 | 99 | 73 | 0.465 | 0.233 | 142 |
| 1 | 736 | 99 | 73 | 0 | 0 | 151 |
| 1 | 737 | 99 | 73 | 0 | 0 | 158 |
| 1 | 738 | 99 | 73 | 0 | 0 | 165 |
| 1 | 739 | 101 | 72 | 0.288 | 0.144 | 153 |
| 1 | 740 | 101 | 72 | 0 | 0 | 152 |
| 1 | 741 | 101 | 72 | 0 | 0 | 156 |
| 1 | 742 | 101 | 72 | 0 | 0 | 151 |
| 1 | 743 | 101 | 72 | 0 | 0 | 156 |
| 1 | 744 | 103 | 69 | 0.465 | 0.233 | 156 |
| 1 | 745 | 103 | 69 | 0 | 0 | 157 |
| 1 | 746 | 103 | 69 | 0 | 0 | 160 |
| 1 | 747 | 103 | 69 | 0 | 0 | 156 |
| 1 | 748 | 103 | 70 | 0.129 | 0.065 | 151 |
| 1 | 749 | 103 | 70 | 0 | 0 | 142 |
| 1 | 750 | 103 | 70 | 0 | 0 | 151 |

| | | | | | | |
|---|---|---|---|---|---|---|
| 1 | 751 | 103 | 70 | 0 | 0 | 142 |
| 1 | 752 | 103 | 70 | 0 | 0 | 145 |
| 1 | 753 | 103 | 70 | 0 | 0 | 145 |
| 1 | 754 | 103 | 70 | 0 | 0 | 149 |
| 1 | 755 | 103 | 70 | 0 | 0 | 153 |
| 1 | 756 | 103 | 70 | 0 | 0 | 152 |
| 1 | 757 | 104 | 71 | 0.182 | 0.091 | 144 |
| 1 | 758 | 104 | 71 | 0 | 0 | 145 |
| 1 | 759 | 104 | 71 | 0 | 0 | 150 |
| 1 | 760 | 109 | 68 | 0.752 | 0.376 | 153 |
| 1 | 761 | 109 | 69 | 0.129 | 0.065 | 159 |
| 1 | 762 | 109 | 69 | 0 | 0 | 156 |
| 1 | 763 | 109 | 69 | 0 | 0 | 150 |
| 1 | 764 | 107 | 69 | 0.258 | 0.129 | 141 |
| 1 | 765 | 105 | 66 | 0.465 | 0.233 | 163 |
| 1 | 766 | 105 | 66 | 0 | 0 | 161 |
| 1 | 767 | 104 | 73 | 0.912 | 0.456 | 172 |
| 1 | 768 | 106 | 69 | 0.577 | 0.288 | 155 |
| 1 | 769 | 106 | 69 | 0 | 0 | 156 |
| 1 | 770 | 106 | 69 | 0 | 0 | 152 |
| 1 | 771 | 106 | 69 | 0 | 0 | 148 |
| 1 | 772 | 106 | 69 | 0 | 0 | 149 |
| 1 | 773 | 106 | 69 | 0 | 0 | 153 |
| 1 | 774 | 106 | 69 | 0 | 0 | 148 |
| 1 | 775 | 104 | 68 | 0.288 | 0.144 | 158 |
| 1 | 776 | 104 | 68 | 0 | 0 | 160 |
| 1 | 777 | 104 | 68 | 0 | 0 | 160 |
| 1 | 778 | 106 | 64 | 0.577 | 0.288 | 153 |
| 1 | 779 | 106 | 64 | 0 | 0 | 156 |
| 1 | 780 | 106 | 64 | 0 | 0 | 152 |
| 1 | 781 | 106 | 64 | 0 | 0 | 161 |
| 1 | 782 | 106 | 64 | 0 | 0 | 160 |
| 1 | 783 | 106 | 64 | 0 | 0 | 163 |
| 1 | 784 | 106 | 64 | 0 | 0 | 152 |
| 1 | 785 | 106 | 64 | 0 | 0 | 148 |
| 1 | 786 | 106 | 64 | 0 | 0 | 149 |
| 1 | 787 | 106 | 64 | 0 | 0 | 149 |
| 1 | 788 | 106 | 64 | 0 | 0 | 145 |
| 1 | 789 | 106 | 63 | 0.129 | 0.065 | 152 |
| 1 | 790 | 106 | 63 | 0 | 0 | 163 |
| 1 | 791 | 106 | 63 | 0 | 0 | 156 |
| 1 | 792 | 107 | 62 | 0.182 | 0.091 | 167 |
| 1 | 793 | 109 | 62 | 0.258 | 0.129 | 145 |
| 1 | 794 | 109 | 62 | 0 | 0 | 140 |
| 1 | 795 | 109 | 62 | 0 | 0 | 137 |
| 1 | 796 | 109 | 62 | 0 | 0 | 138 |
| 1 | 797 | 109 | 62 | 0 | 0 | 142 |

| | | | | | | |
|---|---|---|---|---|---|---|
| 1 | 798 | 109 | 62 | 0 | 0 | 141 |
| 1 | 799 | 109 | 62 | 0 | 0 | 142 |
| 1 | 800 | 109 | 62 | 0 | 0 | 147 |
| 1 | 801 | 109 | 62 | 0 | 0 | 154 |
| 1 | 802 | 109 | 62 | 0 | 0 | 161 |
| 1 | 803 | 109 | 62 | 0 | 0 | 156 |
| 1 | 804 | 109 | 62 | 0 | 0 | 140 |
| 1 | 805 | 109 | 62 | 0 | 0 | 142 |
| 1 | 806 | 109 | 62 | 0 | 0 | 140 |
| 1 | 807 | 109 | 62 | 0 | 0 | 139 |
| 1 | 808 | 109 | 62 | 0 | 0 | 138 |
| 1 | 809 | 109 | 62 | 0 | 0 | 138 |
| 1 | 810 | 109 | 62 | 0 | 0 | 134 |
| 1 | 811 | 109 | 62 | 0 | 0 | 138 |
| 1 | 812 | 109 | 62 | 0 | 0 | 139 |
| 1 | 813 | 109 | 62 | 0 | 0 | 139 |
| 1 | 814 | 106 | 60 | 0.465 | 0.233 | 162 |
| 1 | 815 | 106 | 60 | 0 | 0 | 163 |
| 1 | 816 | 106 | 60 | 0 | 0 | 164 |
| 1 | 817 | 106 | 60 | 0 | 0 | 166 |
| 1 | 818 | 106 | 60 | 0 | 0 | 165 |
| 1 | 819 | 106 | 60 | 0 | 0 | 163 |
| 1 | 820 | 106 | 60 | 0 | 0 | 167 |
| 1 | 821 | 106 | 60 | 0 | 0 | 165 |
| 1 | 822 | 106 | 60 | 0 | 0 | 164 |
| 1 | 823 | 106 | 60 | 0 | 0 | 163 |
| 1 | 824 | 106 | 60 | 0 | 0 | 162 |
| 1 | 825 | 106 | 60 | 0 | 0 | 164 |
| 1 | 826 | 106 | 60 | 0 | 0 | 168 |
| 1 | 827 | 106 | 60 | 0 | 0 | 166 |
| 1 | 828 | 106 | 60 | 0 | 0 | 167 |
| 1 | 829 | 106 | 60 | 0 | 0 | 166 |
| 1 | 830 | 106 | 60 | 0 | 0 | 163 |
| 1 | 831 | 106 | 60 | 0 | 0 | 161 |
| 1 | 832 | 106 | 60 | 0 | 0 | 162 |
| 1 | 833 | 106 | 58 | 0.258 | 0.129 | 163 |
| 1 | 834 | 106 | 57 | 0.129 | 0.065 | 162 |
| 1 | 835 | 106 | 57 | 0 | 0 | 163 |
| 1 | 836 | 106 | 57 | 0 | 0 | 164 |
| 1 | 837 | 107 | 56 | 0.182 | 0.091 | 164 |
| 1 | 838 | 107 | 56 | 0 | 0 | 164 |
| 1 | 839 | 107 | 56 | 0 | 0 | 164 |
| 1 | 840 | 107 | 56 | 0 | 0 | 165 |
| 1 | 841 | 108 | 54 | 0.288 | 0.144 | 162 |
| 1 | 842 | 108 | 54 | 0 | 0 | 161 |
| 1 | 843 | 108 | 54 | 0 | 0 | 162 |
| 1 | 844 | 108 | 54 | 0 | 0 | 160 |

| | | | | | | |
|---|---|---|---|---|---|---|
| 1 | 845 | 109 | 53 | 0.182 | 0.091 | 159 |
| 1 | 846 | 109 | 53 | 0 | 0 | 162 |
| 1 | 847 | 109 | 56 | 0.387 | 0.194 | 148 |
| 1 | 848 | 109 | 56 | 0 | 0 | 153 |
| 1 | 849 | 109 | 56 | 0 | 0 | 161 |
| 1 | 850 | 108 | 59 | 0.408 | 0.204 | 141 |
| 1 | 851 | 108 | 59 | 0 | 0 | 138 |
| 1 | 852 | 108 | 59 | 0 | 0 | 139 |
| 1 | 853 | 107 | 59 | 0.129 | 0.065 | 139 |
| 1 | 854 | 107 | 59 | 0 | 0 | 138 |
| 1 | 855 | 107 | 59 | 0 | 0 | 148 |
| 1 | 856 | 107 | 59 | 0 | 0 | 148 |
| 1 | 857 | 107 | 57 | 0.258 | 0.129 | 139 |
| 1 | 858 | 107 | 56 | 0.129 | 0.065 | 141 |
| 1 | 859 | 107 | 56 | 0 | 0 | 148 |
| 1 | 860 | 107 | 56 | 0 | 0 | 147 |
| 1 | 861 | 107 | 56 | 0 | 0 | 141 |
| 1 | 862 | 107 | 56 | 0 | 0 | 146 |
| 1 | 863 | 107 | 56 | 0 | 0 | 148 |
| 1 | 864 | 107 | 56 | 0 | 0 | 144 |
| 1 | 865 | 107 | 56 | 0 | 0 | 155 |
| 1 | 866 | 107 | 56 | 0 | 0 | 153 |
| 1 | 867 | 107 | 56 | 0 | 0 | 148 |
| 1 | 868 | 107 | 56 | 0 | 0 | 155 |
| 1 | 869 | 107 | 56 | 0 | 0 | 144 |
| 1 | 870 | 107 | 56 | 0 | 0 | 145 |
| 1 | 871 | 107 | 56 | 0 | 0 | 144 |
| 1 | 872 | 107 | 56 | 0 | 0 | 147 |
| 1 | 873 | 106 | 56 | 0.129 | 0.065 | 156 |
| 1 | 874 | 106 | 56 | 0 | 0 | 156 |
| 1 | 875 | 106 | 53 | 0.387 | 0.194 | 160 |
| 1 | 876 | 106 | 53 | 0 | 0 | 165 |
| 1 | 877 | 106 | 53 | 0 | 0 | 158 |
| 1 | 878 | 106 | 53 | 0 | 0 | 160 |
| 1 | 879 | 106 | 53 | 0 | 0 | 162 |
| 1 | 880 | 106 | 53 | 0 | 0 | 163 |
| 1 | 881 | 106 | 53 | 0 | 0 | 160 |
| 1 | 882 | 106 | 53 | 0 | 0 | 161 |
| 1 | 883 | 106 | 53 | 0 | 0 | 160 |
| 1 | 884 | 106 | 53 | 0 | 0 | 160 |
| 1 | 885 | 106 | 53 | 0 | 0 | 161 |
| 1 | 886 | 106 | 53 | 0 | 0 | 162 |
| 1 | 887 | 106 | 53 | 0 | 0 | 161 |
| 1 | 888 | 106 | 53 | 0 | 0 | 162 |
| 1 | 889 | 106 | 53 | 0 | 0 | 165 |
| 1 | 890 | 106 | 53 | 0 | 0 | 158 |
| 1 | 891 | 107 | 52 | 0.182 | 0.091 | 153 |

| | | | | | | |
|---|---|---|---|---|---|---|
| 1 | 892 | 107 | 51 | 0.129 | 0.065 | 160 |
| 1 | 893 | 107 | 51 | 0 | 0 | 159 |
| 1 | 894 | 107 | 51 | 0 | 0 | 156 |
| 1 | 895 | 107 | 51 | 0 | 0 | 161 |
| 1 | 896 | 107 | 51 | 0 | 0 | 160 |
| 1 | 897 | 107 | 51 | 0 | 0 | 153 |
| 1 | 898 | 107 | 51 | 0 | 0 | 156 |
| 1 | 899 | 107 | 51 | 0 | 0 | 160 |
| 1 | 900 | 107 | 51 | 0 | 0 | 157 |
| 1 | 901 | 107 | 51 | 0 | 0 | 146 |
| 1 | 902 | 107 | 51 | 0 | 0 | 162 |
| 1 | 903 | 107 | 51 | 0 | 0 | 167 |
| 1 | 904 | 107 | 51 | 0 | 0 | 160 |
| 1 | 905 | 107 | 51 | 0 | 0 | 162 |
| 1 | 906 | 107 | 51 | 0 | 0 | 156 |
| 1 | 907 | 107 | 51 | 0 | 0 | 154 |
| 1 | 908 | 107 | 51 | 0 | 0 | 146 |
| 1 | 909 | 107 | 51 | 0 | 0 | 146 |
| 1 | 910 | 107 | 51 | 0 | 0 | 145 |
| 1 | 911 | 107 | 50 | 0.129 | 0.065 | 155 |
| 1 | 912 | 107 | 50 | 0 | 0 | 159 |
| 1 | 913 | 107 | 50 | 0 | 0 | 154 |
| 1 | 914 | 107 | 50 | 0 | 0 | 159 |
| 1 | 915 | 107 | 50 | 0 | 0 | 162 |
| 1 | 916 | 107 | 50 | 0 | 0 | 154 |
| 1 | 917 | 107 | 50 | 0 | 0 | 156 |
| 1 | 918 | 107 | 50 | 0 | 0 | 162 |
| 1 | 919 | 107 | 50 | 0 | 0 | 161 |
| 1 | 920 | 107 | 50 | 0 | 0 | 165 |
| 1 | 921 | 107 | 50 | 0 | 0 | 160 |
| 1 | 922 | 107 | 50 | 0 | 0 | 142 |
| 1 | 923 | 106 | 50 | 0.129 | 0.065 | 151 |
| 1 | 924 | 105 | 50 | 0.129 | 0.065 | 158 |
| 1 | 925 | 105 | 50 | 0 | 0 | 158 |
| 1 | 926 | 105 | 50 | 0 | 0 | 156 |
| 1 | 927 | 105 | 50 | 0 | 0 | 156 |
| 1 | 928 | 105 | 50 | 0 | 0 | 147 |
| 1 | 929 | 105 | 49 | 0.129 | 0.065 | 158 |
| 1 | 930 | 105 | 49 | 0 | 0 | 156 |
| 1 | 931 | 105 | 49 | 0 | 0 | 154 |
| 1 | 932 | 105 | 49 | 0 | 0 | 158 |
| 1 | 933 | 105 | 49 | 0 | 0 | 154 |
| 1 | 934 | 105 | 49 | 0 | 0 | 154 |
| 1 | 935 | 105 | 47 | 0.258 | 0.129 | 159 |
| 1 | 936 | 105 | 47 | 0 | 0 | 158 |
| 1 | 937 | 105 | 47 | 0 | 0 | 160 |
| 1 | 938 | 105 | 46 | 0.129 | 0.065 | 159 |

| | | | | | | |
|---|---|---|---|---|---|---|
| 1 | 939 | 105 | 46 | 0 | 0 | 160 |
| 1 | 940 | 105 | 46 | 0 | 0 | 160 |
| 1 | 941 | 106 | 49 | 0.408 | 0.204 | 148 |
| 1 | 942 | 107 | 48 | 0.182 | 0.091 | 145 |
| 1 | 943 | 107 | 48 | 0 | 0 | 149 |
| 1 | 944 | 107 | 48 | 0 | 0 | 148 |
| 1 | 945 | 107 | 48 | 0 | 0 | 136 |
| 1 | 946 | 107 | 48 | 0 | 0 | 136 |
| 1 | 947 | 107 | 48 | 0 | 0 | 141 |
| 1 | 948 | 107 | 48 | 0 | 0 | 144 |
| 1 | 949 | 107 | 48 | 0 | 0 | 138 |
| 1 | 950 | 107 | 47 | 0.129 | 0.065 | 144 |
| 1 | 951 | 107 | 47 | 0 | 0 | 141 |
| 1 | 952 | 105 | 47 | 0.258 | 0.129 | 160 |
| 1 | 953 | 105 | 47 | 0 | 0 | 154 |
| 1 | 954 | 105 | 47 | 0 | 0 | 150 |
| 1 | 955 | 105 | 47 | 0 | 0 | 154 |
| 1 | 956 | 105 | 47 | 0 | 0 | 151 |
| 1 | 957 | 105 | 47 | 0 | 0 | 148 |
| 1 | 958 | 104 | 47 | 0.129 | 0.065 | 155 |
| 1 | 959 | 104 | 45 | 0.258 | 0.129 | 161 |
| 1 | 960 | 104 | 45 | 0 | 0 | 161 |
| 1 | 961 | 104 | 45 | 0 | 0 | 160 |
| 1 | 962 | 104 | 45 | 0 | 0 | 146 |
| 1 | 963 | 104 | 45 | 0 | 0 | 147 |
| 1 | 964 | 104 | 45 | 0 | 0 | 155 |
| 1 | 965 | 102 | 44 | 0.288 | 0.144 | 159 |
| 1 | 966 | 102 | 44 | 0 | 0 | 159 |
| 1 | 967 | 102 | 44 | 0 | 0 | 160 |
| 1 | 968 | 102 | 44 | 0 | 0 | 159 |
| 1 | 969 | 102 | 44 | 0 | 0 | 159 |
| 1 | 970 | 102 | 44 | 0 | 0 | 160 |
| 1 | 971 | 102 | 44 | 0 | 0 | 160 |
| 1 | 972 | 102 | 44 | 0 | 0 | 160 |
| 1 | 973 | 102 | 42 | 0.258 | 0.129 | 162 |
| 1 | 974 | 106 | 45 | 0.645 | 0.323 | 153 |
| 1 | 975 | 106 | 45 | 0 | 0 | 146 |
| 1 | 976 | 106 | 45 | 0 | 0 | 145 |
| 1 | 977 | 106 | 45 | 0 | 0 | 143 |
| 1 | 978 | 106 | 44 | 0.129 | 0.065 | 149 |
| 1 | 979 | 106 | 44 | 0 | 0 | 151 |
| 1 | 980 | 106 | 44 | 0 | 0 | 143 |
| 1 | 981 | 106 | 44 | 0 | 0 | 145 |
| 1 | 982 | 106 | 44 | 0 | 0 | 145 |
| 1 | 983 | 106 | 44 | 0 | 0 | 140 |
| 1 | 984 | 106 | 44 | 0 | 0 | 140 |
| 1 | 985 | 106 | 44 | 0 | 0 | 140 |

| | | | | | | |
|---|---|---|---|---|---|---|
| 1 | 986 | 106 | 44 | 0 | 0 | 139 |
| 1 | 987 | 106 | 43 | 0.129 | 0.065 | 146 |
| 1 | 988 | 106 | 43 | 0 | 0 | 141 |
| 1 | 989 | 106 | 43 | 0 | 0 | 142 |
| 1 | 990 | 106 | 43 | 0 | 0 | 146 |
| 1 | 991 | 106 | 43 | 0 | 0 | 154 |
| 1 | 992 | 106 | 43 | 0 | 0 | 154 |
| 1 | 993 | 105 | 42 | 0.182 | 0.091 | 160 |
| 1 | 994 | 105 | 42 | 0 | 0 | 162 |
| 1 | 995 | 105 | 42 | 0 | 0 | 161 |
| 1 | 996 | 105 | 42 | 0 | 0 | 162 |
| 1 | 997 | 105 | 42 | 0 | 0 | 159 |
| 1 | 998 | 105 | 42 | 0 | 0 | 162 |
| 1 | 999 | 105 | 42 | 0 | 0 | 160 |
| 1 | 1000 | 105 | 42 | 0 | 0 | 147 |
| 1 | 1001 | 106 | 40 | 0.288 | 0.144 | 161 |
| 1 | 1002 | 106 | 40 | 0 | 0 | 165 |
| 1 | 1003 | 106 | 40 | 0 | 0 | 166 |
| 1 | 1004 | 106 | 40 | 0 | 0 | 157 |
| 1 | 1005 | 106 | 40 | 0 | 0 | 161 |
| 1 | 1006 | 106 | 40 | 0 | 0 | 161 |
| 1 | 1007 | 106 | 40 | 0 | 0 | 151 |
| 1 | 1008 | 106 | 40 | 0 | 0 | 167 |
| 1 | 1009 | 106 | 40 | 0 | 0 | 166 |
| 1 | 1010 | 106 | 40 | 0 | 0 | 139 |
| 1 | 1011 | 106 | 40 | 0 | 0 | 143 |
| 1 | 1012 | 106 | 40 | 0 | 0 | 161 |
| 1 | 1013 | 106 | 40 | 0 | 0 | 153 |
| 1 | 1014 | 106 | 40 | 0 | 0 | 134 |
| 1 | 1015 | 106 | 40 | 0 | 0 | 161 |
| 1 | 1016 | 106 | 40 | 0 | 0 | 163 |
| 1 | 1017 | 106 | 40 | 0 | 0 | 163 |
| 1 | 1018 | 102 | 43 | 0.645 | 0.323 | 160 |
| 1 | 1019 | 102 | 44 | 0.129 | 0.065 | 159 |
| 1 | 1020 | 102 | 46 | 0.258 | 0.129 | 161 |
| 1 | 1021 | 102 | 46 | 0 | 0 | 166 |
| 1 | 1022 | 102 | 42 | 0.516 | 0.258 | 160 |
| 1 | 1023 | 102 | 42 | 0 | 0 | 161 |
| 1 | 1024 | 102 | 42 | 0 | 0 | 163 |
| 1 | 1025 | 102 | 42 | 0 | 0 | 161 |
| 1 | 1026 | 102 | 42 | 0 | 0 | 162 |
| 1 | 1027 | 102 | 42 | 0 | 0 | 162 |
| 1 | 1028 | 102 | 42 | 0 | 0 | 162 |
| 1 | 1029 | 102 | 42 | 0 | 0 | 162 |
| 1 | 1030 | 102 | 42 | 0 | 0 | 162 |
| 1 | 1031 | 102 | 42 | 0 | 0 | 167 |
| 1 | 1032 | 102 | 41 | 0.129 | 0.065 | 165 |

| | | | | | | |
|---|---|---|---|---|---|---|
| 1 | 1033 | 102 | 41 | 0 | 0 | 166 |
| 1 | 1034 | 102 | 40 | 0.129 | 0.065 | 163 |
| 1 | 1035 | 98 | 37 | 0.645 | 0.323 | 162 |
| 1 | 1036 | 98 | 37 | 0 | 0 | 162 |
| 1 | 1037 | 98 | 37 | 0 | 0 | 162 |
| 1 | 1038 | 98 | 37 | 0 | 0 | 162 |
| 1 | 1039 | 101 | 41 | 0.645 | 0.323 | 165 |
| 1 | 1040 | 102 | 41 | 0.129 | 0.065 | 165 |
| 1 | 1041 | 102 | 40 | 0.129 | 0.065 | 163 |
| 1 | 1042 | 102 | 39 | 0.129 | 0.065 | 161 |
| 1 | 1043 | 100 | 39 | 0.258 | 0.129 | 162 |
| 1 | 1044 | 100 | 39 | 0 | 0 | 160 |
| 1 | 1045 | 99 | 39 | 0.129 | 0.065 | 160 |
| 1 | 1046 | 99 | 39 | 0 | 0 | 161 |
| 1 | 1047 | 99 | 39 | 0 | 0 | 161 |
| 1 | 1048 | 99 | 39 | 0 | 0 | 161 |
| 1 | 1049 | 99 | 39 | 0 | 0 | 161 |
| 1 | 1050 | 99 | 39 | 0 | 0 | 162 |
| 1 | 1051 | 99 | 39 | 0 | 0 | 164 |
| 1 | 1052 | 99 | 39 | 0 | 0 | 164 |
| 1 | 1053 | 98 | 39 | 0.129 | 0.065 | 161 |
| 1 | 1054 | 98 | 39 | 0 | 0 | 160 |
| 1 | 1055 | 98 | 39 | 0 | 0 | 161 |
| 1 | 1056 | 98 | 39 | 0 | 0 | 161 |
| 1 | 1057 | 98 | 39 | 0 | 0 | 161 |
| 1 | 1058 | 98 | 39 | 0 | 0 | 161 |
| 1 | 1059 | 98 | 39 | 0 | 0 | 161 |
| 1 | 1060 | 98 | 39 | 0 | 0 | 163 |
| 1 | 1061 | 98 | 39 | 0 | 0 | 161 |
| 1 | 1062 | 98 | 39 | 0 | 0 | 163 |
| 1 | 1063 | 98 | 39 | 0 | 0 | 161 |
| 1 | 1064 | 98 | 39 | 0 | 0 | 161 |
| 1 | 1065 | 98 | 39 | 0 | 0 | 163 |
| 1 | 1066 | 99 | 39 | 0.129 | 0.065 | 164 |
| 1 | 1067 | 99 | 39 | 0 | 0 | 165 |
| 1 | 1068 | 99 | 39 | 0 | 0 | 165 |
| 1 | 1069 | 99 | 38 | 0.129 | 0.065 | 165 |
| 1 | 1070 | 99 | 38 | 0 | 0 | 163 |
| 1 | 1071 | 99 | 38 | 0 | 0 | 167 |
| 1 | 1072 | 98 | 38 | 0.129 | 0.065 | 162 |
| 1 | 1073 | 98 | 38 | 0 | 0 | 160 |
| 1 | 1074 | 98 | 38 | 0 | 0 | 163 |
| 1 | 1075 | 98 | 38 | 0 | 0 | 165 |
| 1 | 1076 | 98 | 38 | 0 | 0 | 166 |
| 1 | 1077 | 98 | 38 | 0 | 0 | 163 |
| 1 | 1078 | 98 | 38 | 0 | 0 | 166 |
| 1 | 1079 | 98 | 38 | 0 | 0 | 168 |

| | | | | | | |
|---|---|---|---|---|---|---|
| 1 | 1080 | 98 | 38 | 0 | 0 | 168 |
| 1 | 1081 | 98 | 38 | 0 | 0 | 163 |
| 1 | 1082 | 97 | 36 | 0.288 | 0.144 | 160 |
| 1 | 1083 | 97 | 36 | 0 | 0 | 160 |
| 1 | 1084 | 97 | 36 | 0 | 0 | 160 |
| 1 | 1085 | 97 | 36 | 0 | 0 | 160 |
| 1 | 1086 | 96 | 36 | 0.129 | 0.065 | 159 |
| 1 | 1087 | 98 | 42 | 0.816 | 0.408 | 165 |
| 1 | 1088 | 98 | 42 | 0 | 0 | 165 |
| 1 | 1089 | 98 | 42 | 0 | 0 | 156 |
| 1 | 1090 | 98 | 42 | 0 | 0 | 152 |
| 1 | 1091 | 97 | 40 | 0.288 | 0.144 | 162 |
| 1 | 1092 | 96 | 40 | 0.129 | 0.065 | 162 |
| 1 | 1093 | 96 | 40 | 0 | 0 | 162 |
| 1 | 1094 | 96 | 40 | 0 | 0 | 161 |
| 1 | 1095 | 96 | 40 | 0 | 0 | 162 |
| 1 | 1096 | 96 | 40 | 0 | 0 | 163 |
| 1 | 1097 | 96 | 40 | 0 | 0 | 165 |
| 1 | 1098 | 96 | 40 | 0 | 0 | 167 |
| 1 | 1099 | 96 | 40 | 0 | 0 | 166 |
| 1 | 1100 | 96 | 40 | 0 | 0 | 170 |
| 1 | 1101 | 96 | 40 | 0 | 0 | 172 |
| 1 | 1102 | 96 | 40 | 0 | 0 | 165 |
| 1 | 1103 | 96 | 40 | 0 | 0 | 167 |
| 1 | 1104 | 96 | 40 | 0 | 0 | 168 |
| 1 | 1105 | 96 | 40 | 0 | 0 | 163 |
| 1 | 1106 | 96 | 40 | 0 | 0 | 162 |
| 1 | 1107 | 96 | 40 | 0 | 0 | 162 |
| 1 | 1108 | 96 | 40 | 0 | 0 | 165 |
| 1 | 1109 | 96 | 40 | 0 | 0 | 165 |
| 1 | 1110 | 96 | 40 | 0 | 0 | 171 |
| 1 | 1111 | 96 | 40 | 0 | 0 | 166 |
| 1 | 1112 | 96 | 40 | 0 | 0 | 166 |
| 1 | 1113 | 96 | 40 | 0 | 0 | 164 |
| 1 | 1114 | 95 | 40 | 0.129 | 0.065 | 163 |
| 1 | 1115 | 94 | 40 | 0.129 | 0.065 | 165 |
| 1 | 1116 | 94 | 40 | 0 | 0 | 164 |
| 1 | 1117 | 94 | 40 | 0 | 0 | 161 |
| 1 | 1118 | 94 | 40 | 0 | 0 | 163 |
| 1 | 1119 | 94 | 40 | 0 | 0 | 165 |
| 1 | 1120 | 94 | 40 | 0 | 0 | 145 |
| 1 | 1121 | 94 | 40 | 0 | 0 | 142 |
| 1 | 1122 | 94 | 40 | 0 | 0 | 152 |
| 1 | 1123 | 94 | 40 | 0 | 0 | 166 |
| 1 | 1124 | 94 | 40 | 0 | 0 | 156 |
| 1 | 1125 | 94 | 40 | 0 | 0 | 143 |
| 1 | 1126 | 94 | 40 | 0 | 0 | 148 |

| | | | | | | |
|---|---|---|---|---|---|---|
| 1 | 1127 | 94 | 40 | 0 | 0 | 146 |
| 1 | 1128 | 94 | 40 | 0 | 0 | 140 |
| 1 | 1129 | 94 | 40 | 0 | 0 | 139 |
| 1 | 1130 | 94 | 40 | 0 | 0 | 141 |
| 1 | 1131 | 94 | 40 | 0 | 0 | 138 |
| 1 | 1132 | 94 | 40 | 0 | 0 | 136 |
| 1 | 1133 | 93 | 35 | 0.658 | 0.329 | 166 |
| 1 | 1134 | 93 | 35 | 0 | 0 | 165 |
| 1 | 1135 | 92 | 35 | 0.129 | 0.065 | 166 |
| 1 | 1136 | 92 | 35 | 0 | 0 | 166 |
| 1 | 1137 | 91 | 35 | 0.129 | 0.065 | 166 |
| 1 | 1138 | 91 | 34 | 0.129 | 0.065 | 164 |
| 1 | 1139 | 91 | 34 | 0 | 0 | 166 |
| 1 | 1140 | 91 | 34 | 0 | 0 | 165 |
| 1 | 1141 | 91 | 34 | 0 | 0 | 166 |
| 1 | 1142 | 91 | 34 | 0 | 0 | 165 |
| 1 | 1143 | 91 | 34 | 0 | 0 | 163 |
| 1 | 1144 | 91 | 34 | 0 | 0 | 162 |
| 1 | 1145 | 91 | 34 | 0 | 0 | 162 |
| 1 | 1146 | 91 | 34 | 0 | 0 | 162 |
| 1 | 1147 | 91 | 34 | 0 | 0 | 162 |
| 1 | 1148 | 91 | 34 | 0 | 0 | 163 |
| 1 | 1149 | 91 | 34 | 0 | 0 | 163 |
| 1 | 1150 | 96 | 39 | 0.912 | 0.456 | 139 |
| 1 | 1151 | 95 | 39 | 0.129 | 0.065 | 139 |
| 1 | 1152 | 95 | 39 | 0 | 0 | 141 |
| 1 | 1153 | 95 | 39 | 0 | 0 | 139 |
| 1 | 1154 | 95 | 39 | 0 | 0 | 141 |
| 1 | 1155 | 95 | 39 | 0 | 0 | 139 |
| 1 | 1156 | 95 | 39 | 0 | 0 | 139 |
| 1 | 1157 | 95 | 39 | 0 | 0 | 131 |
| 1 | 1158 | 95 | 39 | 0 | 0 | 145 |
| 1 | 1159 | 94 | 39 | 0.129 | 0.065 | 145 |
| 1 | 1160 | 94 | 39 | 0 | 0 | 147 |
| 1 | 1161 | 94 | 39 | 0 | 0 | 154 |
| 1 | 1162 | 94 | 39 | 0 | 0 | 149 |
| 1 | 1163 | 93 | 39 | 0.129 | 0.065 | 155 |
| 1 | 1164 | 93 | 39 | 0 | 0 | 148 |
| 1 | 1165 | 93 | 39 | 0 | 0 | 149 |
| 1 | 1166 | 93 | 39 | 0 | 0 | 156 |
| 1 | 1167 | 93 | 39 | 0 | 0 | 163 |
| 1 | 1168 | 92 | 38 | 0.182 | 0.091 | 164 |
| 1 | 1169 | 92 | 38 | 0 | 0 | 169 |
| 1 | 1170 | 92 | 38 | 0 | 0 | 168 |
| 1 | 1171 | 92 | 38 | 0 | 0 | 165 |
| 1 | 1172 | 92 | 38 | 0 | 0 | 164 |
| 1 | 1173 | 92 | 38 | 0 | 0 | 165 |

| | | | | | | |
|---|---|---|---|---|---|---|
| 1 | 1174 | 92 | 38 | 0 | 0 | 163 |
| 1 | 1175 | 92 | 38 | 0 | 0 | 158 |
| 1 | 1176 | 92 | 38 | 0 | 0 | 152 |
| 1 | 1177 | 92 | 38 | 0 | 0 | 158 |
| 1 | 1178 | 92 | 38 | 0 | 0 | 155 |
| 1 | 1179 | 92 | 38 | 0 | 0 | 149 |
| 1 | 1180 | 92 | 38 | 0 | 0 | 140 |
| 1 | 1181 | 92 | 38 | 0 | 0 | 138 |
| 1 | 1182 | 92 | 38 | 0 | 0 | 142 |
| 1 | 1183 | 92 | 38 | 0 | 0 | 139 |
| 1 | 1184 | 90 | 33 | 0.695 | 0.347 | 166 |
| 1 | 1185 | 90 | 33 | 0 | 0 | 164 |
| 1 | 1186 | 90 | 33 | 0 | 0 | 166 |
| 1 | 1187 | 90 | 33 | 0 | 0 | 170 |
| 1 | 1188 | 88 | 33 | 0.258 | 0.129 | 162 |
| 1 | 1189 | 88 | 33 | 0 | 0 | 161 |
| 1 | 1190 | 88 | 33 | 0 | 0 | 161 |
| 1 | 1191 | 88 | 33 | 0 | 0 | 161 |
| 1 | 1192 | 91 | 33 | 0.387 | 0.194 | 156 |
| 1 | 1193 | 91 | 33 | 0 | 0 | 159 |
| 1 | 1194 | 91 | 33 | 0 | 0 | 163 |
| 1 | 1195 | 91 | 32 | 0.129 | 0.065 | 168 |
| 1 | 1196 | 91 | 32 | 0 | 0 | 169 |
| 1 | 1197 | 91 | 32 | 0 | 0 | 169 |
| 1 | 1198 | 91 | 32 | 0 | 0 | 170 |
| 1 | 1199 | 91 | 32 | 0 | 0 | 165 |
| 1 | 1200 | 91 | 32 | 0 | 0 | 163 |
| 1 | 1201 | 91 | 32 | 0 | 0 | 166 |
| 1 | 1202 | 91 | 32 | 0 | 0 | 165 |
| 1 | 1203 | 91 | 32 | 0 | 0 | 165 |
| 1 | 1204 | 93 | 29 | 0.465 | 0.233 | 162 |
| 1 | 1205 | 93 | 29 | 0 | 0 | 169 |
| 1 | 1206 | 93 | 29 | 0 | 0 | 159 |
| 1 | 1207 | 94 | 28 | 0.182 | 0.091 | 162 |
| 1 | 1208 | 94 | 28 | 0 | 0 | 166 |
| 1 | 1209 | 94 | 28 | 0 | 0 | 164 |
| 1 | 1210 | 94 | 28 | 0 | 0 | 169 |
| 1 | 1211 | 94 | 28 | 0 | 0 | 163 |
| 1 | 1212 | 94 | 28 | 0 | 0 | 164 |
| 1 | 1213 | 94 | 28 | 0 | 0 | 164 |
| 1 | 1214 | 94 | 28 | 0 | 0 | 167 |
| 1 | 1215 | 94 | 28 | 0 | 0 | 165 |
| 1 | 1216 | 93 | 28 | 0.129 | 0.065 | 164 |
| 1 | 1217 | 93 | 32 | 0.516 | 0.258 | 170 |
| 1 | 1218 | 93 | 32 | 0 | 0 | 168 |
| 1 | 1219 | 93 | 32 | 0 | 0 | 167 |
| 1 | 1220 | 94 | 31 | 0.182 | 0.091 | 156 |

| | | | | | | |
|---|---|---|---|---|---|---|
| 1 | 1221 | 95 | 30 | 0.182 | 0.091 | 146 |
| 1 | 1222 | 95 | 30 | 0 | 0 | 164 |
| 1 | 1223 | 95 | 29 | 0.129 | 0.065 | 168 |
| 1 | 1224 | 95 | 29 | 0 | 0 | 156 |
| 1 | 1225 | 94 | 29 | 0.129 | 0.065 | 165 |
| 1 | 1226 | 93 | 29 | 0.129 | 0.065 | 163 |
| 1 | 1227 | 93 | 29 | 0 | 0 | 165 |
| 1 | 1228 | 93 | 29 | 0 | 0 | 163 |
| 1 | 1229 | 93 | 25 | 0.516 | 0.258 | 159 |
| 1 | 1230 | 96 | 31 | 0.865 | 0.433 | 149 |
| 1 | 1231 | 96 | 31 | 0 | 0 | 156 |
| 1 | 1232 | 96 | 29 | 0.258 | 0.129 | 163 |
| 1 | 1233 | 96 | 29 | 0 | 0 | 163 |
| 1 | 1234 | 96 | 28 | 0.129 | 0.065 | 165 |
| 1 | 1235 | 97 | 27 | 0.182 | 0.091 | 163 |
| 1 | 1236 | 97 | 27 | 0 | 0 | 163 |
| 1 | 1237 | 97 | 31 | 0.516 | 0.258 | 147 |
| 1 | 1238 | 97 | 31 | 0 | 0 | 144 |
| 1 | 1239 | 97 | 31 | 0 | 0 | 139 |
| 1 | 1240 | 97 | 32 | 0.129 | 0.065 | 137 |
| 1 | 1241 | 97 | 32 | 0 | 0 | 143 |
| 1 | 1242 | 97 | 32 | 0 | 0 | 139 |
| 1 | 1243 | 97 | 32 | 0 | 0 | 144 |
| 1 | 1244 | 97 | 32 | 0 | 0 | 146 |
| 1 | 1245 | 95 | 26 | 0.816 | 0.408 | 168 |
| 1 | 1246 | 95 | 26 | 0 | 0 | 165 |
| 1 | 1247 | 95 | 26 | 0 | 0 | 165 |
| 1 | 1248 | 95 | 26 | 0 | 0 | 163 |
| 1 | 1249 | 94 | 25 | 0.182 | 0.091 | 159 |
| 1 | 1250 | 94 | 26 | 0.129 | 0.065 | 165 |
| 1 | 1251 | 94 | 26 | 0 | 0 | 166 |
| 1 | 1252 | 93 | 26 | 0.129 | 0.065 | 162 |
| 1 | 1253 | 90 | 26 | 0.387 | 0.194 | 158 |
| 1 | 1254 | 88 | 26 | 0.258 | 0.129 | 158 |
| 1 | 1255 | 95 | 31 | 1.11 | 0.555 | 137 |
| 1 | 1256 | 95 | 31 | 0 | 0 | 140 |
| 1 | 1257 | 95 | 31 | 0 | 0 | 146 |
| 1 | 1258 | 95 | 31 | 0 | 0 | 163 |
| 1 | 1259 | 95 | 31 | 0 | 0 | 148 |
| 1 | 1260 | 95 | 31 | 0 | 0 | 143 |
| 1 | 1261 | 95 | 30 | 0.129 | 0.065 | 156 |
| 1 | 1262 | 95 | 30 | 0 | 0 | 154 |
| 1 | 1263 | 95 | 30 | 0 | 0 | 150 |
| 1 | 1264 | 94 | 30 | 0.129 | 0.065 | 163 |
| 1 | 1265 | 93 | 31 | 0.182 | 0.091 | 164 |
| 1 | 1266 | 93 | 31 | 0 | 0 | 163 |
| 1 | 1267 | 93 | 32 | 0.129 | 0.065 | 161 |

| | | | | | | |
|---|---|---|---|---|---|---|
| 1 | 1268 | 93 | 32 | 0 | 0 | 162 |
| 1 | 1269 | 93 | 33 | 0.129 | 0.065 | 162 |
| 1 | 1270 | 93 | 33 | 0 | 0 | 155 |
| 1 | 1271 | 93 | 33 | 0 | 0 | 157 |
| 1 | 1272 | 93 | 33 | 0 | 0 | 159 |
| 1 | 1273 | 93 | 33 | 0 | 0 | 156 |
| 1 | 1274 | 93 | 33 | 0 | 0 | 158 |
| 1 | 1275 | 93 | 33 | 0 | 0 | 165 |
| 1 | 1276 | 93 | 33 | 0 | 0 | 163 |
| 1 | 1277 | 93 | 33 | 0 | 0 | 165 |
| 1 | 1278 | 93 | 33 | 0 | 0 | 164 |
| 1 | 1279 | 97 | 31 | 0.577 | 0.288 | 166 |
| 1 | 1280 | 97 | 31 | 0 | 0 | 161 |
| 1 | 1281 | 97 | 31 | 0 | 0 | 157 |
| 1 | 1282 | 97 | 31 | 0 | 0 | 159 |
| 1 | 1283 | 97 | 31 | 0 | 0 | 143 |
| 1 | 1284 | 97 | 31 | 0 | 0 | 142 |
| 1 | 1285 | 97 | 31 | 0 | 0 | 167 |
| 1 | 1286 | 97 | 31 | 0 | 0 | 154 |
| 1 | 1287 | 97 | 31 | 0 | 0 | 145 |
| 1 | 1288 | 97 | 30 | 0.129 | 0.065 | 157 |
| 1 | 1289 | 97 | 30 | 0 | 0 | 154 |
| 1 | 1290 | 97 | 30 | 0 | 0 | 159 |
| 1 | 1291 | 97 | 30 | 0 | 0 | 161 |
| 1 | 1292 | 97 | 30 | 0 | 0 | 158 |
| 1 | 1293 | 97 | 29 | 0.129 | 0.065 | 152 |
| 1 | 1294 | 97 | 29 | 0 | 0 | 141 |
| 1 | 1295 | 96 | 28 | 0.182 | 0.091 | 168 |
| 1 | 1296 | 95 | 32 | 0.532 | 0.266 | 141 |
| 1 | 1297 | 95 | 32 | 0 | 0 | 147 |
| 1 | 1298 | 95 | 32 | 0 | 0 | 147 |
| 1 | 1299 | 95 | 32 | 0 | 0 | 160 |
| 1 | 1300 | 95 | 32 | 0 | 0 | 162 |
| 1 | 1301 | 95 | 33 | 0.129 | 0.065 | 157 |
| 1 | 1302 | 95 | 34 | 0.129 | 0.065 | 140 |
| 1 | 1303 | 95 | 34 | 0 | 0 | 146 |
| 1 | 1304 | 94 | 34 | 0.129 | 0.065 | 152 |
| 1 | 1305 | 93 | 32 | 0.288 | 0.144 | 164 |
| 1 | 1306 | 93 | 32 | 0 | 0 | 163 |
| 1 | 1307 | 93 | 32 | 0 | 0 | 162 |
| 1 | 1308 | 93 | 32 | 0 | 0 | 165 |
| 1 | 1309 | 94 | 31 | 0.182 | 0.091 | 165 |
| 1 | 1310 | 94 | 31 | 0 | 0 | 163 |
| 1 | 1311 | 94 | 31 | 0 | 0 | 161 |
| 1 | 1312 | 94 | 31 | 0 | 0 | 161 |
| 1 | 1313 | 94 | 31 | 0 | 0 | 162 |
| 1 | 1314 | 94 | 31 | 0 | 0 | 149 |

| | | | | | | |
|---|---|---|---|---|---|---|
| 1 | 1315 | 94 | 31 | 0 | 0 | 146 |
| 1 | 1316 | 94 | 31 | 0 | 0 | 150 |
| 1 | 1317 | 95 | 25 | 0.785 | 0.392 | 163 |
| 1 | 1318 | 95 | 25 | 0 | 0 | 163 |
| 1 | 1319 | 95 | 25 | 0 | 0 | 161 |
| 1 | 1320 | 95 | 25 | 0 | 0 | 161 |
| 1 | 1321 | 95 | 28 | 0.387 | 0.194 | 162 |
| 1 | 1322 | 95 | 28 | 0 | 0 | 169 |
| 1 | 1323 | 95 | 28 | 0 | 0 | 161 |
| 1 | 1324 | 95 | 28 | 0 | 0 | 143 |
| 1 | 1325 | 95 | 28 | 0 | 0 | 146 |
| 1 | 1326 | 95 | 28 | 0 | 0 | 155 |
| 1 | 1327 | 95 | 28 | 0 | 0 | 164 |
| 1 | 1328 | 95 | 28 | 0 | 0 | 161 |
| 1 | 1329 | 95 | 28 | 0 | 0 | 162 |
| 1 | 1330 | 95 | 28 | 0 | 0 | 163 |
| 1 | 1331 | 95 | 28 | 0 | 0 | 163 |
| 1 | 1332 | 95 | 28 | 0 | 0 | 164 |
| 1 | 1333 | 95 | 28 | 0 | 0 | 164 |
| 1 | 1334 | 95 | 28 | 0 | 0 | 163 |
| 1 | 1335 | 95 | 28 | 0 | 0 | 163 |
| 1 | 1336 | 95 | 28 | 0 | 0 | 149 |
| 1 | 1337 | 95 | 28 | 0 | 0 | 165 |
| 1 | 1338 | 95 | 28 | 0 | 0 | 164 |
| 1 | 1339 | 95 | 28 | 0 | 0 | 164 |
| 1 | 1340 | 95 | 28 | 0 | 0 | 166 |
| 1 | 1341 | 95 | 28 | 0 | 0 | 164 |
| 1 | 1342 | 94 | 29 | 0.182 | 0.091 | 163 |
| 1 | 1343 | 94 | 28 | 0.129 | 0.065 | 164 |
| 1 | 1344 | 94 | 28 | 0 | 0 | 160 |
| 1 | 1345 | 94 | 28 | 0 | 0 | 161 |
| 1 | 1346 | 94 | 28 | 0 | 0 | 167 |
| 1 | 1347 | 94 | 28 | 0 | 0 | 168 |
| 1 | 1348 | 94 | 28 | 0 | 0 | 167 |
| 1 | 1349 | 94 | 28 | 0 | 0 | 167 |
| 1 | 1350 | 94 | 28 | 0 | 0 | 162 |
| 1 | 1351 | 94 | 28 | 0 | 0 | 163 |
| 1 | 1352 | 94 | 28 | 0 | 0 | 164 |
| 1 | 1353 | 94 | 28 | 0 | 0 | 163 |
| 1 | 1354 | 94 | 29 | 0.129 | 0.065 | 165 |
| 1 | 1355 | 93 | 30 | 0.182 | 0.091 | 158 |
| 1 | 1356 | 93 | 30 | 0 | 0 | 161 |
| 1 | 1357 | 92 | 30 | 0.129 | 0.065 | 163 |
| 1 | 1358 | 92 | 30 | 0 | 0 | 163 |
| 1 | 1359 | 92 | 29 | 0.129 | 0.065 | 163 |
| 1 | 1360 | 92 | 29 | 0 | 0 | 163 |
| 1 | 1361 | 92 | 29 | 0 | 0 | 163 |

| | | | | | | |
|---|---|---|---|---|---|---|
| 1 | 1362 | 92 | 29 | 0 | 0 | 162 |
| 1 | 1363 | 92 | 28 | 0.129 | 0.065 | 163 |
| 1 | 1364 | 92 | 27 | 0.129 | 0.065 | 161 |
| 1 | 1365 | 92 | 27 | 0 | 0 | 162 |
| 1 | 1366 | 92 | 27 | 0 | 0 | 161 |
| 1 | 1367 | 92 | 27 | 0 | 0 | 160 |
| 1 | 1368 | 92 | 27 | 0 | 0 | 160 |
| 1 | 1369 | 92 | 27 | 0 | 0 | 163 |
| 1 | 1370 | 92 | 27 | 0 | 0 | 163 |
| 1 | 1371 | 92 | 27 | 0 | 0 | 161 |
| 1 | 1372 | 92 | 27 | 0 | 0 | 162 |
| 1 | 1373 | 92 | 27 | 0 | 0 | 160 |
| 1 | 1374 | 92 | 27 | 0 | 0 | 160 |
| 1 | 1375 | 92 | 27 | 0 | 0 | 159 |
| 1 | 1376 | 92 | 27 | 0 | 0 | 159 |
| 1 | 1377 | 92 | 27 | 0 | 0 | 159 |
| 1 | 1378 | 92 | 27 | 0 | 0 | 158 |
| 1 | 1379 | 92 | 27 | 0 | 0 | 158 |
| 1 | 1380 | 92 | 26 | 0.129 | 0.065 | 157 |
| 1 | 1381 | 92 | 26 | 0 | 0 | 159 |
| 1 | 1382 | 92 | 26 | 0 | 0 | 158 |
| 1 | 1383 | 92 | 26 | 0 | 0 | 158 |
| 1 | 1384 | 92 | 26 | 0 | 0 | 158 |
| 1 | 1385 | 92 | 26 | 0 | 0 | 158 |
| 1 | 1386 | 92 | 26 | 0 | 0 | 158 |
| 1 | 1387 | 92 | 26 | 0 | 0 | 158 |
| 1 | 1388 | 92 | 26 | 0 | 0 | 157 |
| 1 | 1389 | 92 | 26 | 0 | 0 | 157 |
| 1 | 1390 | 92 | 26 | 0 | 0 | 159 |
| 1 | 1391 | 92 | 26 | 0 | 0 | 159 |
| 1 | 1392 | 92 | 26 | 0 | 0 | 159 |
| 1 | 1393 | 92 | 26 | 0 | 0 | 159 |
| 1 | 1394 | 92 | 26 | 0 | 0 | 159 |
| 1 | 1395 | 92 | 26 | 0 | 0 | 160 |
| 1 | 1396 | 92 | 26 | 0 | 0 | 160 |
| 1 | 1397 | 92 | 26 | 0 | 0 | 160 |
| 1 | 1398 | 92 | 26 | 0 | 0 | 159 |
| 1 | 1399 | 92 | 26 | 0 | 0 | 158 |
| 1 | 1400 | 92 | 26 | 0 | 0 | 160 |
| 1 | 1401 | 92 | 26 | 0 | 0 | 160 |
| 1 | 1402 | 92 | 26 | 0 | 0 | 160 |
| 1 | 1403 | 92 | 26 | 0 | 0 | 160 |
| 1 | 1404 | 91 | 31 | 0.658 | 0.329 | 161 |
| 1 | 1405 | 89 | 32 | 0.288 | 0.144 | 161 |
| 1 | 1406 | 89 | 32 | 0 | 0 | 162 |
| 1 | 1407 | 89 | 32 | 0 | 0 | 163 |
| 1 | 1408 | 89 | 32 | 0 | 0 | 161 |

| | | | | | | |
|---|---|---|---|---|---|---|
| 1 | 1409 | 89 | 32 | 0 | 0 | 162 |
| 1 | 1410 | 89 | 32 | 0 | 0 | 163 |
| 1 | 1411 | 89 | 32 | 0 | 0 | 162 |
| 1 | 1412 | 89 | 32 | 0 | 0 | 161 |
| 1 | 1413 | 89 | 32 | 0 | 0 | 159 |
| 1 | 1414 | 89 | 32 | 0 | 0 | 160 |
| 1 | 1415 | 89 | 32 | 0 | 0 | 160 |
| 1 | 1416 | 89 | 32 | 0 | 0 | 162 |
| 1 | 1417 | 89 | 32 | 0 | 0 | 160 |
| 1 | 1418 | 89 | 32 | 0 | 0 | 162 |
| 1 | 1419 | 89 | 32 | 0 | 0 | 162 |
| 1 | 1420 | 89 | 32 | 0 | 0 | 162 |
| 1 | 1421 | 88 | 32 | 0.129 | 0.065 | 162 |
| 1 | 1422 | 88 | 32 | 0 | 0 | 162 |
| 1 | 1423 | 88 | 32 | 0 | 0 | 162 |
| 1 | 1424 | 88 | 32 | 0 | 0 | 162 |
| 1 | 1425 | 88 | 32 | 0 | 0 | 163 |
| 1 | 1426 | 88 | 32 | 0 | 0 | 161 |
| 1 | 1427 | 88 | 32 | 0 | 0 | 161 |
| 1 | 1428 | 88 | 32 | 0 | 0 | 163 |
| 1 | 1429 | 88 | 32 | 0 | 0 | 162 |
| 1 | 1430 | 88 | 32 | 0 | 0 | 161 |
| 1 | 1431 | 88 | 32 | 0 | 0 | 161 |
| 1 | 1432 | 88 | 32 | 0 | 0 | 160 |
| 1 | 1433 | 88 | 32 | 0 | 0 | 161 |
| 1 | 1434 | 88 | 32 | 0 | 0 | 160 |
| 1 | 1435 | 88 | 32 | 0 | 0 | 159 |
| 1 | 1436 | 88 | 32 | 0 | 0 | 160 |
| 1 | 1437 | 88 | 32 | 0 | 0 | 160 |
| 1 | 1438 | 88 | 32 | 0 | 0 | 160 |
| 1 | 1439 | 88 | 32 | 0 | 0 | 160 |
| 1 | 1440 | 88 | 32 | 0 | 0 | 160 |
| 1 | 1441 | 88 | 32 | 0 | 0 | 160 |
| 1 | 1442 | 91 | 38 | 0.865 | 0.433 | 140 |
| 1 | 1443 | 91 | 38 | 0 | 0 | 138 |
| 1 | 1444 | 91 | 38 | 0 | 0 | 140 |
| 1 | 1445 | 91 | 38 | 0 | 0 | 140 |
| 1 | 1446 | 91 | 38 | 0 | 0 | 145 |
| 1 | 1447 | 93 | 35 | 0.465 | 0.233 | 156 |
| 1 | 1448 | 93 | 36 | 0.129 | 0.065 | 142 |
| 1 | 1449 | 93 | 36 | 0 | 0 | 140 |
| 1 | 1450 | 93 | 36 | 0 | 0 | 145 |
| 1 | 1451 | 93 | 36 | 0 | 0 | 142 |
| 1 | 1452 | 93 | 36 | 0 | 0 | 144 |
| 1 | 1453 | 93 | 36 | 0 | 0 | 144 |
| 1 | 1454 | 93 | 36 | 0 | 0 | 145 |
| 1 | 1455 | 93 | 37 | 0.129 | 0.065 | 142 |

| | | | | | | |
|---|---|---|---|---|---|---|
| 1 | 1456 | 93 | 37 | 0 | 0 | 143 |
| 1 | 1457 | 93 | 37 | 0 | 0 | 142 |
| 1 | 1458 | 93 | 37 | 0 | 0 | 142 |
| 1 | 1459 | 93 | 37 | 0 | 0 | 140 |
| 1 | 1460 | 88 | 38 | 0.658 | 0.329 | 168 |
| 1 | 1461 | 88 | 38 | 0 | 0 | 166 |
| 1 | 1462 | 88 | 38 | 0 | 0 | 165 |
| 1 | 1463 | 90 | 38 | 0.258 | 0.129 | 154 |
| 1 | 1464 | 91 | 38 | 0.129 | 0.065 | 144 |
| 1 | 1465 | 91 | 37 | 0.129 | 0.065 | 149 |
| 1 | 1466 | 91 | 37 | 0 | 0 | 145 |
| 1 | 1467 | 91 | 37 | 0 | 0 | 146 |
| 1 | 1468 | 91 | 37 | 0 | 0 | 149 |
| 1 | 1469 | 90 | 34 | 0.408 | 0.204 | 161 |
| 1 | 1470 | 90 | 34 | 0 | 0 | 160 |
| 1 | 1471 | 90 | 34 | 0 | 0 | 161 |
| 1 | 1472 | 90 | 36 | 0.258 | 0.129 | 155 |
| 1 | 1473 | 90 | 36 | 0 | 0 | 153 |
| 1 | 1474 | 90 | 36 | 0 | 0 | 149 |
| 1 | 1475 | 90 | 36 | 0 | 0 | 144 |
| 1 | 1476 | 90 | 34 | 0.258 | 0.129 | 147 |
| 1 | 1477 | 90 | 34 | 0 | 0 | 151 |
| 1 | 1478 | 90 | 34 | 0 | 0 | 149 |
| 1 | 1479 | 89 | 36 | 0.288 | 0.144 | 144 |
| 1 | 1480 | 88 | 36 | 0.129 | 0.065 | 153 |
| 1 | 1481 | 87 | 36 | 0.129 | 0.065 | 165 |
| 1 | 1482 | 87 | 36 | 0 | 0 | 164 |
| 1 | 1483 | 88 | 36 | 0.129 | 0.065 | 152 |
| 1 | 1484 | 88 | 36 | 0 | 0 | 159 |
| 1 | 1485 | 88 | 36 | 0 | 0 | 147 |
| 1 | 1486 | 88 | 36 | 0 | 0 | 161 |
| 1 | 1487 | 88 | 36 | 0 | 0 | 161 |
| 1 | 1488 | 88 | 36 | 0 | 0 | 164 |
| 1 | 1489 | 88 | 36 | 0 | 0 | 167 |
| 1 | 1490 | 88 | 36 | 0 | 0 | 161 |
| 1 | 1491 | 88 | 36 | 0 | 0 | 160 |
| 1 | 1492 | 88 | 36 | 0 | 0 | 153 |
| 1 | 1493 | 87 | 36 | 0.129 | 0.065 | 158 |
| 1 | 1494 | 87 | 36 | 0 | 0 | 163 |
| 1 | 1495 | 87 | 36 | 0 | 0 | 162 |
| 1 | 1496 | 86 | 36 | 0.129 | 0.065 | 161 |
| 1 | 1497 | 86 | 36 | 0 | 0 | 162 |
| 1 | 1498 | 86 | 36 | 0 | 0 | 158 |
| 1 | 1499 | 86 | 36 | 0 | 0 | 162 |
| 1 | 1500 | 85 | 36 | 0.129 | 0.065 | 162 |
| 1 | 1501 | 85 | 36 | 0 | 0 | 163 |
| 1 | 1502 | 85 | 36 | 0 | 0 | 163 |

| | | | | | | |
|---|---|---|---|---|---|---|
| 1 | 1503 | 85 | 36 | 0 | 0 | 163 |
| 1 | 1504 | 85 | 36 | 0 | 0 | 165 |
| 1 | 1505 | 85 | 36 | 0 | 0 | 162 |
| 1 | 1506 | 85 | 36 | 0 | 0 | 165 |
| 1 | 1507 | 85 | 36 | 0 | 0 | 166 |
| 1 | 1508 | 85 | 36 | 0 | 0 | 161 |
| 1 | 1509 | 85 | 36 | 0 | 0 | 163 |
| 1 | 1510 | 84 | 36 | 0.129 | 0.065 | 161 |
| 1 | 1511 | 84 | 36 | 0 | 0 | 166 |
| 1 | 1512 | 84 | 36 | 0 | 0 | 165 |
| 1 | 1513 | 84 | 36 | 0 | 0 | 163 |
| 1 | 1514 | 84 | 36 | 0 | 0 | 163 |
| 1 | 1515 | 84 | 36 | 0 | 0 | 164 |
| 1 | 1516 | 84 | 36 | 0 | 0 | 161 |
| 1 | 1517 | 84 | 36 | 0 | 0 | 161 |
| 1 | 1518 | 84 | 36 | 0 | 0 | 163 |
| 1 | 1519 | 84 | 36 | 0 | 0 | 163 |
| 1 | 1520 | 84 | 36 | 0 | 0 | 163 |
| 1 | 1521 | 84 | 36 | 0 | 0 | 164 |
| 1 | 1522 | 84 | 36 | 0 | 0 | 163 |
| 1 | 1523 | 84 | 36 | 0 | 0 | 160 |
| 1 | 1524 | 84 | 43 | 0.903 | 0.452 | 166 |
| 1 | 1525 | 84 | 43 | 0 | 0 | 165 |
| 1 | 1526 | 84 | 43 | 0 | 0 | 170 |
| 1 | 1527 | 84 | 43 | 0 | 0 | 167 |
| 1 | 1528 | 84 | 43 | 0 | 0 | 167 |
| 1 | 1529 | 84 | 42 | 0.129 | 0.065 | 164 |
| 1 | 1530 | 84 | 42 | 0 | 0 | 161 |
| 1 | 1531 | 84 | 42 | 0 | 0 | 163 |
| 1 | 1532 | 84 | 42 | 0 | 0 | 165 |
| 1 | 1533 | 84 | 42 | 0 | 0 | 163 |
| 1 | 1534 | 84 | 42 | 0 | 0 | 162 |
| 1 | 1535 | 84 | 42 | 0 | 0 | 163 |
| 1 | 1536 | 84 | 42 | 0 | 0 | 165 |
| 1 | 1537 | 84 | 42 | 0 | 0 | 165 |
| 1 | 1538 | 83 | 43 | 0.182 | 0.091 | 166 |
| 1 | 1539 | 82 | 43 | 0.129 | 0.065 | 163 |
| 1 | 1540 | 81 | 43 | 0.129 | 0.065 | 163 |
| 1 | 1541 | 81 | 43 | 0 | 0 | 160 |
| 1 | 1542 | 81 | 43 | 0 | 0 | 159 |
| 1 | 1543 | 81 | 43 | 0 | 0 | 158 |
| 1 | 1544 | 81 | 43 | 0 | 0 | 160 |
| 1 | 1545 | 86 | 43 | 0.645 | 0.323 | 161 |
| 1 | 1546 | 86 | 43 | 0 | 0 | 158 |
| 1 | 1547 | 86 | 44 | 0.129 | 0.065 | 149 |
| 1 | 1548 | 85 | 46 | 0.288 | 0.144 | 164 |
| 1 | 1549 | 85 | 46 | 0 | 0 | 162 |

| | | | | | | |
|---|---|---|---|---|---|---|
| 1 | 1550 | 85 | 47 | 0.129 | 0.065 | 165 |
| 1 | 1551 | 85 | 47 | 0 | 0 | 163 |
| 1 | 1552 | 85 | 47 | 0 | 0 | 163 |
| 1 | 1553 | 85 | 47 | 0 | 0 | 162 |
| 1 | 1554 | 85 | 47 | 0 | 0 | 159 |
| 1 | 1555 | 85 | 47 | 0 | 0 | 155 |
| 1 | 1556 | 85 | 47 | 0 | 0 | 157 |
| 1 | 1557 | 85 | 47 | 0 | 0 | 153 |
| 1 | 1558 | 84 | 47 | 0.129 | 0.065 | 163 |
| 1 | 1559 | 84 | 47 | 0 | 0 | 163 |
| 1 | 1560 | 84 | 47 | 0 | 0 | 164 |
| 1 | 1561 | 84 | 47 | 0 | 0 | 166 |
| 1 | 1562 | 84 | 47 | 0 | 0 | 161 |
| 1 | 1563 | 83 | 47 | 0.129 | 0.065 | 171 |
| 1 | 1564 | 83 | 45 | 0.258 | 0.129 | 165 |
| 1 | 1565 | 83 | 45 | 0 | 0 | 168 |
| 1 | 1566 | 83 | 45 | 0 | 0 | 170 |
| 1 | 1567 | 83 | 44 | 0.129 | 0.065 | 166 |
| 1 | 1568 | 83 | 44 | 0 | 0 | 167 |
| 1 | 1569 | 83 | 44 | 0 | 0 | 172 |
| 1 | 1570 | 83 | 44 | 0 | 0 | 173 |
| 1 | 1571 | 83 | 44 | 0 | 0 | 169 |
| 1 | 1572 | 83 | 44 | 0 | 0 | 168 |
| 1 | 1573 | 83 | 44 | 0 | 0 | 169 |
| 1 | 1574 | 83 | 44 | 0 | 0 | 170 |
| 1 | 1575 | 83 | 44 | 0 | 0 | 170 |
| 1 | 1576 | 83 | 44 | 0 | 0 | 170 |
| 1 | 1577 | 83 | 44 | 0 | 0 | 169 |
| 1 | 1578 | 81 | 43 | 0.288 | 0.144 | 170 |
| 1 | 1579 | 81 | 43 | 0 | 0 | 168 |
| 1 | 1580 | 81 | 43 | 0 | 0 | 170 |
| 1 | 1581 | 81 | 43 | 0 | 0 | 169 |
| 1 | 1582 | 81 | 43 | 0 | 0 | 167 |
| 1 | 1583 | 81 | 43 | 0 | 0 | 170 |
| 1 | 1584 | 81 | 43 | 0 | 0 | 169 |
| 1 | 1585 | 81 | 43 | 0 | 0 | 170 |
| 1 | 1586 | 81 | 43 | 0 | 0 | 169 |
| 1 | 1587 | 81 | 43 | 0 | 0 | 167 |
| 1 | 1588 | 81 | 41 | 0.258 | 0.129 | 165 |
| 1 | 1589 | 81 | 41 | 0 | 0 | 165 |
| 1 | 1590 | 81 | 41 | 0 | 0 | 167 |
| 1 | 1591 | 81 | 41 | 0 | 0 | 167 |
| 1 | 1592 | 81 | 41 | 0 | 0 | 167 |
| 1 | 1593 | 81 | 41 | 0 | 0 | 156 |
| 1 | 1594 | 81 | 41 | 0 | 0 | 161 |
| 1 | 1595 | 81 | 41 | 0 | 0 | 157 |
| 1 | 1596 | 81 | 41 | 0 | 0 | 139 |

| | | | | | | |
|---|---|---|---|---|---|---|
| 1 | 1597 | 81 | 41 | 0 | 0 | 145 |
| 1 | 1598 | 81 | 40 | 0.129 | 0.065 | 145 |
| 1 | 1599 | 81 | 40 | 0 | 0 | 144 |
| 1 | 1600 | 81 | 40 | 0 | 0 | 145 |
| 1 | 1601 | 81 | 40 | 0 | 0 | 145 |
| 1 | 1602 | 81 | 40 | 0 | 0 | 152 |
| 1 | 1603 | 78 | 39 | 0.408 | 0.204 | 170 |
| 1 | 1604 | 78 | 39 | 0 | 0 | 167 |
| 1 | 1605 | 78 | 39 | 0 | 0 | 167 |
| 1 | 1606 | 78 | 39 | 0 | 0 | 167 |
| 1 | 1607 | 78 | 39 | 0 | 0 | 167 |
| 1 | 1608 | 77 | 40 | 0.182 | 0.091 | 166 |
| 1 | 1609 | 78 | 43 | 0.408 | 0.204 | 152 |
| 1 | 1610 | 78 | 43 | 0 | 0 | 155 |
| 1 | 1611 | 78 | 43 | 0 | 0 | 154 |
| 1 | 1612 | 78 | 43 | 0 | 0 | 145 |
| 1 | 1613 | 78 | 43 | 0 | 0 | 154 |
| 1 | 1614 | 78 | 43 | 0 | 0 | 149 |
| 1 | 1615 | 78 | 43 | 0 | 0 | 147 |
| 1 | 1616 | 78 | 43 | 0 | 0 | 149 |
| 1 | 1617 | 78 | 43 | 0 | 0 | 151 |
| 1 | 1618 | 78 | 43 | 0 | 0 | 146 |
| 1 | 1619 | 78 | 42 | 0.129 | 0.065 | 147 |
| 1 | 1620 | 78 | 42 | 0 | 0 | 149 |
| 1 | 1621 | 78 | 41 | 0.129 | 0.065 | 160 |
| 1 | 1622 | 78 | 41 | 0 | 0 | 163 |
| 1 | 1623 | 78 | 41 | 0 | 0 | 162 |
| 1 | 1624 | 78 | 41 | 0 | 0 | 155 |
| 1 | 1625 | 76 | 41 | 0.258 | 0.129 | 163 |
| 1 | 1626 | 76 | 41 | 0 | 0 | 162 |
| 1 | 1627 | 76 | 41 | 0 | 0 | 161 |
| 1 | 1628 | 76 | 41 | 0 | 0 | 161 |
| 1 | 1629 | 76 | 41 | 0 | 0 | 162 |
| 1 | 1630 | 76 | 41 | 0 | 0 | 162 |
| 1 | 1631 | 76 | 41 | 0 | 0 | 161 |
| 1 | 1632 | 76 | 41 | 0 | 0 | 158 |
| 1 | 1633 | 76 | 41 | 0 | 0 | 160 |
| 1 | 1634 | 76 | 41 | 0 | 0 | 162 |
| 1 | 1635 | 76 | 41 | 0 | 0 | 165 |
| 1 | 1636 | 76 | 41 | 0 | 0 | 161 |
| 1 | 1637 | 76 | 41 | 0 | 0 | 160 |
| 1 | 1638 | 73 | 49 | 1.102 | 0.551 | 145 |
| 1 | 1639 | 73 | 49 | 0 | 0 | 146 |
| 1 | 1640 | 73 | 49 | 0 | 0 | 148 |
| 1 | 1641 | 73 | 49 | 0 | 0 | 150 |
| 1 | 1642 | 73 | 49 | 0 | 0 | 155 |
| 1 | 1643 | 73 | 49 | 0 | 0 | 148 |

| | | | | | | |
|---|---|---|---|---|---|---|
| 1 | 1644 | 73 | 49 | 0 | 0 | 150 |
| 1 | 1645 | 73 | 49 | 0 | 0 | 142 |
| 1 | 1646 | 73 | 49 | 0 | 0 | 142 |
| 1 | 1647 | 73 | 49 | 0 | 0 | 143 |
| 1 | 1648 | 72 | 49 | 0.129 | 0.065 | 149 |
| 1 | 1649 | 72 | 49 | 0 | 0 | 147 |
| 1 | 1650 | 68 | 47 | 0.577 | 0.288 | 165 |
| 1 | 1651 | 68 | 47 | 0 | 0 | 161 |
| 1 | 1652 | 70 | 47 | 0.258 | 0.129 | 161 |
| 1 | 1653 | 70 | 47 | 0 | 0 | 163 |
| 1 | 1654 | 70 | 47 | 0 | 0 | 164 |
| 1 | 1655 | 70 | 47 | 0 | 0 | 161 |
| 1 | 1656 | 69 | 47 | 0.129 | 0.065 | 165 |
| 1 | 1657 | 69 | 47 | 0 | 0 | 166 |
| 1 | 1658 | 69 | 47 | 0 | 0 | 161 |
| 1 | 1659 | 69 | 47 | 0 | 0 | 159 |
| 1 | 1660 | 69 | 47 | 0 | 0 | 164 |
| 1 | 1661 | 69 | 47 | 0 | 0 | 165 |
| 1 | 1662 | 69 | 47 | 0 | 0 | 161 |
| 1 | 1663 | 69 | 47 | 0 | 0 | 155 |
| 1 | 1664 | 68 | 43 | 0.532 | 0.266 | 162 |
| 1 | 1665 | 71 | 44 | 0.408 | 0.204 | 159 |
| 1 | 1666 | 71 | 44 | 0 | 0 | 153 |
| 1 | 1667 | 71 | 44 | 0 | 0 | 154 |
| 1 | 1668 | 71 | 45 | 0.129 | 0.065 | 143 |
| 1 | 1669 | 71 | 45 | 0 | 0 | 143 |
| 1 | 1670 | 71 | 45 | 0 | 0 | 144 |
| 1 | 1671 | 71 | 45 | 0 | 0 | 144 |
| 1 | 1672 | 70 | 42 | 0.408 | 0.204 | 157 |
| 1 | 1673 | 70 | 42 | 0 | 0 | 157 |
| 1 | 1674 | 70 | 42 | 0 | 0 | 151 |
| 1 | 1675 | 70 | 42 | 0 | 0 | 142 |
| 1 | 1676 | 70 | 42 | 0 | 0 | 145 |
| 1 | 1677 | 69 | 40 | 0.288 | 0.144 | 154 |
| 1 | 1678 | 69 | 40 | 0 | 0 | 149 |
| 1 | 1679 | 69 | 40 | 0 | 0 | 153 |
| 1 | 1680 | 69 | 40 | 0 | 0 | 154 |
| 1 | 1681 | 69 | 40 | 0 | 0 | 147 |
| 1 | 1682 | 69 | 40 | 0 | 0 | 156 |
| 1 | 1683 | 68 | 46 | 0.785 | 0.392 | 159 |
| 1 | 1684 | 68 | 41 | 0.645 | 0.323 | 160 |
| 1 | 1685 | 68 | 41 | 0 | 0 | 158 |
| 1 | 1686 | 68 | 41 | 0 | 0 | 153 |
| 1 | 1687 | 68 | 41 | 0 | 0 | 156 |
| 1 | 1688 | 68 | 41 | 0 | 0 | 152 |
| 1 | 1689 | 68 | 40 | 0.129 | 0.065 | 158 |
| 1 | 1690 | 68 | 40 | 0 | 0 | 160 |

| | | | | | | |
|---|---|---|---|---|---|---|
| 1 | 1691 | 68 | 40 | 0 | 0 | 163 |
| 1 | 1692 | 68 | 40 | 0 | 0 | 154 |
| 1 | 1693 | 68 | 40 | 0 | 0 | 164 |
| 1 | 1694 | 68 | 40 | 0 | 0 | 160 |
| 1 | 1695 | 68 | 40 | 0 | 0 | 152 |
| 1 | 1696 | 68 | 40 | 0 | 0 | 149 |
| 1 | 1697 | 68 | 39 | 0.129 | 0.065 | 161 |
| 1 | 1698 | 67 | 39 | 0.129 | 0.065 | 158 |
| 1 | 1699 | 67 | 39 | 0 | 0 | 159 |
| 1 | 1700 | 67 | 39 | 0 | 0 | 163 |
| 1 | 1701 | 67 | 39 | 0 | 0 | 168 |
| 1 | 1702 | 67 | 39 | 0 | 0 | 169 |
| 1 | 1703 | 67 | 39 | 0 | 0 | 169 |
| 1 | 1704 | 67 | 38 | 0.129 | 0.065 | 161 |
| 1 | 1705 | 67 | 37 | 0.129 | 0.065 | 155 |
| 1 | 1706 | 67 | 37 | 0 | 0 | 165 |
| 1 | 1707 | 67 | 37 | 0 | 0 | 163 |
| 1 | 1708 | 67 | 37 | 0 | 0 | 166 |
| 1 | 1709 | 67 | 37 | 0 | 0 | 160 |
| 1 | 1710 | 67 | 37 | 0 | 0 | 164 |
| 1 | 1711 | 67 | 37 | 0 | 0 | 164 |
| 1 | 1712 | 67 | 37 | 0 | 0 | 164 |
| 1 | 1713 | 67 | 37 | 0 | 0 | 154 |
| 1 | 1714 | 67 | 37 | 0 | 0 | 152 |
| 1 | 1715 | 67 | 37 | 0 | 0 | 157 |
| 1 | 1716 | 67 | 37 | 0 | 0 | 157 |
| 1 | 1717 | 67 | 37 | 0 | 0 | 155 |
| 1 | 1718 | 67 | 37 | 0 | 0 | 154 |
| 1 | 1719 | 67 | 37 | 0 | 0 | 156 |
| 1 | 1720 | 67 | 37 | 0 | 0 | 161 |
| 1 | 1721 | 67 | 37 | 0 | 0 | 161 |
| 1 | 1722 | 67 | 37 | 0 | 0 | 157 |
| 1 | 1723 | 67 | 37 | 0 | 0 | 149 |
| 1 | 1724 | 67 | 37 | 0 | 0 | 137 |
| 1 | 1725 | 62 | 40 | 0.752 | 0.376 | 164 |
| 1 | 1726 | 64 | 41 | 0.288 | 0.144 | 166 |
| 1 | 1727 | 65 | 41 | 0.129 | 0.065 | 159 |
| 1 | 1728 | 65 | 40 | 0.129 | 0.065 | 146 |
| 1 | 1729 | 64 | 40 | 0.129 | 0.065 | 149 |
| 1 | 1730 | 64 | 40 | 0 | 0 | 166 |
| 1 | 1731 | 64 | 39 | 0.129 | 0.065 | 166 |
| 1 | 1732 | 64 | 39 | 0 | 0 | 159 |
| 1 | 1733 | 64 | 39 | 0 | 0 | 148 |
| 1 | 1734 | 62 | 37 | 0.365 | 0.182 | 166 |
| 1 | 1735 | 62 | 36 | 0.129 | 0.065 | 161 |
| 1 | 1736 | 62 | 36 | 0 | 0 | 163 |
| 1 | 1737 | 62 | 36 | 0 | 0 | 159 |

| | | | | | | |
|---|---|---|---|---|---|---|
| 1 | 1738 | 61 | 43 | 0.912 | 0.456 | 168 |
| 1 | 1739 | 61 | 42 | 0.129 | 0.065 | 168 |
| 1 | 1740 | 61 | 41 | 0.129 | 0.065 | 163 |
| 1 | 1741 | 61 | 41 | 0 | 0 | 163 |
| 1 | 1742 | 61 | 41 | 0 | 0 | 164 |
| 1 | 1743 | 60 | 40 | 0.182 | 0.091 | 166 |
| 1 | 1744 | 59 | 39 | 0.182 | 0.091 | 162 |
| 1 | 1745 | 59 | 39 | 0 | 0 | 162 |
| 1 | 1746 | 60 | 40 | 0.182 | 0.091 | 163 |
| 1 | 1747 | 60 | 40 | 0 | 0 | 162 |
| 1 | 1748 | 60 | 40 | 0 | 0 | 163 |
| 1 | 1749 | 60 | 40 | 0 | 0 | 162 |
| 1 | 1750 | 60 | 40 | 0 | 0 | 164 |
| 1 | 1751 | 60 | 40 | 0 | 0 | 164 |
| 1 | 1752 | 60 | 38 | 0.258 | 0.129 | 160 |
| 1 | 1753 | 60 | 38 | 0 | 0 | 163 |
| 1 | 1754 | 60 | 38 | 0 | 0 | 150 |
| 1 | 1755 | 58 | 43 | 0.695 | 0.347 | 156 |
| 1 | 1756 | 58 | 43 | 0 | 0 | 161 |
| 1 | 1757 | 58 | 43 | 0 | 0 | 167 |
| 1 | 1758 | 58 | 43 | 0 | 0 | 160 |
| 1 | 1759 | 58 | 43 | 0 | 0 | 144 |
| 1 | 1760 | 58 | 43 | 0 | 0 | 144 |
| 1 | 1761 | 58 | 43 | 0 | 0 | 143 |
| 1 | 1762 | 58 | 43 | 0 | 0 | 148 |
| 1 | 1763 | 58 | 43 | 0 | 0 | 141 |
| 1 | 1764 | 58 | 43 | 0 | 0 | 149 |
| 1 | 1765 | 58 | 43 | 0 | 0 | 146 |
| 1 | 1766 | 58 | 43 | 0 | 0 | 139 |
| 1 | 1767 | 54 | 41 | 0.577 | 0.288 | 160 |
| 1 | 1768 | 54 | 41 | 0 | 0 | 162 |
| 1 | 1769 | 54 | 41 | 0 | 0 | 160 |
| 1 | 1770 | 55 | 47 | 0.785 | 0.392 | 166 |
| 1 | 1771 | 55 | 47 | 0 | 0 | 165 |
| 1 | 1772 | 55 | 47 | 0 | 0 | 144 |
| 1 | 1773 | 55 | 47 | 0 | 0 | 143 |
| 1 | 1774 | 55 | 47 | 0 | 0 | 139 |
| 1 | 1775 | 55 | 47 | 0 | 0 | 142 |
| 1 | 1776 | 55 | 47 | 0 | 0 | 154 |
| 1 | 1777 | 55 | 45 | 0.258 | 0.129 | 154 |
| 1 | 1778 | 54 | 46 | 0.182 | 0.091 | 142 |
| 1 | 1779 | 54 | 44 | 0.258 | 0.129 | 158 |
| 1 | 1780 | 53 | 42 | 0.288 | 0.144 | 160 |
| 1 | 1781 | 53 | 42 | 0 | 0 | 160 |
| 1 | 1782 | 53 | 42 | 0 | 0 | 160 |
| 1 | 1783 | 53 | 42 | 0 | 0 | 159 |
| 1 | 1784 | 53 | 42 | 0 | 0 | 159 |

| | | | | | | |
|---|---|---|---|---|---|---|
| 1 | 1785 | 53 | 42 | 0 | 0 | 160 |
| 1 | 1786 | 52 | 42 | 0.129 | 0.065 | 160 |
| 1 | 1787 | 52 | 42 | 0 | 0 | 159 |
| 1 | 1788 | 54 | 47 | 0.695 | 0.347 | 150 |
| 1 | 1789 | 54 | 47 | 0 | 0 | 149 |
| 1 | 1790 | 54 | 47 | 0 | 0 | 156 |
| 1 | 1791 | 54 | 47 | 0 | 0 | 145 |
| 1 | 1792 | 54 | 47 | 0 | 0 | 154 |
| 1 | 1793 | 54 | 47 | 0 | 0 | 154 |
| 1 | 1794 | 55 | 48 | 0.182 | 0.091 | 137 |
| 1 | 1795 | 55 | 48 | 0 | 0 | 138 |
| 1 | 1796 | 52 | 47 | 0.408 | 0.204 | 157 |
| 1 | 1797 | 52 | 47 | 0 | 0 | 156 |
| 1 | 1798 | 52 | 47 | 0 | 0 | 152 |
| 1 | 1799 | 52 | 47 | 0 | 0 | 148 |
| 1 | 1800 | 52 | 47 | 0 | 0 | 159 |
| 1 | 1801 | 52 | 47 | 0 | 0 | 157 |
| 1 | 1802 | 52 | 47 | 0 | 0 | 149 |
| 1 | 1803 | 52 | 47 | 0 | 0 | 150 |
| 1 | 1804 | 52 | 47 | 0 | 0 | 151 |
| 1 | 1805 | 52 | 47 | 0 | 0 | 151 |
| 1 | 1806 | 52 | 47 | 0 | 0 | 145 |
| 1 | 1807 | 52 | 47 | 0 | 0 | 142 |
| 1 | 1808 | 52 | 47 | 0 | 0 | 143 |
| 1 | 1809 | 52 | 47 | 0 | 0 | 142 |
| 1 | 1810 | 52 | 47 | 0 | 0 | 147 |
| 1 | 1811 | 49 | 47 | 0.387 | 0.194 | 158 |
| 1 | 1812 | 49 | 47 | 0 | 0 | 158 |
| 1 | 1813 | 49 | 47 | 0 | 0 | 163 |
| 1 | 1814 | 49 | 47 | 0 | 0 | 161 |
| 1 | 1815 | 48 | 47 | 0.129 | 0.065 | 161 |
| 1 | 1816 | 48 | 47 | 0 | 0 | 162 |
| 1 | 1817 | 48 | 47 | 0 | 0 | 163 |
| 1 | 1818 | 48 | 47 | 0 | 0 | 164 |
| 1 | 1819 | 48 | 48 | 0.129 | 0.065 | 160 |
| 1 | 1820 | 48 | 48 | 0 | 0 | 161 |
| 1 | 1821 | 48 | 48 | 0 | 0 | 163 |
| 1 | 1822 | 48 | 48 | 0 | 0 | 163 |
| 1 | 1823 | 47 | 48 | 0.129 | 0.065 | 161 |
| 1 | 1824 | 47 | 48 | 0 | 0 | 160 |
| 1 | 1825 | 47 | 48 | 0 | 0 | 160 |
| 1 | 1826 | 50 | 50 | 0.465 | 0.233 | 157 |
| 1 | 1827 | 50 | 50 | 0 | 0 | 148 |
| 1 | 1828 | 50 | 50 | 0 | 0 | 150 |
| 1 | 1829 | 50 | 50 | 0 | 0 | 159 |
| 1 | 1830 | 47 | 54 | 0.645 | 0.323 | 157 |
| 1 | 1831 | 47 | 54 | 0 | 0 | 158 |

| | | | | | | |
|---|---|---|---|---|---|---|
| 1 | 1832 | 47 | 54 | 0 | 0 | 159 |
| 1 | 1833 | 47 | 54 | 0 | 0 | 145 |
| 1 | 1834 | 47 | 54 | 0 | 0 | 150 |
| 1 | 1835 | 47 | 54 | 0 | 0 | 154 |
| 1 | 1836 | 47 | 54 | 0 | 0 | 146 |
| 1 | 1837 | 46 | 54 | 0.129 | 0.065 | 159 |
| 1 | 1838 | 46 | 54 | 0 | 0 | 157 |
| 1 | 1839 | 46 | 54 | 0 | 0 | 160 |
| 1 | 1840 | 46 | 54 | 0 | 0 | 156 |
| 1 | 1841 | 46 | 54 | 0 | 0 | 157 |
| 1 | 1842 | 46 | 54 | 0 | 0 | 153 |
| 1 | 1843 | 46 | 54 | 0 | 0 | 140 |
| 1 | 1844 | 46 | 54 | 0 | 0 | 132 |
| 1 | 1845 | 43 | 52 | 0.465 | 0.233 | 161 |
| 1 | 1846 | 43 | 52 | 0 | 0 | 163 |
| 1 | 1847 | 43 | 52 | 0 | 0 | 161 |
| 1 | 1848 | 43 | 52 | 0 | 0 | 160 |
| 1 | 1849 | 43 | 52 | 0 | 0 | 158 |
| 1 | 1850 | 42 | 51 | 0.182 | 0.091 | 162 |
| 1 | 1851 | 42 | 51 | 0 | 0 | 161 |
| 1 | 1852 | 42 | 51 | 0 | 0 | 161 |
| 1 | 1853 | 42 | 51 | 0 | 0 | 161 |
| 1 | 1854 | 42 | 54 | 0.387 | 0.194 | 158 |
| 1 | 1855 | 42 | 54 | 0 | 0 | 162 |
| 1 | 1856 | 42 | 54 | 0 | 0 | 162 |
| 1 | 1857 | 42 | 54 | 0 | 0 | 163 |
| 1 | 1858 | 42 | 54 | 0 | 0 | 163 |
| 1 | 1859 | 42 | 54 | 0 | 0 | 154 |
| 1 | 1860 | 42 | 54 | 0 | 0 | 157 |
| 1 | 1861 | 41 | 56 | 0.288 | 0.144 | 146 |
| 1 | 1862 | 41 | 56 | 0 | 0 | 146 |
| 1 | 1863 | 41 | 56 | 0 | 0 | 146 |
| 1 | 1864 | 41 | 56 | 0 | 0 | 144 |
| 1 | 1865 | 41 | 56 | 0 | 0 | 138 |
| 1 | 1866 | 41 | 55 | 0.129 | 0.065 | 139 |
| 1 | 1867 | 41 | 54 | 0.129 | 0.065 | 140 |
| 1 | 1868 | 41 | 53 | 0.129 | 0.065 | 152 |
| 1 | 1869 | 41 | 53 | 0 | 0 | 146 |
| 1 | 1870 | 41 | 53 | 0 | 0 | 149 |
| 1 | 1871 | 41 | 53 | 0 | 0 | 148 |
| 1 | 1872 | 41 | 53 | 0 | 0 | 144 |
| 1 | 1873 | 41 | 53 | 0 | 0 | 144 |
| 1 | 1874 | 41 | 53 | 0 | 0 | 145 |
| 1 | 1875 | 41 | 53 | 0 | 0 | 145 |
| 1 | 1876 | 41 | 53 | 0 | 0 | 148 |
| 1 | 1877 | 41 | 51 | 0.258 | 0.129 | 163 |
| 1 | 1878 | 41 | 51 | 0 | 0 | 166 |

| | | | | | | |
|---|---|---|---|---|---|---|
| 1 | 1879 | 41 | 51 | 0 | 0 | 162 |
| 1 | 1880 | 41 | 51 | 0 | 0 | 166 |
| 1 | 1881 | 41 | 51 | 0 | 0 | 162 |
| 1 | 1882 | 41 | 51 | 0 | 0 | 160 |
| 1 | 1883 | 41 | 51 | 0 | 0 | 163 |
| 1 | 1884 | 41 | 51 | 0 | 0 | 166 |
| 1 | 1885 | 41 | 51 | 0 | 0 | 165 |
| 1 | 1886 | 41 | 51 | 0 | 0 | 165 |
| 1 | 1887 | 41 | 51 | 0 | 0 | 165 |
| 1 | 1888 | 41 | 51 | 0 | 0 | 165 |
| 1 | 1889 | 41 | 51 | 0 | 0 | 161 |
| 1 | 1890 | 41 | 51 | 0 | 0 | 160 |
| 1 | 1891 | 38 | 54 | 0.547 | 0.274 | 162 |
| 1 | 1892 | 38 | 54 | 0 | 0 | 164 |
| 1 | 1893 | 38 | 54 | 0 | 0 | 163 |
| 1 | 1894 | 38 | 54 | 0 | 0 | 164 |
| 1 | 1895 | 38 | 54 | 0 | 0 | 166 |
| 1 | 1896 | 38 | 54 | 0 | 0 | 161 |
| 1 | 1897 | 38 | 54 | 0 | 0 | 162 |
| 1 | 1898 | 38 | 54 | 0 | 0 | 162 |
| 1 | 1899 | 38 | 54 | 0 | 0 | 161 |
| 1 | 1900 | 38 | 54 | 0 | 0 | 161 |
| 1 | 1901 | 38 | 54 | 0 | 0 | 165 |
| 1 | 1902 | 41 | 49 | 0.752 | 0.376 | 165 |
| 1 | 1903 | 41 | 49 | 0 | 0 | 165 |
| 1 | 1904 | 41 | 49 | 0 | 0 | 165 |
| 1 | 1905 | 41 | 49 | 0 | 0 | 164 |
| 1 | 1906 | 41 | 49 | 0 | 0 | 162 |
| 1 | 1907 | 41 | 49 | 0 | 0 | 165 |
| 1 | 1908 | 42 | 51 | 0.288 | 0.144 | 150 |
| 1 | 1909 | 42 | 51 | 0 | 0 | 152 |
| 1 | 1910 | 42 | 51 | 0 | 0 | 153 |
| 1 | 1911 | 42 | 51 | 0 | 0 | 150 |
| 1 | 1912 | 42 | 51 | 0 | 0 | 147 |
| 1 | 1913 | 42 | 51 | 0 | 0 | 152 |
| 1 | 1914 | 42 | 51 | 0 | 0 | 155 |
| 1 | 1915 | 42 | 51 | 0 | 0 | 156 |
| 1 | 1916 | 42 | 51 | 0 | 0 | 152 |
| 1 | 1917 | 43 | 50 | 0.182 | 0.091 | 153 |
| 1 | 1918 | 44 | 49 | 0.182 | 0.091 | 161 |
| 1 | 1919 | 44 | 49 | 0 | 0 | 157 |
| 1 | 1920 | 44 | 49 | 0 | 0 | 160 |
| 1 | 1921 | 44 | 49 | 0 | 0 | 156 |
| 1 | 1922 | 44 | 49 | 0 | 0 | 156 |
| 1 | 1923 | 44 | 48 | 0.129 | 0.065 | 157 |
| 1 | 1924 | 44 | 48 | 0 | 0 | 157 |
| 1 | 1925 | 44 | 48 | 0 | 0 | 155 |

| | | | | | | |
|---|---|---|---|---|---|---|
| 1 | 1926 | 45 | 46 | 0.288 | 0.144 | 163 |
| 1 | 1927 | 45 | 46 | 0 | 0 | 161 |
| 1 | 1928 | 45 | 46 | 0 | 0 | 149 |
| 1 | 1929 | 45 | 46 | 0 | 0 | 138 |
| 1 | 1930 | 47 | 45 | 0.288 | 0.144 | 135 |
| 1 | 1931 | 47 | 45 | 0 | 0 | 136 |
| 1 | 1932 | 47 | 45 | 0 | 0 | 137 |
| 1 | 1933 | 47 | 45 | 0 | 0 | 137 |
| 1 | 1934 | 47 | 45 | 0 | 0 | 136 |
| 1 | 1935 | 47 | 45 | 0 | 0 | 136 |
| 1 | 1936 | 47 | 45 | 0 | 0 | 152 |
| 1 | 1937 | 43 | 46 | 0.532 | 0.266 | 156 |
| 1 | 1938 | 43 | 46 | 0 | 0 | 157 |
| 1 | 1939 | 43 | 46 | 0 | 0 | 151 |
| 1 | 1940 | 43 | 46 | 0 | 0 | 145 |
| 1 | 1941 | 43 | 46 | 0 | 0 | 145 |
| 1 | 1942 | 43 | 46 | 0 | 0 | 141 |
| 1 | 1943 | 43 | 46 | 0 | 0 | 136 |
| 1 | 1944 | 43 | 46 | 0 | 0 | 137 |
| 1 | 1945 | 43 | 46 | 0 | 0 | 136 |
| 1 | 1946 | 43 | 46 | 0 | 0 | 132 |
| 1 | 1947 | 43 | 46 | 0 | 0 | 137 |
| 1 | 1948 | 43 | 46 | 0 | 0 | 138 |
| 1 | 1949 | 43 | 46 | 0 | 0 | 132 |
| 1 | 1950 | 43 | 44 | 0.258 | 0.129 | 155 |
| 1 | 1951 | 43 | 44 | 0 | 0 | 161 |
| 1 | 1952 | 43 | 44 | 0 | 0 | 158 |
| 1 | 1953 | 43 | 45 | 0.129 | 0.065 | 156 |
| 1 | 1954 | 43 | 45 | 0 | 0 | 135 |
| 1 | 1955 | 43 | 45 | 0 | 0 | 141 |
| 1 | 1956 | 43 | 45 | 0 | 0 | 137 |
| 1 | 1957 | 43 | 45 | 0 | 0 | 139 |
| 1 | 1958 | 43 | 45 | 0 | 0 | 132 |
| 1 | 1959 | 43 | 45 | 0 | 0 | 131 |
| 1 | 1960 | 43 | 45 | 0 | 0 | 135 |
| 1 | 1961 | 42 | 45 | 0.129 | 0.065 | 132 |
| 1 | 1962 | 42 | 45 | 0 | 0 | 139 |
| 1 | 1963 | 42 | 45 | 0 | 0 | 136 |
| 1 | 1964 | 41 | 45 | 0.129 | 0.065 | 136 |
| 1 | 1965 | 40 | 44 | 0.182 | 0.091 | 152 |
| 1 | 1966 | 40 | 44 | 0 | 0 | 157 |
| 1 | 1967 | 40 | 44 | 0 | 0 | 164 |
| 1 | 1968 | 40 | 44 | 0 | 0 | 161 |
| 1 | 1969 | 40 | 44 | 0 | 0 | 161 |
| 1 | 1970 | 40 | 44 | 0 | 0 | 164 |
| 1 | 1971 | 40 | 44 | 0 | 0 | 161 |
| 1 | 1972 | 40 | 44 | 0 | 0 | 158 |

| | | | | | | |
|---|---|---|---|---|---|---|
| 1 | 1973 | 40 | 44 | 0 | 0 | 158 |
| 1 | 1974 | 40 | 44 | 0 | 0 | 158 |
| 1 | 1975 | 40 | 44 | 0 | 0 | 157 |
| 1 | 1976 | 40 | 44 | 0 | 0 | 160 |
| 1 | 1977 | 40 | 44 | 0 | 0 | 154 |
| 1 | 1978 | 40 | 44 | 0 | 0 | 160 |
| 1 | 1979 | 40 | 44 | 0 | 0 | 159 |
| 1 | 1980 | 40 | 44 | 0 | 0 | 159 |
| 1 | 1981 | 40 | 44 | 0 | 0 | 160 |
| 1 | 1982 | 40 | 44 | 0 | 0 | 159 |
| 1 | 1983 | 39 | 43 | 0.182 | 0.091 | 159 |
| 1 | 1984 | 39 | 43 | 0 | 0 | 159 |
| 1 | 1985 | 39 | 43 | 0 | 0 | 160 |
| 1 | 1986 | 39 | 43 | 0 | 0 | 161 |
| 1 | 1987 | 39 | 43 | 0 | 0 | 162 |
| 1 | 1988 | 39 | 43 | 0 | 0 | 158 |
| 1 | 1989 | 39 | 43 | 0 | 0 | 159 |
| 1 | 1990 | 39 | 43 | 0 | 0 | 161 |
| 1 | 1991 | 39 | 43 | 0 | 0 | 160 |
| 1 | 1992 | 39 | 43 | 0 | 0 | 161 |
| 1 | 1993 | 39 | 43 | 0 | 0 | 161 |
| 1 | 1994 | 39 | 43 | 0 | 0 | 161 |
| 1 | 1995 | 39 | 43 | 0 | 0 | 161 |
| 1 | 1996 | 39 | 43 | 0 | 0 | 161 |
| 1 | 1997 | 39 | 43 | 0 | 0 | 161 |
| 1 | 1998 | 39 | 43 | 0 | 0 | 160 |
| 1 | 1999 | 39 | 43 | 0 | 0 | 163 |
| 1 | 2000 | 39 | 43 | 0 | 0 | 162 |
| 1 | 2001 | 39 | 43 | 0 | 0 | 166 |
| 1 | 2002 | 39 | 43 | 0 | 0 | 165 |
| 1 | 2003 | 39 | 43 | 0 | 0 | 165 |
| 1 | 2004 | 39 | 43 | 0 | 0 | 164 |
| 1 | 2005 | 39 | 43 | 0 | 0 | 163 |
| 1 | 2006 | 39 | 43 | 0 | 0 | 157 |
| 1 | 2007 | 39 | 43 | 0 | 0 | 161 |
| 1 | 2008 | 39 | 43 | 0 | 0 | 153 |
| 1 | 2009 | 39 | 41 | 0.258 | 0.129 | 162 |
| 1 | 2010 | 39 | 41 | 0 | 0 | 161 |
| 1 | 2011 | 39 | 38 | 0.387 | 0.194 | 159 |
| 1 | 2012 | 39 | 42 | 0.516 | 0.258 | 162 |
| 1 | 2013 | 40 | 42 | 0.129 | 0.065 | 149 |
| 1 | 2014 | 41 | 41 | 0.182 | 0.091 | 148 |
| 1 | 2015 | 41 | 41 | 0 | 0 | 137 |
| 1 | 2016 | 41 | 41 | 0 | 0 | 152 |
| 1 | 2017 | 42 | 40 | 0.182 | 0.091 | 138 |
| 1 | 2018 | 42 | 40 | 0 | 0 | 134 |
| 1 | 2019 | 42 | 40 | 0 | 0 | 138 |

| | | | | | | |
|---|---|---|---|---|---|---|
| 1 | 2020 | 42 | 40 | 0 | 0 | 138 |
| 1 | 2021 | 42 | 39 | 0.129 | 0.065 | 137 |
| 1 | 2022 | 42 | 39 | 0 | 0 | 136 |
| 1 | 2023 | 42 | 38 | 0.129 | 0.065 | 161 |
| 1 | 2024 | 43 | 37 | 0.182 | 0.091 | 158 |
| 1 | 2025 | 43 | 37 | 0 | 0 | 155 |
| 1 | 2026 | 43 | 37 | 0 | 0 | 161 |
| 1 | 2027 | 43 | 37 | 0 | 0 | 154 |
| 1 | 2028 | 39 | 34 | 0.645 | 0.323 | 158 |
| 1 | 2029 | 39 | 34 | 0 | 0 | 159 |
| 1 | 2030 | 39 | 34 | 0 | 0 | 159 |
| 1 | 2031 | 37 | 38 | 0.577 | 0.288 | 159 |
| 1 | 2032 | 37 | 38 | 0 | 0 | 161 |
| 1 | 2033 | 37 | 38 | 0 | 0 | 161 |
| 1 | 2034 | 37 | 38 | 0 | 0 | 161 |
| 1 | 2035 | 37 | 38 | 0 | 0 | 158 |
| 1 | 2036 | 36 | 36 | 0.288 | 0.144 | 159 |
| 1 | 2037 | 36 | 36 | 0 | 0 | 157 |
| 1 | 2038 | 36 | 36 | 0 | 0 | 157 |
| 1 | 2039 | 36 | 36 | 0 | 0 | 157 |
| 1 | 2040 | 36 | 36 | 0 | 0 | 157 |
| 1 | 2041 | 36 | 36 | 0 | 0 | 157 |
| 1 | 2042 | 36 | 36 | 0 | 0 | 158 |
| 1 | 2043 | 36 | 36 | 0 | 0 | 157 |
| 1 | 2044 | 36 | 36 | 0 | 0 | 157 |
| 1 | 2045 | 36 | 36 | 0 | 0 | 157 |
| 1 | 2046 | 36 | 36 | 0 | 0 | 157 |
| 1 | 2047 | 36 | 36 | 0 | 0 | 158 |
| 1 | 2048 | 36 | 36 | 0 | 0 | 160 |
| 1 | 2049 | 36 | 36 | 0 | 0 | 158 |
| 1 | 2050 | 36 | 36 | 0 | 0 | 158 |
| 1 | 2051 | 36 | 36 | 0 | 0 | 157 |
| 1 | 2052 | 36 | 36 | 0 | 0 | 156 |
| 1 | 2053 | 36 | 36 | 0 | 0 | 157 |
| 1 | 2054 | 36 | 35 | 0.129 | 0.065 | 161 |
| 1 | 2055 | 36 | 40 | 0.645 | 0.323 | 153 |
| 1 | 2056 | 36 | 39 | 0.129 | 0.065 | 160 |
| 1 | 2057 | 36 | 39 | 0 | 0 | 159 |
| 1 | 2058 | 36 | 39 | 0 | 0 | 153 |
| 1 | 2059 | 36 | 39 | 0 | 0 | 150 |
| 1 | 2060 | 36 | 38 | 0.129 | 0.065 | 159 |
| 1 | 2061 | 36 | 38 | 0 | 0 | 159 |
| 1 | 2062 | 36 | 38 | 0 | 0 | 157 |
| 1 | 2063 | 36 | 38 | 0 | 0 | 160 |
| 1 | 2064 | 36 | 37 | 0.129 | 0.065 | 167 |
| 1 | 2065 | 36 | 34 | 0.387 | 0.194 | 161 |
| 1 | 2066 | 36 | 34 | 0 | 0 | 165 |

| | | | | | | |
|---|---|---|---|---|---|---|
| 1 | 2067 | 38 | 39 | 0.695 | 0.347 | 140 |
| 1 | 2068 | 38 | 39 | 0 | 0 | 140 |
| 1 | 2069 | 38 | 39 | 0 | 0 | 138 |
| 1 | 2070 | 38 | 39 | 0 | 0 | 137 |
| 1 | 2071 | 38 | 39 | 0 | 0 | 139 |
| 1 | 2072 | 38 | 39 | 0 | 0 | 132 |
| 1 | 2073 | 38 | 39 | 0 | 0 | 139 |
| 1 | 2074 | 38 | 39 | 0 | 0 | 139 |
| 1 | 2075 | 38 | 39 | 0 | 0 | 136 |
| 1 | 2076 | 38 | 39 | 0 | 0 | 132 |
| 1 | 2077 | 38 | 38 | 0.129 | 0.065 | 135 |
| 1 | 2078 | 38 | 38 | 0 | 0 | 153 |
| 1 | 2079 | 38 | 38 | 0 | 0 | 152 |
| 1 | 2080 | 38 | 38 | 0 | 0 | 151 |
| 1 | 2081 | 38 | 38 | 0 | 0 | 160 |
| 1 | 2082 | 38 | 38 | 0 | 0 | 157 |
| 1 | 2083 | 38 | 38 | 0 | 0 | 160 |
| 1 | 2084 | 38 | 38 | 0 | 0 | 151 |
| 1 | 2085 | 38 | 38 | 0 | 0 | 157 |
| 1 | 2086 | 38 | 38 | 0 | 0 | 156 |
| 1 | 2087 | 38 | 38 | 0 | 0 | 156 |
| 1 | 2088 | 38 | 38 | 0 | 0 | 150 |
| 1 | 2089 | 38 | 38 | 0 | 0 | 151 |
| 1 | 2090 | 38 | 38 | 0 | 0 | 147 |
| 1 | 2091 | 38 | 38 | 0 | 0 | 156 |
| 1 | 2092 | 38 | 38 | 0 | 0 | 144 |
| 1 | 2093 | 38 | 38 | 0 | 0 | 150 |
| 1 | 2094 | 38 | 38 | 0 | 0 | 143 |
| 1 | 2095 | 38 | 38 | 0 | 0 | 154 |
| 1 | 2096 | 38 | 38 | 0 | 0 | 152 |
| 1 | 2097 | 38 | 38 | 0 | 0 | 162 |
| 1 | 2098 | 38 | 38 | 0 | 0 | 164 |
| 1 | 2099 | 38 | 38 | 0 | 0 | 161 |
| 1 | 2100 | 38 | 38 | 0 | 0 | 150 |
| 1 | 2101 | 38 | 38 | 0 | 0 | 158 |
| 1 | 2102 | 38 | 38 | 0 | 0 | 160 |
| 1 | 2103 | 38 | 38 | 0 | 0 | 155 |
| 1 | 2104 | 38 | 38 | 0 | 0 | 155 |
| 1 | 2105 | 38 | 38 | 0 | 0 | 156 |
| 1 | 2106 | 38 | 38 | 0 | 0 | 162 |
| 1 | 2107 | 38 | 38 | 0 | 0 | 163 |
| 1 | 2108 | 38 | 38 | 0 | 0 | 162 |
| 1 | 2109 | 38 | 38 | 0 | 0 | 161 |
| 1 | 2110 | 38 | 38 | 0 | 0 | 146 |
| 1 | 2111 | 38 | 38 | 0 | 0 | 142 |
| 1 | 2112 | 38 | 38 | 0 | 0 | 151 |
| 1 | 2113 | 38 | 38 | 0 | 0 | 141 |

| | | | | | | |
|---|---|---|---|---|---|---|
| 1 | 2114 | 33 | 39 | 0.658 | 0.329 | 163 |
| 1 | 2115 | 32 | 39 | 0.129 | 0.065 | 163 |
| 1 | 2116 | 32 | 39 | 0 | 0 | 166 |
| 1 | 2117 | 32 | 39 | 0 | 0 | 165 |
| 1 | 2118 | 32 | 39 | 0 | 0 | 161 |
| 1 | 2119 | 32 | 39 | 0 | 0 | 163 |
| 1 | 2120 | 32 | 39 | 0 | 0 | 162 |
| 1 | 2121 | 32 | 39 | 0 | 0 | 159 |
| 1 | 2122 | 32 | 39 | 0 | 0 | 162 |
| 1 | 2123 | 31 | 39 | 0.129 | 0.065 | 161 |
| 1 | 2124 | 31 | 39 | 0 | 0 | 161 |
| 1 | 2125 | 31 | 39 | 0 | 0 | 161 |
| 1 | 2126 | 30 | 37 | 0.288 | 0.144 | 159 |
| 1 | 2127 | 30 | 37 | 0 | 0 | 156 |
| 1 | 2128 | 30 | 37 | 0 | 0 | 159 |
| 1 | 2129 | 30 | 37 | 0 | 0 | 158 |
| 1 | 2130 | 30 | 37 | 0 | 0 | 159 |
| 1 | 2131 | 30 | 37 | 0 | 0 | 159 |
| 1 | 2132 | 30 | 37 | 0 | 0 | 159 |
| 1 | 2133 | 30 | 37 | 0 | 0 | 158 |
| 1 | 2134 | 30 | 37 | 0 | 0 | 159 |
| 1 | 2135 | 30 | 37 | 0 | 0 | 159 |
| 1 | 2136 | 30 | 37 | 0 | 0 | 159 |
| 1 | 2137 | 30 | 37 | 0 | 0 | 156 |
| 1 | 2138 | 30 | 37 | 0 | 0 | 159 |
| 1 | 2139 | 30 | 37 | 0 | 0 | 161 |
| 1 | 2140 | 30 | 37 | 0 | 0 | 161 |
| 1 | 2141 | 30 | 37 | 0 | 0 | 161 |
| 1 | 2142 | 30 | 37 | 0 | 0 | 162 |
| 1 | 2143 | 30 | 37 | 0 | 0 | 162 |
| 1 | 2144 | 30 | 37 | 0 | 0 | 160 |
| 1 | 2145 | 30 | 37 | 0 | 0 | 160 |
| 1 | 2146 | 30 | 37 | 0 | 0 | 160 |
| 1 | 2147 | 30 | 37 | 0 | 0 | 161 |
| 1 | 2148 | 30 | 37 | 0 | 0 | 159 |
| 1 | 2149 | 30 | 37 | 0 | 0 | 161 |
| 1 | 2150 | 30 | 37 | 0 | 0 | 163 |
| 1 | 2151 | 30 | 37 | 0 | 0 | 162 |
| 1 | 2152 | 30 | 37 | 0 | 0 | 160 |
| 1 | 2153 | 30 | 37 | 0 | 0 | 160 |
| 1 | 2154 | 30 | 37 | 0 | 0 | 158 |
| 1 | 2155 | 30 | 37 | 0 | 0 | 159 |
| 1 | 2156 | 30 | 37 | 0 | 0 | 159 |
| 1 | 2157 | 30 | 37 | 0 | 0 | 161 |
| 1 | 2158 | 30 | 37 | 0 | 0 | 158 |
| 1 | 2159 | 30 | 37 | 0 | 0 | 156 |
| 1 | 2160 | 30 | 36 | 0.129 | 0.065 | 156 |

| | | | | | | |
|---|---|---|---|---|---|---|
| 1 | 2161 | 30 | 36 | 0 | 0 | 156 |
| 1 | 2162 | 30 | 36 | 0 | 0 | 158 |
| 1 | 2163 | 30 | 36 | 0 | 0 | 158 |
| 1 | 2164 | 30 | 35 | 0.129 | 0.065 | 156 |
| 1 | 2165 | 30 | 35 | 0 | 0 | 156 |
| 1 | 2166 | 30 | 35 | 0 | 0 | 158 |
| 1 | 2167 | 29 | 34 | 0.182 | 0.091 | 157 |
| 1 | 2168 | 29 | 33 | 0.129 | 0.065 | 159 |
| 1 | 2169 | 29 | 33 | 0 | 0 | 159 |
| 1 | 2170 | 29 | 33 | 0 | 0 | 156 |
| 1 | 2171 | 29 | 33 | 0 | 0 | 159 |
| 1 | 2172 | 29 | 33 | 0 | 0 | 159 |
| 1 | 2173 | 29 | 33 | 0 | 0 | 161 |
| 1 | 2174 | 29 | 33 | 0 | 0 | 161 |
| 1 | 2175 | 29 | 33 | 0 | 0 | 159 |
| 1 | 2176 | 29 | 32 | 0.129 | 0.065 | 159 |
| 1 | 2177 | 29 | 32 | 0 | 0 | 160 |
| 1 | 2178 | 29 | 32 | 0 | 0 | 160 |
| 1 | 2179 | 29 | 32 | 0 | 0 | 159 |
| 1 | 2180 | 29 | 32 | 0 | 0 | 161 |
| 1 | 2181 | 29 | 32 | 0 | 0 | 164 |
| 1 | 2182 | 29 | 32 | 0 | 0 | 161 |
| 1 | 2183 | 29 | 32 | 0 | 0 | 160 |
| 1 | 2184 | 29 | 32 | 0 | 0 | 161 |
| 1 | 2185 | 29 | 32 | 0 | 0 | 160 |
| 1 | 2186 | 29 | 32 | 0 | 0 | 159 |
| 1 | 2187 | 29 | 32 | 0 | 0 | 160 |
| 1 | 2188 | 29 | 32 | 0 | 0 | 160 |
| 1 | 2189 | 29 | 32 | 0 | 0 | 159 |
| 1 | 2190 | 29 | 32 | 0 | 0 | 160 |
| 1 | 2191 | 29 | 32 | 0 | 0 | 159 |
| 1 | 2192 | 29 | 32 | 0 | 0 | 159 |
| 1 | 2193 | 29 | 32 | 0 | 0 | 159 |
| 1 | 2194 | 29 | 32 | 0 | 0 | 158 |
| 1 | 2195 | 29 | 32 | 0 | 0 | 158 |
| 1 | 2196 | 29 | 32 | 0 | 0 | 159 |
| 1 | 2197 | 29 | 32 | 0 | 0 | 159 |
| 1 | 2198 | 29 | 32 | 0 | 0 | 159 |
| 1 | 2199 | 29 | 32 | 0 | 0 | 159 |
| 1 | 2200 | 29 | 32 | 0 | 0 | 159 |
| 1 | 2201 | 29 | 32 | 0 | 0 | 159 |
| 1 | 2202 | 29 | 32 | 0 | 0 | 161 |
| 1 | 2203 | 29 | 32 | 0 | 0 | 160 |
| 1 | 2204 | 29 | 32 | 0 | 0 | 161 |
| 1 | 2205 | 29 | 32 | 0 | 0 | 160 |
| 1 | 2206 | 29 | 32 | 0 | 0 | 158 |
| 1 | 2207 | 29 | 32 | 0 | 0 | 158 |

| | | | | | | |
|---|---|---|---|---|---|---|
| 1 | 2208 | 29 | 32 | 0 | 0 | 161 |
| 1 | 2209 | 29 | 32 | 0 | 0 | 161 |
| 1 | 2210 | 29 | 32 | 0 | 0 | 160 |
| 1 | 2211 | 29 | 32 | 0 | 0 | 160 |
| 1 | 2212 | 29 | 32 | 0 | 0 | 158 |
| 1 | 2213 | 29 | 32 | 0 | 0 | 156 |
| 1 | 2214 | 29 | 32 | 0 | 0 | 158 |
| 1 | 2215 | 29 | 32 | 0 | 0 | 158 |
| 1 | 2216 | 29 | 32 | 0 | 0 | 159 |
| 1 | 2217 | 29 | 31 | 0.129 | 0.065 | 158 |
| 1 | 2218 | 29 | 31 | 0 | 0 | 159 |
| 1 | 2219 | 29 | 31 | 0 | 0 | 163 |
| 1 | 2220 | 29 | 31 | 0 | 0 | 162 |
| 1 | 2221 | 29 | 31 | 0 | 0 | 163 |
| 1 | 2222 | 29 | 31 | 0 | 0 | 162 |
| 1 | 2223 | 29 | 31 | 0 | 0 | 160 |
| 1 | 2224 | 29 | 31 | 0 | 0 | 158 |
| 1 | 2225 | 29 | 31 | 0 | 0 | 157 |
| 1 | 2226 | 29 | 31 | 0 | 0 | 160 |
| 1 | 2227 | 29 | 31 | 0 | 0 | 161 |
| 1 | 2228 | 29 | 31 | 0 | 0 | 158 |
| 1 | 2229 | 29 | 31 | 0 | 0 | 158 |
| 1 | 2230 | 29 | 31 | 0 | 0 | 158 |
| 1 | 2231 | 29 | 31 | 0 | 0 | 162 |
| 1 | 2232 | 29 | 31 | 0 | 0 | 162 |
| 1 | 2233 | 29 | 31 | 0 | 0 | 158 |
| 1 | 2234 | 29 | 31 | 0 | 0 | 160 |
| 1 | 2235 | 29 | 31 | 0 | 0 | 162 |
| 1 | 2236 | 29 | 31 | 0 | 0 | 163 |
| 1 | 2237 | 29 | 29 | 0.258 | 0.129 | 161 |
| 1 | 2238 | 29 | 29 | 0 | 0 | 162 |
| 1 | 2239 | 29 | 29 | 0 | 0 | 162 |
| 1 | 2240 | 29 | 29 | 0 | 0 | 162 |
| 1 | 2241 | 29 | 29 | 0 | 0 | 161 |
| 1 | 2242 | 29 | 29 | 0 | 0 | 162 |
| 1 | 2243 | 29 | 29 | 0 | 0 | 163 |
| 1 | 2244 | 29 | 27 | 0.258 | 0.129 | 161 |
| 1 | 2245 | 29 | 26 | 0.129 | 0.065 | 161 |
| 1 | 2246 | 29 | 26 | 0 | 0 | 161 |
| 1 | 2247 | 29 | 26 | 0 | 0 | 159 |
| 1 | 2248 | 27 | 26 | 0.258 | 0.129 | 162 |
| 1 | 2249 | 29 | 30 | 0.577 | 0.288 | 161 |
| 1 | 2250 | 29 | 30 | 0 | 0 | 159 |
| 1 | 2251 | 29 | 30 | 0 | 0 | 158 |
| 1 | 2252 | 29 | 30 | 0 | 0 | 159 |
| 1 | 2253 | 29 | 30 | 0 | 0 | 159 |
| 1 | 2254 | 29 | 30 | 0 | 0 | 163 |

| | | | | | | |
|---|---|---|---|---|---|---|
| 1 | 2255 | 29 | 30 | 0 | 0 | 161 |
| 1 | 2256 | 29 | 30 | 0 | 0 | 156 |
| 1 | 2257 | 29 | 30 | 0 | 0 | 158 |
| 1 | 2258 | 29 | 30 | 0 | 0 | 159 |
| 1 | 2259 | 29 | 30 | 0 | 0 | 158 |
| 1 | 2260 | 29 | 30 | 0 | 0 | 159 |
| 1 | 2261 | 29 | 30 | 0 | 0 | 158 |
| 1 | 2262 | 29 | 30 | 0 | 0 | 159 |
| 1 | 2263 | 29 | 30 | 0 | 0 | 161 |
| 1 | 2264 | 29 | 30 | 0 | 0 | 161 |
| 1 | 2265 | 29 | 30 | 0 | 0 | 163 |
| 1 | 2266 | 29 | 30 | 0 | 0 | 158 |
| 1 | 2267 | 29 | 30 | 0 | 0 | 157 |
| 1 | 2268 | 29 | 29 | 0.129 | 0.065 | 162 |
| 1 | 2269 | 29 | 29 | 0 | 0 | 163 |
| 1 | 2270 | 29 | 29 | 0 | 0 | 161 |
| 1 | 2271 | 29 | 29 | 0 | 0 | 161 |
| 1 | 2272 | 29 | 29 | 0 | 0 | 163 |
| 1 | 2273 | 29 | 29 | 0 | 0 | 164 |
| 1 | 2274 | 29 | 29 | 0 | 0 | 164 |
| 1 | 2275 | 29 | 29 | 0 | 0 | 161 |
| 1 | 2276 | 29 | 29 | 0 | 0 | 160 |
| 1 | 2277 | 29 | 29 | 0 | 0 | 159 |
| 1 | 2278 | 29 | 29 | 0 | 0 | 158 |
| 1 | 2279 | 29 | 29 | 0 | 0 | 157 |
| 1 | 2280 | 29 | 29 | 0 | 0 | 159 |
| 1 | 2281 | 29 | 29 | 0 | 0 | 157 |
| 1 | 2282 | 29 | 29 | 0 | 0 | 157 |
| 1 | 2283 | 29 | 29 | 0 | 0 | 139 |
| 1 | 2284 | 29 | 29 | 0 | 0 | 140 |
| 1 | 2285 | 29 | 29 | 0 | 0 | 149 |
| 1 | 2286 | 29 | 29 | 0 | 0 | 148 |
| 1 | 2287 | 29 | 29 | 0 | 0 | 150 |
| 1 | 2288 | 29 | 29 | 0 | 0 | 147 |
| 1 | 2289 | 29 | 29 | 0 | 0 | 148 |
| 1 | 2290 | 29 | 29 | 0 | 0 | 156 |
| 1 | 2291 | 29 | 29 | 0 | 0 | 148 |
| 1 | 2292 | 29 | 29 | 0 | 0 | 148 |
| 1 | 2293 | 29 | 29 | 0 | 0 | 144 |
| 1 | 2294 | 29 | 29 | 0 | 0 | 144 |
| 1 | 2295 | 29 | 29 | 0 | 0 | 143 |
| 1 | 2296 | 29 | 29 | 0 | 0 | 138 |
| 1 | 2297 | 29 | 29 | 0 | 0 | 138 |
| 1 | 2298 | 29 | 27 | 0.258 | 0.129 | 160 |
| 1 | 2299 | 29 | 27 | 0 | 0 | 155 |
| 1 | 2300 | 29 | 27 | 0 | 0 | 155 |
| 1 | 2301 | 29 | 27 | 0 | 0 | 164 |

| | | | | | | |
|---|---|---|---|---|---|---|
| 1 | 2302 | 28 | 25 | 0.288 | 0.144 | 161 |
| 1 | 2303 | 28 | 25 | 0 | 0 | 161 |
| 1 | 2304 | 28 | 25 | 0 | 0 | 161 |
| 1 | 2305 | 24 | 31 | 0.93 | 0.465 | 162 |
| 1 | 2306 | 24 | 31 | 0 | 0 | 162 |
| 1 | 2307 | 24 | 31 | 0 | 0 | 163 |
| 1 | 2308 | 24 | 31 | 0 | 0 | 163 |
| 1 | 2309 | 25 | 31 | 0.129 | 0.065 | 161 |
| 1 | 2310 | 25 | 31 | 0 | 0 | 161 |
| 1 | 2311 | 25 | 30 | 0.129 | 0.065 | 161 |
| 1 | 2312 | 25 | 29 | 0.129 | 0.065 | 162 |
| 1 | 2313 | 25 | 29 | 0 | 0 | 162 |
| 1 | 2314 | 25 | 28 | 0.129 | 0.065 | 162 |
| 1 | 2315 | 25 | 28 | 0 | 0 | 162 |
| 1 | 2316 | 25 | 28 | 0 | 0 | 162 |
| 1 | 2317 | 25 | 28 | 0 | 0 | 163 |
| 1 | 2318 | 25 | 28 | 0 | 0 | 163 |
| 1 | 2319 | 25 | 28 | 0 | 0 | 162 |
| 1 | 2320 | 25 | 28 | 0 | 0 | 160 |
| 1 | 2321 | 25 | 28 | 0 | 0 | 160 |
| 1 | 2322 | 25 | 28 | 0 | 0 | 161 |
| 1 | 2323 | 25 | 28 | 0 | 0 | 161 |
| 1 | 2324 | 25 | 28 | 0 | 0 | 160 |
| 1 | 2325 | 25 | 28 | 0 | 0 | 164 |
| 1 | 2326 | 25 | 28 | 0 | 0 | 161 |
| 1 | 2327 | 25 | 28 | 0 | 0 | 163 |
| 1 | 2328 | 25 | 28 | 0 | 0 | 160 |
| 1 | 2329 | 25 | 27 | 0.129 | 0.065 | 163 |
| 1 | 2330 | 25 | 27 | 0 | 0 | 164 |
| 1 | 2331 | 25 | 27 | 0 | 0 | 164 |
| 1 | 2332 | 25 | 27 | 0 | 0 | 165 |
| 1 | 2333 | 25 | 27 | 0 | 0 | 165 |
| 1 | 2334 | 25 | 27 | 0 | 0 | 165 |
| 1 | 2335 | 25 | 27 | 0 | 0 | 165 |
| 1 | 2336 | 25 | 27 | 0 | 0 | 163 |
| 1 | 2337 | 25 | 27 | 0 | 0 | 164 |
| 1 | 2338 | 25 | 27 | 0 | 0 | 164 |
| 1 | 2339 | 25 | 27 | 0 | 0 | 164 |
| 1 | 2340 | 25 | 27 | 0 | 0 | 164 |
| 1 | 2341 | 25 | 27 | 0 | 0 | 164 |
| 1 | 2342 | 25 | 27 | 0 | 0 | 164 |
| 1 | 2343 | 25 | 27 | 0 | 0 | 164 |
| 1 | 2344 | 25 | 27 | 0 | 0 | 164 |
| 1 | 2345 | 25 | 27 | 0 | 0 | 165 |
| 1 | 2346 | 25 | 27 | 0 | 0 | 162 |
| 1 | 2347 | 25 | 26 | 0.129 | 0.065 | 162 |
| 1 | 2348 | 25 | 26 | 0 | 0 | 163 |

| | | | | | | |
|---|---|---|---|---|---|---|
| 1 | 2349 | 25 | 26 | 0 | 0 | 163 |
| 1 | 2350 | 25 | 26 | 0 | 0 | 163 |
| 1 | 2351 | 25 | 26 | 0 | 0 | 164 |
| 1 | 2352 | 25 | 26 | 0 | 0 | 163 |
| 1 | 2353 | 25 | 26 | 0 | 0 | 159 |
| 1 | 2354 | 25 | 26 | 0 | 0 | 157 |
| 1 | 2355 | 25 | 26 | 0 | 0 | 156 |
| 1 | 2356 | 25 | 26 | 0 | 0 | 152 |
| 1 | 2357 | 25 | 26 | 0 | 0 | 159 |
| 1 | 2358 | 25 | 26 | 0 | 0 | 162 |
| 1 | 2359 | 25 | 26 | 0 | 0 | 159 |
| 1 | 2360 | 25 | 26 | 0 | 0 | 157 |
| 1 | 2361 | 25 | 26 | 0 | 0 | 157 |
| 1 | 2362 | 25 | 26 | 0 | 0 | 162 |
| 1 | 2363 | 25 | 26 | 0 | 0 | 154 |
| 1 | 2364 | 25 | 26 | 0 | 0 | 149 |
| 1 | 2365 | 25 | 26 | 0 | 0 | 156 |
| 1 | 2366 | 25 | 26 | 0 | 0 | 163 |
| 1 | 2367 | 25 | 26 | 0 | 0 | 154 |
| 1 | 2368 | 25 | 26 | 0 | 0 | 162 |
| 1 | 2369 | 25 | 26 | 0 | 0 | 152 |
| 1 | 2370 | 25 | 26 | 0 | 0 | 152 |
| 1 | 2371 | 25 | 26 | 0 | 0 | 154 |
| 1 | 2372 | 25 | 26 | 0 | 0 | 144 |
| 1 | 2373 | 25 | 26 | 0 | 0 | 146 |
| 1 | 2374 | 25 | 26 | 0 | 0 | 142 |
| 1 | 2375 | 25 | 26 | 0 | 0 | 141 |
| 1 | 2376 | 25 | 26 | 0 | 0 | 143 |
| 1 | 2377 | 25 | 26 | 0 | 0 | 140 |
| 1 | 2378 | 25 | 26 | 0 | 0 | 141 |
| 1 | 2379 | 25 | 26 | 0 | 0 | 139 |
| 1 | 2380 | 25 | 26 | 0 | 0 | 135 |
| 1 | 2381 | 25 | 26 | 0 | 0 | 133 |
| 1 | 2382 | 25 | 26 | 0 | 0 | 135 |
| 1 | 2383 | 25 | 26 | 0 | 0 | 150 |
| 1 | 2384 | 25 | 26 | 0 | 0 | 157 |
| 1 | 2385 | 25 | 26 | 0 | 0 | 150 |
| 1 | 2386 | 25 | 26 | 0 | 0 | 151 |
| 1 | 2387 | 25 | 26 | 0 | 0 | 147 |
| 1 | 2388 | 25 | 26 | 0 | 0 | 151 |
| 1 | 2389 | 23 | 23 | 0.465 | 0.233 | 161 |
| 1 | 2390 | 23 | 23 | 0 | 0 | 145 |
| 1 | 2391 | 23 | 23 | 0 | 0 | 139 |
| 1 | 2392 | 23 | 23 | 0 | 0 | 140 |
| 1 | 2393 | 23 | 23 | 0 | 0 | 139 |
| 1 | 2394 | 23 | 23 | 0 | 0 | 140 |
| 1 | 2395 | 23 | 23 | 0 | 0 | 142 |

| | | | | | | |
|---|---|---|---|---|---|---|
| 1 | 2396 | 23 | 23 | 0 | 0 | 154 |
| 1 | 2397 | 23 | 23 | 0 | 0 | 149 |
| 1 | 2398 | 23 | 23 | 0 | 0 | 162 |
| 1 | 2399 | 23 | 23 | 0 | 0 | 145 |
| 1 | 2400 | 23 | 23 | 0 | 0 | 142 |
| 1 | 2401 | 23 | 23 | 0 | 0 | 152 |
| 1 | 2402 | 23 | 23 | 0 | 0 | 138 |
| 1 | 2403 | 23 | 23 | 0 | 0 | 146 |
| 1 | 2404 | 23 | 23 | 0 | 0 | 153 |
| 1 | 2405 | 23 | 23 | 0 | 0 | 149 |
| 1 | 2406 | 22 | 22 | 0.182 | 0.091 | 165 |
| 1 | 2407 | 22 | 22 | 0 | 0 | 165 |
| 1 | 2408 | 22 | 22 | 0 | 0 | 165 |
| | | | **Total distance travelled (in µm)** | 127.531 | | |
| | | | **Average Velocity (µm/sec)** | | 0.02655 | |
