## Supplementary Data 1B for "Moving Yeasts: Resolving the Mystery"

| Supplementary Data 1B | | | | | | |
|---|---|---|---|---|---|---|
| Track n° | Slice n° | X | Y | Distance | Velocity | Pixel Value |
| 1 | 1 | 358 | 219 | -1 | -1 | 122 |
| 1 | 2 | 358 | 216 | 0.387 | 0.194 | 132 |
| 1 | 3 | 353 | 217 | 0.658 | 0.329 | 97 |
| 1 | 4 | 353 | 217 | 0 | 0 | 96 |
| 1 | 5 | 358 | 215 | 0.695 | 0.347 | 123 |
| 1 | 6 | 358 | 215 | 0 | 0 | 111 |
| 1 | 7 | 353 | 215 | 0.645 | 0.323 | 116 |
| 1 | 8 | 348 | 212 | 0.752 | 0.376 | 116 |
| 1 | 9 | 351 | 214 | 0.465 | 0.233 | 109 |
| 1 | 10 | 351 | 214 | 0 | 0 | 114 |
| 1 | 11 | 359 | 214 | 1.032 | 0.516 | 112 |
| 1 | 12 | 358 | 214 | 0.129 | 0.065 | 108 |
| 1 | 13 | 353 | 217 | 0.752 | 0.376 | 104 |
| 1 | 14 | 353 | 216 | 0.129 | 0.065 | 105 |
| 1 | 15 | 356 | 211 | 0.752 | 0.376 | 139 |
| 1 | 16 | 358 | 214 | 0.465 | 0.233 | 128 |
| 1 | 17 | 354 | 218 | 0.73 | 0.365 | 110 |
| 1 | 18 | 353 | 218 | 0.129 | 0.065 | 106 |
| 1 | 19 | 353 | 218 | 0 | 0 | 110 |
| 1 | 20 | 353 | 218 | 0 | 0 | 118 |
| 1 | 21 | 353 | 218 | 0 | 0 | 98 |
| 1 | 22 | 352 | 217 | 0.182 | 0.091 | 95 |
| 1 | 23 | 352 | 217 | 0 | 0 | 94 |
| 1 | 24 | 352 | 217 | 0 | 0 | 107 |
| 1 | 25 | 353 | 213 | 0.532 | 0.266 | 120 |
| 1 | 26 | 353 | 213 | 0 | 0 | 184 |
| 1 | 27 | 366 | 215 | 1.697 | 0.848 | 167 |
| 1 | 28 | 365 | 215 | 0.129 | 0.065 | 129 |
| 1 | 29 | 359 | 219 | 0.93 | 0.465 | 140 |
| 1 | 30 | 359 | 219 | 0 | 0 | 137 |
| 1 | 31 | 359 | 219 | 0 | 0 | 125 |
| 1 | 32 | 359 | 219 | 0 | 0 | 115 |
| 1 | 33 | 359 | 219 | 0 | 0 | 111 |
| 1 | 34 | 359 | 219 | 0 | 0 | 139 |
| 1 | 35 | 359 | 219 | 0 | 0 | 142 |
| 1 | 36 | 353 | 219 | 0.774 | 0.387 | 107 |
| 1 | 37 | 352 | 214 | 0.658 | 0.329 | 134 |
| 1 | 38 | 352 | 214 | 0 | 0 | 158 |
| 1 | 39 | 348 | 221 | 1.04 | 0.52 | 114 |
| 1 | 40 | 348 | 221 | 0 | 0 | 114 |
| 1 | 41 | 346 | 218 | 0.465 | 0.233 | 104 |
| 1 | 42 | 346 | 218 | 0 | 0 | 111 |
| 1 | 43 | 346 | 218 | 0 | 0 | 128 |
| 1 | 44 | 346 | 218 | 0 | 0 | 115 |
| 1 | 45 | 346 | 218 | 0 | 0 | 112 |

| | | | | | | |
|---|---|---|---|---|---|---|
| 1 | 46 | 346 | 218 | 0 | 0 | 107 |
| 1 | 47 | 346 | 218 | 0 | 0 | 103 |
| 1 | 48 | 346 | 218 | 0 | 0 | 108 |
| 1 | 49 | 347 | 213 | 0.658 | 0.329 | 150 |
| 1 | 50 | 352 | 213 | 0.645 | 0.323 | 153 |
| 1 | 51 | 352 | 213 | 0 | 0 | 129 |
| 1 | 52 | 352 | 213 | 0 | 0 | 151 |
| 1 | 53 | 352 | 213 | 0 | 0 | 111 |
| 1 | 54 | 346 | 211 | 0.816 | 0.408 | 148 |
| 1 | 55 | 348 | 217 | 0.816 | 0.408 | 103 |
| 1 | 56 | 348 | 217 | 0 | 0 | 101 |
| 1 | 57 | 348 | 216 | 0.129 | 0.065 | 101 |
| 1 | 58 | 348 | 216 | 0 | 0 | 102 |
| 1 | 59 | 348 | 216 | 0 | 0 | 115 |
| 1 | 60 | 348 | 216 | 0 | 0 | 103 |
| 1 | 61 | 347 | 211 | 0.658 | 0.329 | 136 |
| 1 | 62 | 347 | 211 | 0 | 0 | 119 |
| 1 | 63 | 347 | 211 | 0 | 0 | 127 |
| 1 | 64 | 347 | 211 | 0 | 0 | 134 |
| 1 | 65 | 347 | 211 | 0 | 0 | 145 |
| 1 | 66 | 347 | 211 | 0 | 0 | 138 |
| 1 | 67 | 355 | 209 | 1.064 | 0.532 | 151 |
| 1 | 68 | 356 | 209 | 0.129 | 0.065 | 146 |
| 1 | 69 | 356 | 209 | 0 | 0 | 139 |
| 1 | 70 | 356 | 209 | 0 | 0 | 147 |
| 1 | 71 | 356 | 209 | 0 | 0 | 103 |
| 1 | 72 | 357 | 216 | 0.912 | 0.456 | 120 |
| 1 | 73 | 357 | 214 | 0.258 | 0.129 | 116 |
| 1 | 74 | 359 | 208 | 0.816 | 0.408 | 138 |
| 1 | 75 | 359 | 208 | 0 | 0 | 138 |
| 1 | 76 | 359 | 211 | 0.387 | 0.194 | 145 |
| 1 | 77 | 359 | 211 | 0 | 0 | 121 |
| 1 | 78 | 359 | 211 | 0 | 0 | 126 |
| 1 | 79 | 360 | 206 | 0.658 | 0.329 | 129 |
| 1 | 80 | 360 | 207 | 0.129 | 0.065 | 135 |
| 1 | 81 | 360 | 207 | 0 | 0 | 129 |
| 1 | 82 | 358 | 207 | 0.258 | 0.129 | 129 |
| 1 | 83 | 358 | 207 | 0 | 0 | 121 |
| 1 | 84 | 358 | 206 | 0.129 | 0.065 | 137 |
| 1 | 85 | 358 | 206 | 0 | 0 | 132 |
| 1 | 86 | 358 | 206 | 0 | 0 | 128 |
| 1 | 87 | 358 | 208 | 0.258 | 0.129 | 134 |
| 1 | 88 | 358 | 208 | 0 | 0 | 119 |
| 1 | 89 | 358 | 207 | 0.129 | 0.065 | 121 |
| 1 | 90 | 358 | 207 | 0 | 0 | 128 |
| 1 | 91 | 358 | 207 | 0 | 0 | 132 |
| 1 | 92 | 358 | 206 | 0.129 | 0.065 | 129 |

| | | | | | | |
|---|---|---|---|---|---|---|
| 1 | 93 | 358 | 206 | 0 | 0 | 119 |
| 1 | 94 | 358 | 206 | 0 | 0 | 169 |
| 1 | 95 | 358 | 206 | 0 | 0 | 121 |
| 1 | 96 | 358 | 206 | 0 | 0 | 165 |
| 1 | 97 | 358 | 204 | 0.258 | 0.129 | 137 |
| 1 | 98 | 358 | 204 | 0 | 0 | 150 |
| 1 | 99 | 358 | 204 | 0 | 0 | 125 |
| 1 | 100 | 358 | 206 | 0.258 | 0.129 | 150 |
| 1 | 101 | 359 | 206 | 0.129 | 0.065 | 168 |
| 1 | 102 | 360 | 206 | 0.129 | 0.065 | 123 |
| 1 | 103 | 360 | 206 | 0 | 0 | 121 |
| 1 | 104 | 360 | 206 | 0 | 0 | 124 |
| 1 | 105 | 360 | 206 | 0 | 0 | 119 |
| 1 | 106 | 360 | 206 | 0 | 0 | 137 |
| 1 | 107 | 360 | 206 | 0 | 0 | 122 |
| 1 | 108 | 360 | 206 | 0 | 0 | 109 |
| 1 | 109 | 360 | 206 | 0 | 0 | 117 |
| 1 | 110 | 360 | 206 | 0 | 0 | 125 |
| 1 | 111 | 360 | 203 | 0.387 | 0.194 | 139 |
| 1 | 112 | 360 | 203 | 0 | 0 | 135 |
| 1 | 113 | 364 | 206 | 0.645 | 0.323 | 149 |
| 1 | 114 | 365 | 205 | 0.182 | 0.091 | 134 |
| 1 | 115 | 366 | 208 | 0.408 | 0.204 | 120 |
| 1 | 116 | 375 | 206 | 1.189 | 0.595 | 145 |
| 1 | 117 | 375 | 206 | 0 | 0 | 126 |
| 1 | 118 | 375 | 206 | 0 | 0 | 140 |
| 1 | 119 | 375 | 206 | 0 | 0 | 131 |
| 1 | 120 | 375 | 206 | 0 | 0 | 114 |
| 1 | 121 | 375 | 206 | 0 | 0 | 123 |
| 1 | 122 | 375 | 206 | 0 | 0 | 137 |
| 1 | 123 | 375 | 206 | 0 | 0 | 138 |
| 1 | 124 | 375 | 206 | 0 | 0 | 124 |
| 1 | 125 | 375 | 206 | 0 | 0 | 128 |
| 1 | 126 | 372 | 201 | 0.752 | 0.376 | 147 |
| 1 | 127 | 372 | 201 | 0 | 0 | 149 |
| 1 | 128 | 372 | 201 | 0 | 0 | 123 |
| 1 | 129 | 372 | 201 | 0 | 0 | 116 |
| 1 | 130 | 371 | 199 | 0.288 | 0.144 | 162 |
| 1 | 131 | 378 | 202 | 0.982 | 0.491 | 167 |
| 1 | 132 | 378 | 208 | 0.774 | 0.387 | 112 |
| 1 | 133 | 369 | 206 | 1.189 | 0.595 | 122 |
| 1 | 134 | 369 | 204 | 0.258 | 0.129 | 128 |
| 1 | 135 | 370 | 203 | 0.182 | 0.091 | 116 |
| 1 | 136 | 378 | 204 | 1.04 | 0.52 | 151 |
| 1 | 137 | 385 | 204 | 0.903 | 0.452 | 127 |
| 1 | 138 | 383 | 207 | 0.465 | 0.233 | 125 |
| 1 | 139 | 369 | 201 | 1.965 | 0.982 | 146 |

| | | | | | | |
|---|---|---|---|---|---|---|
| 1 | 140 | 369 | 201 | 0 | 0 | 112 |
| 1 | 141 | 369 | 201 | 0 | 0 | 119 |
| 1 | 142 | 369 | 201 | 0 | 0 | 144 |
| 1 | 143 | 377 | 203 | 1.064 | 0.532 | 114 |
| 1 | 144 | 377 | 203 | 0 | 0 | 108 |
| 1 | 145 | 377 | 203 | 0 | 0 | 112 |
| 1 | 146 | 377 | 203 | 0 | 0 | 110 |
| 1 | 147 | 377 | 203 | 0 | 0 | 123 |
| 1 | 148 | 377 | 203 | 0 | 0 | 121 |
| 1 | 149 | 377 | 203 | 0 | 0 | 114 |
| 1 | 150 | 377 | 203 | 0 | 0 | 114 |
| 1 | 151 | 377 | 203 | 0 | 0 | 120 |
| 1 | 152 | 377 | 203 | 0 | 0 | 117 |
| 1 | 153 | 377 | 203 | 0 | 0 | 115 |
| 1 | 154 | 377 | 203 | 0 | 0 | 134 |
| 1 | 155 | 377 | 203 | 0 | 0 | 143 |
| 1 | 156 | 377 | 203 | 0 | 0 | 139 |
| 1 | 157 | 377 | 203 | 0 | 0 | 121 |
| 1 | 158 | 377 | 195 | 1.032 | 0.516 | 123 |
| 1 | 159 | 377 | 195 | 0 | 0 | 123 |
| 1 | 160 | 377 | 194 | 0.129 | 0.065 | 124 |
| 1 | 161 | 377 | 194 | 0 | 0 | 123 |
| 1 | 162 | 378 | 190 | 0.532 | 0.266 | 122 |
| 1 | 163 | 378 | 190 | 0 | 0 | 115 |
| 1 | 164 | 378 | 190 | 0 | 0 | 120 |
| 1 | 165 | 378 | 190 | 0 | 0 | 124 |
| 1 | 166 | 378 | 190 | 0 | 0 | 134 |
| 1 | 167 | 378 | 190 | 0 | 0 | 132 |
| 1 | 168 | 378 | 190 | 0 | 0 | 142 |
| 1 | 169 | 378 | 190 | 0 | 0 | 135 |
| 1 | 170 | 378 | 190 | 0 | 0 | 127 |
| 1 | 171 | 377 | 182 | 1.04 | 0.52 | 131 |
| 1 | 172 | 375 | 178 | 0.577 | 0.288 | 138 |
| 1 | 173 | 375 | 178 | 0 | 0 | 148 |
| 1 | 174 | 375 | 174 | 0.516 | 0.258 | 140 |
| 1 | 175 | 375 | 174 | 0 | 0 | 167 |
| 1 | 176 | 373 | 175 | 0.288 | 0.144 | 162 |
| 1 | 177 | 373 | 175 | 0 | 0 | 143 |
| 1 | 178 | 370 | 172 | 0.547 | 0.274 | 136 |
| 1 | 179 | 370 | 168 | 0.516 | 0.258 | 154 |
| 1 | 180 | 370 | 173 | 0.645 | 0.323 | 125 |
| 1 | 181 | 370 | 173 | 0 | 0 | 129 |
| 1 | 182 | 376 | 176 | 0.865 | 0.433 | 122 |
| 1 | 183 | 376 | 176 | 0 | 0 | 116 |
| 1 | 184 | 376 | 176 | 0 | 0 | 121 |
| 1 | 185 | 376 | 176 | 0 | 0 | 119 |
| 1 | 186 | 376 | 176 | 0 | 0 | 118 |

| | | | | | | |
|---|---|---|---|---|---|---|
| 1 | 187 | 376 | 176 | 0 | 0 | 116 |
| 1 | 188 | 376 | 176 | 0 | 0 | 122 |
| 1 | 189 | 376 | 176 | 0 | 0 | 141 |
| 1 | 190 | 376 | 176 | 0 | 0 | 105 |
| 1 | 191 | 376 | 176 | 0 | 0 | 119 |
| 1 | 192 | 376 | 176 | 0 | 0 | 108 |
| 1 | 193 | 375 | 176 | 0.129 | 0.065 | 106 |
| 1 | 194 | 375 | 173 | 0.387 | 0.194 | 126 |
| 1 | 195 | 375 | 168 | 0.645 | 0.323 | 152 |
| 1 | 196 | 375 | 172 | 0.516 | 0.258 | 134 |
| 1 | 197 | 375 | 172 | 0 | 0 | 129 |
| 1 | 198 | 375 | 172 | 0 | 0 | 136 |
| 1 | 199 | 375 | 172 | 0 | 0 | 121 |
| 1 | 200 | 375 | 172 | 0 | 0 | 130 |
| 1 | 201 | 375 | 172 | 0 | 0 | 128 |
| 1 | 202 | 375 | 172 | 0 | 0 | 118 |
| 1 | 203 | 375 | 171 | 0.129 | 0.065 | 118 |
| 1 | 204 | 376 | 173 | 0.288 | 0.144 | 123 |
| 1 | 205 | 376 | 173 | 0 | 0 | 133 |
| 1 | 206 | 376 | 173 | 0 | 0 | 114 |
| 1 | 207 | 376 | 168 | 0.645 | 0.323 | 139 |
| 1 | 208 | 376 | 170 | 0.258 | 0.129 | 139 |
| 1 | 209 | 376 | 174 | 0.516 | 0.258 | 126 |
| 1 | 210 | 382 | 175 | 0.785 | 0.392 | 110 |
| 1 | 211 | 382 | 175 | 0 | 0 | 115 |
| 1 | 212 | 382 | 175 | 0 | 0 | 116 |
| 1 | 213 | 382 | 174 | 0.129 | 0.065 | 111 |
| 1 | 214 | 382 | 174 | 0 | 0 | 112 |
| 1 | 215 | 382 | 174 | 0 | 0 | 111 |
| 1 | 216 | 382 | 174 | 0 | 0 | 111 |
| 1 | 217 | 379 | 168 | 0.865 | 0.433 | 130 |
| 1 | 218 | 379 | 168 | 0 | 0 | 158 |
| 1 | 219 | 387 | 167 | 1.04 | 0.52 | 141 |
| 1 | 220 | 386 | 168 | 0.182 | 0.091 | 145 |
| 1 | 221 | 386 | 168 | 0 | 0 | 146 |
| 1 | 222 | 383 | 166 | 0.465 | 0.233 | 126 |
| 1 | 223 | 392 | 167 | 1.168 | 0.584 | 111 |
| 1 | 224 | 392 | 168 | 0.129 | 0.065 | 117 |
| 1 | 225 | 392 | 168 | 0 | 0 | 119 |
| 1 | 226 | 389 | 165 | 0.547 | 0.274 | 126 |
| 1 | 227 | 390 | 165 | 0.129 | 0.065 | 151 |
| 1 | 228 | 395 | 168 | 0.752 | 0.376 | 126 |
| 1 | 229 | 395 | 168 | 0 | 0 | 115 |
| 1 | 230 | 395 | 168 | 0 | 0 | 109 |
| 1 | 231 | 396 | 164 | 0.532 | 0.266 | 128 |
| 1 | 232 | 396 | 164 | 0 | 0 | 123 |
| 1 | 233 | 393 | 161 | 0.547 | 0.274 | 128 |

| | | | | | | |
|---|---|---|---|---|---|---|
| 1 | 234 | 392 | 168 | 0.912 | 0.456 | 133 |
| 1 | 235 | 392 | 168 | 0 | 0 | 111 |
| 1 | 236 | 392 | 162 | 0.774 | 0.387 | 133 |
| 1 | 237 | 392 | 162 | 0 | 0 | 113 |
| 1 | 238 | 392 | 162 | 0 | 0 | 110 |
| 1 | 239 | 392 | 162 | 0 | 0 | 129 |
| 1 | 240 | 392 | 162 | 0 | 0 | 130 |
| 1 | 241 | 392 | 162 | 0 | 0 | 91 |
| 1 | 242 | 392 | 161 | 0.129 | 0.065 | 91 |
| 1 | 243 | 391 | 159 | 0.288 | 0.144 | 164 |
| 1 | 244 | 391 | 159 | 0 | 0 | 153 |
| 1 | 245 | 391 | 159 | 0 | 0 | 157 |
| 1 | 246 | 391 | 159 | 0 | 0 | 123 |
| 1 | 247 | 391 | 159 | 0 | 0 | 152 |
| 1 | 248 | 391 | 159 | 0 | 0 | 146 |
| 1 | 249 | 391 | 159 | 0 | 0 | 145 |
| 1 | 250 | 391 | 159 | 0 | 0 | 133 |
| 1 | 251 | 391 | 159 | 0 | 0 | 145 |
| 1 | 252 | 391 | 159 | 0 | 0 | 114 |
| 1 | 253 | 391 | 159 | 0 | 0 | 133 |
| 1 | 254 | 391 | 159 | 0 | 0 | 128 |
| 1 | 255 | 391 | 159 | 0 | 0 | 125 |
| 1 | 256 | 391 | 159 | 0 | 0 | 144 |
| 1 | 257 | 392 | 159 | 0.129 | 0.065 | 140 |
| 1 | 258 | 392 | 159 | 0 | 0 | 140 |
| 1 | 259 | 392 | 159 | 0 | 0 | 127 |
| 1 | 260 | 392 | 159 | 0 | 0 | 106 |
| 1 | 261 | 392 | 153 | 0.774 | 0.387 | 130 |
| 1 | 262 | 392 | 153 | 0 | 0 | 136 |
| 1 | 263 | 392 | 153 | 0 | 0 | 146 |
| 1 | 264 | 392 | 158 | 0.645 | 0.323 | 105 |
| 1 | 265 | 392 | 158 | 0 | 0 | 79 |
| 1 | 266 | 392 | 155 | 0.387 | 0.194 | 104 |
| 1 | 267 | 392 | 155 | 0 | 0 | 102 |
| 1 | 268 | 392 | 155 | 0 | 0 | 118 |
| 1 | 269 | 392 | 155 | 0 | 0 | 78 |
| 1 | 270 | 391 | 152 | 0.408 | 0.204 | 134 |
| 1 | 271 | 391 | 152 | 0 | 0 | 117 |
| 1 | 272 | 391 | 152 | 0 | 0 | 127 |
| 1 | 273 | 391 | 152 | 0 | 0 | 126 |
| 1 | 274 | 391 | 152 | 0 | 0 | 139 |
| 1 | 275 | 391 | 152 | 0 | 0 | 139 |
| 1 | 276 | 391 | 152 | 0 | 0 | 125 |
| 1 | 277 | 391 | 152 | 0 | 0 | 104 |
| 1 | 278 | 391 | 152 | 0 | 0 | 78 |
| 1 | 279 | 391 | 152 | 0 | 0 | 98 |
| 1 | 280 | 392 | 154 | 0.288 | 0.144 | 91 |

| | | | | | | |
|---|---|---|---|---|---|---|
| 1 | 281 | 392 | 154 | 0 | 0 | 98 |
| 1 | 282 | 392 | 154 | 0 | 0 | 155 |
| 1 | 283 | 397 | 154 | 0.645 | 0.323 | 111 |
| 1 | 284 | 397 | 154 | 0 | 0 | 151 |
| 1 | 285 | 397 | 154 | 0 | 0 | 124 |
| 1 | 286 | 398 | 154 | 0.129 | 0.065 | 105 |
| 1 | 287 | 398 | 154 | 0 | 0 | 90 |
| 1 | 288 | 399 | 153 | 0.182 | 0.091 | 87 |
| 1 | 289 | 399 | 147 | 0.774 | 0.387 | 117 |
| 1 | 290 | 399 | 147 | 0 | 0 | 94 |
| 1 | 291 | 402 | 146 | 0.408 | 0.204 | 99 |
| 1 | 292 | 402 | 146 | 0 | 0 | 110 |
| 1 | 293 | 402 | 144 | 0.258 | 0.129 | 104 |
| 1 | 294 | 402 | 144 | 0 | 0 | 94 |
| 1 | 295 | 402 | 144 | 0 | 0 | 108 |
| 1 | 296 | 402 | 144 | 0 | 0 | 139 |
| 1 | 297 | 402 | 144 | 0 | 0 | 136 |
| 1 | 298 | 402 | 144 | 0 | 0 | 127 |
| 1 | 299 | 402 | 141 | 0.387 | 0.194 | 132 |
| 1 | 300 | 402 | 141 | 0 | 0 | 132 |
| 1 | 301 | 402 | 141 | 0 | 0 | 135 |
| 1 | 302 | 402 | 141 | 0 | 0 | 145 |
| 1 | 303 | 402 | 141 | 0 | 0 | 149 |
| 1 | 304 | 402 | 141 | 0 | 0 | 158 |
| 1 | 305 | 402 | 140 | 0.129 | 0.065 | 143 |
| 1 | 306 | 402 | 140 | 0 | 0 | 150 |
| 1 | 307 | 403 | 134 | 0.785 | 0.392 | 145 |
| 1 | 308 | 403 | 133 | 0.129 | 0.065 | 182 |
| 1 | 309 | 403 | 133 | 0 | 0 | 185 |
| 1 | 310 | 403 | 133 | 0 | 0 | 187 |
| 1 | 311 | 403 | 133 | 0 | 0 | 189 |
| 1 | 312 | 403 | 133 | 0 | 0 | 187 |
| 1 | 313 | 403 | 133 | 0 | 0 | 185 |
| 1 | 314 | 403 | 140 | 0.903 | 0.452 | 183 |
| 1 | 315 | 403 | 140 | 0 | 0 | 167 |
| 1 | 316 | 403 | 140 | 0 | 0 | 160 |
| 1 | 317 | 403 | 140 | 0 | 0 | 201 |
| 1 | 318 | 403 | 139 | 0.129 | 0.065 | 181 |
| 1 | 319 | 403 | 139 | 0 | 0 | 177 |
| 1 | 320 | 403 | 138 | 0.129 | 0.065 | 159 |
| 1 | 321 | 403 | 138 | 0 | 0 | 191 |
| 1 | 322 | 403 | 138 | 0 | 0 | 133 |
| 1 | 323 | 403 | 138 | 0 | 0 | 176 |
| 1 | 324 | 403 | 138 | 0 | 0 | 135 |
| 1 | 325 | 403 | 138 | 0 | 0 | 153 |
| 1 | 326 | 403 | 138 | 0 | 0 | 169 |
| 1 | 327 | 403 | 138 | 0 | 0 | 168 |

| | | | | | | |
|---|---|---|---|---|---|---|
| 1 | 328 | 403 | 138 | 0 | 0 | 128 |
| 1 | 329 | 403 | 138 | 0 | 0 | 175 |
| 1 | 330 | 403 | 138 | 0 | 0 | 147 |
| 1 | 331 | 403 | 138 | 0 | 0 | 169 |
| 1 | 332 | 403 | 138 | 0 | 0 | 176 |
| 1 | 333 | 403 | 138 | 0 | 0 | 154 |
| 1 | 334 | 403 | 138 | 0 | 0 | 184 |
| 1 | 335 | 403 | 138 | 0 | 0 | 179 |
| 1 | 336 | 403 | 138 | 0 | 0 | 178 |
| 1 | 337 | 403 | 138 | 0 | 0 | 164 |
| 1 | 338 | 403 | 138 | 0 | 0 | 132 |
| 1 | 339 | 403 | 138 | 0 | 0 | 119 |
| 1 | 340 | 403 | 138 | 0 | 0 | 136 |
| 1 | 341 | 403 | 138 | 0 | 0 | 136 |
| 1 | 342 | 403 | 138 | 0 | 0 | 130 |
| 1 | 343 | 403 | 138 | 0 | 0 | 135 |
| 1 | 344 | 403 | 138 | 0 | 0 | 144 |
| 1 | 345 | 403 | 138 | 0 | 0 | 126 |
| 1 | 346 | 403 | 138 | 0 | 0 | 143 |
| 1 | 347 | 403 | 136 | 0.258 | 0.129 | 144 |
| 1 | 348 | 403 | 136 | 0 | 0 | 129 |
| 1 | 349 | 403 | 136 | 0 | 0 | 134 |
| 1 | 350 | 403 | 136 | 0 | 0 | 131 |
| 1 | 351 | 403 | 136 | 0 | 0 | 117 |
| 1 | 352 | 403 | 136 | 0 | 0 | 122 |
| 1 | 353 | 403 | 132 | 0.516 | 0.258 | 153 |
| 1 | 354 | 403 | 132 | 0 | 0 | 167 |
| 1 | 355 | 403 | 132 | 0 | 0 | 121 |
| 1 | 356 | 403 | 132 | 0 | 0 | 125 |
| 1 | 357 | 403 | 132 | 0 | 0 | 121 |
| 1 | 358 | 403 | 132 | 0 | 0 | 119 |
| 1 | 359 | 403 | 132 | 0 | 0 | 107 |
| 1 | 360 | 403 | 129 | 0.387 | 0.194 | 132 |
| 1 | 361 | 403 | 129 | 0 | 0 | 114 |
| 1 | 362 | 403 | 129 | 0 | 0 | 171 |
| 1 | 363 | 403 | 129 | 0 | 0 | 122 |
| 1 | 364 | 403 | 129 | 0 | 0 | 122 |
| 1 | 365 | 403 | 129 | 0 | 0 | 150 |
| 1 | 366 | 403 | 129 | 0 | 0 | 154 |
| 1 | 367 | 403 | 129 | 0 | 0 | 148 |
| 1 | 368 | 403 | 129 | 0 | 0 | 175 |
| 1 | 369 | 403 | 129 | 0 | 0 | 135 |
| 1 | 370 | 403 | 129 | 0 | 0 | 178 |
| 1 | 371 | 403 | 129 | 0 | 0 | 177 |
| 1 | 372 | 400 | 127 | 0.465 | 0.233 | 149 |
| 1 | 373 | 400 | 127 | 0 | 0 | 166 |
| 1 | 374 | 400 | 127 | 0 | 0 | 168 |

| | | | | | | |
|---|---|---|---|---|---|---|
| 1 | 375 | 404 | 136 | 1.271 | 0.635 | 132 |
| 1 | 376 | 404 | 136 | 0 | 0 | 101 |
| 1 | 377 | 404 | 136 | 0 | 0 | 115 |
| 1 | 378 | 404 | 136 | 0 | 0 | 98 |
| 1 | 379 | 404 | 132 | 0.516 | 0.258 | 99 |
| 1 | 380 | 404 | 132 | 0 | 0 | 149 |
| 1 | 381 | 402 | 128 | 0.577 | 0.288 | 118 |
| 1 | 382 | 402 | 128 | 0 | 0 | 155 |
| 1 | 383 | 402 | 128 | 0 | 0 | 93 |
| 1 | 384 | 402 | 128 | 0 | 0 | 131 |
| 1 | 385 | 402 | 128 | 0 | 0 | 150 |
| 1 | 386 | 402 | 128 | 0 | 0 | 167 |
| 1 | 387 | 403 | 133 | 0.658 | 0.329 | 92 |
| 1 | 388 | 403 | 133 | 0 | 0 | 84 |
| 1 | 389 | 403 | 133 | 0 | 0 | 76 |
| 1 | 390 | 403 | 133 | 0 | 0 | 87 |
| 1 | 391 | 404 | 132 | 0.182 | 0.091 | 87 |
| 1 | 392 | 404 | 132 | 0 | 0 | 79 |
| 1 | 393 | 404 | 132 | 0 | 0 | 78 |
| 1 | 394 | 404 | 132 | 0 | 0 | 81 |
| 1 | 395 | 404 | 132 | 0 | 0 | 76 |
| 1 | 396 | 405 | 127 | 0.658 | 0.329 | 71 |
| 1 | 397 | 405 | 127 | 0 | 0 | 82 |
| 1 | 398 | 405 | 122 | 0.645 | 0.323 | 91 |
| 1 | 399 | 405 | 122 | 0 | 0 | 84 |
| 1 | 400 | 406 | 121 | 0.182 | 0.091 | 118 |
| 1 | 401 | 406 | 121 | 0 | 0 | 96 |
| 1 | 402 | 408 | 122 | 0.288 | 0.144 | 110 |
| 1 | 403 | 408 | 122 | 0 | 0 | 88 |
| 1 | 404 | 408 | 122 | 0 | 0 | 84 |
| 1 | 405 | 408 | 122 | 0 | 0 | 72 |
| 1 | 406 | 408 | 122 | 0 | 0 | 80 |
| 1 | 407 | 408 | 112 | 1.29 | 0.645 | 121 |
| 1 | 408 | 408 | 112 | 0 | 0 | 155 |
| 1 | 409 | 408 | 112 | 0 | 0 | 127 |
| 1 | 410 | 408 | 112 | 0 | 0 | 124 |
| 1 | 411 | 408 | 112 | 0 | 0 | 156 |
| 1 | 412 | 408 | 112 | 0 | 0 | 121 |
| 1 | 413 | 408 | 112 | 0 | 0 | 126 |
| 1 | 414 | 408 | 112 | 0 | 0 | 160 |
| 1 | 415 | 409 | 107 | 0.658 | 0.329 | 147 |
| 1 | 416 | 409 | 104 | 0.387 | 0.194 | 148 |
| 1 | 417 | 409 | 104 | 0 | 0 | 154 |
| 1 | 418 | 409 | 104 | 0 | 0 | 161 |
| 1 | 419 | 409 | 104 | 0 | 0 | 170 |
| 1 | 420 | 409 | 104 | 0 | 0 | 134 |
| 1 | 421 | 409 | 104 | 0 | 0 | 146 |

| | | | | | | |
|---|---|---|---|---|---|---|
| 1 | 422 | 409 | 104 | 0 | 0 | 174 |
| 1 | 423 | 409 | 104 | 0 | 0 | 146 |
| 1 | 424 | 409 | 104 | 0 | 0 | 133 |
| 1 | 425 | 409 | 104 | 0 | 0 | 127 |
| 1 | 426 | 409 | 104 | 0 | 0 | 142 |
| 1 | 427 | 409 | 104 | 0 | 0 | 129 |
| 1 | 428 | 409 | 104 | 0 | 0 | 139 |
| 1 | 429 | 401 | 103 | 1.04 | 0.52 | 134 |
| 1 | 430 | 401 | 103 | 0 | 0 | 138 |
| 1 | 431 | 401 | 103 | 0 | 0 | 147 |
| 1 | 432 | 401 | 103 | 0 | 0 | 130 |
| 1 | 433 | 401 | 106 | 0.387 | 0.194 | 141 |
| 1 | 434 | 401 | 106 | 0 | 0 | 163 |
| 1 | 435 | 401 | 106 | 0 | 0 | 179 |
| 1 | 436 | 401 | 106 | 0 | 0 | 177 |
| 1 | 437 | 400 | 103 | 0.408 | 0.204 | 154 |
| 1 | 438 | 396 | 103 | 0.516 | 0.258 | 144 |
| 1 | 439 | 397 | 103 | 0.129 | 0.065 | 154 |
| 1 | 440 | 397 | 103 | 0 | 0 | 178 |
| 1 | 441 | 397 | 98 | 0.645 | 0.323 | 138 |
| 1 | 442 | 397 | 98 | 0 | 0 | 130 |
| 1 | 443 | 397 | 98 | 0 | 0 | 142 |
| 1 | 444 | 397 | 98 | 0 | 0 | 136 |
| 1 | 445 | 397 | 98 | 0 | 0 | 144 |
| 1 | 446 | 397 | 98 | 0 | 0 | 136 |
| 1 | 447 | 397 | 98 | 0 | 0 | 157 |
| 1 | 448 | 397 | 98 | 0 | 0 | 141 |
| 1 | 449 | 397 | 98 | 0 | 0 | 151 |
| 1 | 450 | 397 | 98 | 0 | 0 | 157 |
| 1 | 451 | 397 | 98 | 0 | 0 | 157 |
| 1 | 452 | 397 | 98 | 0 | 0 | 162 |
| 1 | 453 | 397 | 98 | 0 | 0 | 149 |
| 1 | 454 | 397 | 98 | 0 | 0 | 142 |
| 1 | 455 | 397 | 98 | 0 | 0 | 154 |
| 1 | 456 | 397 | 98 | 0 | 0 | 134 |
| 1 | 457 | 397 | 98 | 0 | 0 | 143 |
| 1 | 458 | 397 | 98 | 0 | 0 | 148 |
| 1 | 459 | 397 | 98 | 0 | 0 | 143 |
| 1 | 460 | 397 | 98 | 0 | 0 | 141 |
| 1 | 461 | 397 | 98 | 0 | 0 | 139 |
| 1 | 462 | 397 | 98 | 0 | 0 | 153 |
| 1 | 463 | 397 | 98 | 0 | 0 | 148 |
| 1 | 464 | 397 | 98 | 0 | 0 | 146 |
| 1 | 465 | 400 | 100 | 0.465 | 0.233 | 147 |
| 1 | 466 | 400 | 100 | 0 | 0 | 140 |
| 1 | 467 | 400 | 100 | 0 | 0 | 138 |
| 1 | 468 | 400 | 100 | 0 | 0 | 142 |

| | | | | | | |
|---|---|---|---|---|---|---|
| 1 | 469 | 403 | 97 | 0.547 | 0.274 | 158 |
| 1 | 470 | 403 | 103 | 0.774 | 0.387 | 147 |
| 1 | 471 | 403 | 103 | 0 | 0 | 149 |
| 1 | 472 | 403 | 103 | 0 | 0 | 176 |
| 1 | 473 | 403 | 100 | 0.387 | 0.194 | 134 |
| 1 | 474 | 403 | 100 | 0 | 0 | 135 |
| 1 | 475 | 403 | 100 | 0 | 0 | 159 |
| 1 | 476 | 403 | 98 | 0.258 | 0.129 | 140 |
| 1 | 477 | 403 | 98 | 0 | 0 | 135 |
| 1 | 478 | 403 | 98 | 0 | 0 | 133 |
| 1 | 479 | 403 | 98 | 0 | 0 | 153 |
| 1 | 480 | 403 | 98 | 0 | 0 | 145 |
| 1 | 481 | 403 | 98 | 0 | 0 | 147 |
| 1 | 482 | 403 | 98 | 0 | 0 | 142 |
| 1 | 483 | 403 | 98 | 0 | 0 | 159 |
| 1 | 484 | 403 | 98 | 0 | 0 | 142 |
| 1 | 485 | 403 | 98 | 0 | 0 | 172 |
| 1 | 486 | 403 | 98 | 0 | 0 | 167 |
| 1 | 487 | 403 | 98 | 0 | 0 | 134 |
| 1 | 488 | 403 | 98 | 0 | 0 | 152 |
| 1 | 489 | 403 | 98 | 0 | 0 | 160 |
| 1 | 490 | 403 | 98 | 0 | 0 | 142 |
| 1 | 491 | 405 | 94 | 0.577 | 0.288 | 147 |
| 1 | 492 | 405 | 94 | 0 | 0 | 163 |
| 1 | 493 | 405 | 94 | 0 | 0 | 150 |
| 1 | 494 | 405 | 94 | 0 | 0 | 149 |
| 1 | 495 | 405 | 94 | 0 | 0 | 129 |
| 1 | 496 | 405 | 94 | 0 | 0 | 139 |
| 1 | 497 | 405 | 94 | 0 | 0 | 134 |
| 1 | 498 | 394 | 93 | 1.425 | 0.712 | 137 |
| 1 | 499 | 395 | 90 | 0.408 | 0.204 | 136 |
| 1 | 500 | 401 | 91 | 0.785 | 0.392 | 135 |
| 1 | 501 | 401 | 91 | 0 | 0 | 129 |
| 1 | 502 | 405 | 87 | 0.73 | 0.365 | 131 |
| 1 | 503 | 405 | 87 | 0 | 0 | 135 |
| 1 | 504 | 396 | 90 | 1.224 | 0.612 | 146 |
| 1 | 505 | 396 | 90 | 0 | 0 | 129 |
| 1 | 506 | 400 | 90 | 0.516 | 0.258 | 155 |
| 1 | 507 | 400 | 90 | 0 | 0 | 128 |
| 1 | 508 | 400 | 90 | 0 | 0 | 129 |
| 1 | 509 | 400 | 90 | 0 | 0 | 167 |
| 1 | 510 | 400 | 90 | 0 | 0 | 155 |
| 1 | 511 | 400 | 90 | 0 | 0 | 169 |
| 1 | 512 | 400 | 89 | 0.129 | 0.065 | 157 |
| 1 | 513 | 400 | 89 | 0 | 0 | 177 |
| 1 | 514 | 400 | 89 | 0 | 0 | 172 |
| 1 | 515 | 400 | 84 | 0.645 | 0.323 | 135 |

| | | | | | | |
|---|---|---|---|---|---|---|
| 1 | 516 | 400 | 84 | 0 | 0 | 143 |
| 1 | 517 | 400 | 84 | 0 | 0 | 145 |
| 1 | 518 | 404 | 84 | 0.516 | 0.258 | 141 |
| 1 | 519 | 404 | 84 | 0 | 0 | 138 |
| 1 | 520 | 404 | 84 | 0 | 0 | 130 |
| 1 | 521 | 404 | 84 | 0 | 0 | 154 |
| 1 | 522 | 404 | 84 | 0 | 0 | 151 |
| 1 | 523 | 404 | 84 | 0 | 0 | 154 |
| 1 | 524 | 404 | 84 | 0 | 0 | 166 |
| 1 | 525 | 404 | 84 | 0 | 0 | 161 |
| 1 | 526 | 404 | 84 | 0 | 0 | 161 |
| 1 | 527 | 404 | 84 | 0 | 0 | 150 |
| 1 | 528 | 404 | 84 | 0 | 0 | 134 |
| 1 | 529 | 404 | 84 | 0 | 0 | 142 |
| 1 | 530 | 404 | 84 | 0 | 0 | 131 |
| 1 | 531 | 404 | 84 | 0 | 0 | 142 |
| 1 | 532 | 404 | 84 | 0 | 0 | 139 |
| 1 | 533 | 404 | 84 | 0 | 0 | 147 |
| 1 | 534 | 404 | 84 | 0 | 0 | 145 |
| 1 | 535 | 404 | 84 | 0 | 0 | 149 |
| 1 | 536 | 404 | 84 | 0 | 0 | 139 |
| 1 | 537 | 404 | 84 | 0 | 0 | 144 |
| 1 | 538 | 404 | 84 | 0 | 0 | 145 |
| 1 | 539 | 404 | 84 | 0 | 0 | 166 |
| 1 | 540 | 404 | 84 | 0 | 0 | 164 |
| 1 | 541 | 404 | 84 | 0 | 0 | 196 |
| 1 | 542 | 404 | 84 | 0 | 0 | 142 |
| 1 | 543 | 404 | 84 | 0 | 0 | 148 |
| 1 | 544 | 404 | 84 | 0 | 0 | 140 |
| 1 | 545 | 404 | 84 | 0 | 0 | 161 |
| 1 | 546 | 404 | 84 | 0 | 0 | 149 |
| 1 | 547 | 404 | 84 | 0 | 0 | 161 |
| 1 | 548 | 405 | 81 | 0.408 | 0.204 | 159 |
| 1 | 549 | 405 | 81 | 0 | 0 | 150 |
| 1 | 550 | 405 | 81 | 0 | 0 | 170 |
| 1 | 551 | 405 | 81 | 0 | 0 | 156 |
| 1 | 552 | 405 | 81 | 0 | 0 | 133 |
| 1 | 553 | 404 | 77 | 0.532 | 0.266 | 136 |
| 1 | 554 | 404 | 77 | 0 | 0 | 175 |
| 1 | 555 | 404 | 77 | 0 | 0 | 132 |
| 1 | 556 | 404 | 77 | 0 | 0 | 162 |
| 1 | 557 | 404 | 77 | 0 | 0 | 184 |
| 1 | 558 | 404 | 77 | 0 | 0 | 164 |
| 1 | 559 | 404 | 77 | 0 | 0 | 162 |
| 1 | 560 | 404 | 77 | 0 | 0 | 190 |
| 1 | 561 | 406 | 81 | 0.577 | 0.288 | 137 |
| 1 | 562 | 406 | 81 | 0 | 0 | 176 |

| | | | | | | |
|---|---|---|---|---|---|---|
| 1 | 563 | 406 | 81 | 0 | 0 | 147 |
| 1 | 564 | 406 | 81 | 0 | 0 | 125 |
| 1 | 565 | 406 | 81 | 0 | 0 | 180 |
| 1 | 566 | 406 | 81 | 0 | 0 | 150 |
| 1 | 567 | 406 | 81 | 0 | 0 | 138 |
| 1 | 568 | 406 | 81 | 0 | 0 | 139 |
| 1 | 569 | 393 | 80 | 1.682 | 0.841 | 134 |
| 1 | 570 | 393 | 80 | 0 | 0 | 134 |
| 1 | 571 | 393 | 80 | 0 | 0 | 134 |
| 1 | 572 | 393 | 80 | 0 | 0 | 146 |
| 1 | 573 | 393 | 80 | 0 | 0 | 126 |
| 1 | 574 | 393 | 80 | 0 | 0 | 155 |
| 1 | 575 | 393 | 80 | 0 | 0 | 150 |
| 1 | 576 | 393 | 80 | 0 | 0 | 182 |
| 1 | 577 | 393 | 80 | 0 | 0 | 170 |
| 1 | 578 | 393 | 80 | 0 | 0 | 142 |
| 1 | 579 | 393 | 80 | 0 | 0 | 160 |
| 1 | 580 | 393 | 80 | 0 | 0 | 146 |
| 1 | 581 | 393 | 80 | 0 | 0 | 151 |
| 1 | 582 | 393 | 80 | 0 | 0 | 110 |
| 1 | 583 | 393 | 80 | 0 | 0 | 131 |
| 1 | 584 | 393 | 80 | 0 | 0 | 141 |
| 1 | 585 | 393 | 76 | 0.516 | 0.258 | 122 |
| 1 | 586 | 393 | 76 | 0 | 0 | 144 |
| 1 | 587 | 393 | 76 | 0 | 0 | 163 |
| 1 | 588 | 393 | 76 | 0 | 0 | 130 |
| 1 | 589 | 393 | 76 | 0 | 0 | 162 |
| 1 | 590 | 393 | 76 | 0 | 0 | 149 |
| 1 | 591 | 393 | 76 | 0 | 0 | 125 |
| 1 | 592 | 393 | 73 | 0.387 | 0.194 | 121 |
| 1 | 593 | 393 | 73 | 0 | 0 | 121 |
| 1 | 594 | 393 | 73 | 0 | 0 | 148 |
| 1 | 595 | 393 | 73 | 0 | 0 | 117 |
| 1 | 596 | 393 | 73 | 0 | 0 | 119 |
| 1 | 597 | 393 | 73 | 0 | 0 | 136 |
| 1 | 598 | 393 | 73 | 0 | 0 | 114 |
| 1 | 599 | 393 | 73 | 0 | 0 | 104 |
| 1 | 600 | 393 | 73 | 0 | 0 | 118 |
| 1 | 601 | 393 | 73 | 0 | 0 | 101 |
| 1 | 602 | 393 | 73 | 0 | 0 | 120 |
| 1 | 603 | 393 | 73 | 0 | 0 | 137 |
| 1 | 604 | 393 | 73 | 0 | 0 | 112 |
| 1 | 605 | 393 | 73 | 0 | 0 | 133 |
| 1 | 606 | 393 | 73 | 0 | 0 | 139 |
| 1 | 607 | 393 | 73 | 0 | 0 | 117 |
| 1 | 608 | 393 | 73 | 0 | 0 | 128 |
| 1 | 609 | 393 | 73 | 0 | 0 | 117 |

| | | | | | | |
|---|---|---|---|---|---|---|
| 1 | 610 | 393 | 73 | 0 | 0 | 129 |
| 1 | 611 | 393 | 73 | 0 | 0 | 114 |
| 1 | 612 | 393 | 73 | 0 | 0 | 105 |
| 1 | 613 | 393 | 71 | 0.258 | 0.129 | 118 |
| 1 | 614 | 393 | 71 | 0 | 0 | 123 |
| 1 | 615 | 393 | 71 | 0 | 0 | 108 |
| 1 | 616 | 393 | 71 | 0 | 0 | 115 |
| 1 | 617 | 393 | 71 | 0 | 0 | 109 |
| 1 | 618 | 393 | 71 | 0 | 0 | 113 |
| 1 | 619 | 393 | 71 | 0 | 0 | 157 |
| 1 | 620 | 393 | 71 | 0 | 0 | 117 |
| 1 | 621 | 393 | 71 | 0 | 0 | 160 |
| 1 | 622 | 393 | 71 | 0 | 0 | 188 |
| 1 | 623 | 393 | 70 | 0.129 | 0.065 | 177 |
| 1 | 624 | 393 | 70 | 0 | 0 | 108 |
| 1 | 625 | 393 | 70 | 0 | 0 | 114 |
| 1 | 626 | 393 | 70 | 0 | 0 | 95 |
| 1 | 627 | 393 | 70 | 0 | 0 | 112 |
| 1 | 628 | 393 | 70 | 0 | 0 | 111 |
| 1 | 629 | 393 | 70 | 0 | 0 | 103 |
| 1 | 630 | 393 | 70 | 0 | 0 | 115 |
| 1 | 631 | 393 | 70 | 0 | 0 | 101 |
| 1 | 632 | 393 | 70 | 0 | 0 | 105 |
| 1 | 633 | 393 | 70 | 0 | 0 | 137 |
| 1 | 634 | 393 | 70 | 0 | 0 | 105 |
| 1 | 635 | 393 | 70 | 0 | 0 | 102 |
| 1 | 636 | 393 | 70 | 0 | 0 | 105 |
| 1 | 637 | 393 | 65 | 0.645 | 0.323 | 171 |
| 1 | 638 | 393 | 65 | 0 | 0 | 111 |
| 1 | 639 | 393 | 65 | 0 | 0 | 172 |
| 1 | 640 | 393 | 65 | 0 | 0 | 184 |
| 1 | 641 | 393 | 65 | 0 | 0 | 155 |
| 1 | 642 | 393 | 65 | 0 | 0 | 144 |
| 1 | 643 | 393 | 65 | 0 | 0 | 165 |
| 1 | 644 | 393 | 65 | 0 | 0 | 130 |
| 1 | 645 | 393 | 65 | 0 | 0 | 117 |
| 1 | 646 | 393 | 65 | 0 | 0 | 105 |
| 1 | 647 | 393 | 65 | 0 | 0 | 100 |
| 1 | 648 | 393 | 65 | 0 | 0 | 127 |
| 1 | 649 | 393 | 65 | 0 | 0 | 127 |
| 1 | 650 | 393 | 65 | 0 | 0 | 104 |
| 1 | 651 | 393 | 65 | 0 | 0 | 90 |
| 1 | 652 | 393 | 65 | 0 | 0 | 101 |
| 1 | 653 | 393 | 65 | 0 | 0 | 101 |
| 1 | 654 | 393 | 65 | 0 | 0 | 134 |
| 1 | 655 | 393 | 65 | 0 | 0 | 106 |
| 1 | 656 | 393 | 65 | 0 | 0 | 95 |

| | | | | | | |
|---|---|---|---|---|---|---|
| 1 | 657 | 393 | 65 | 0 | 0 | 101 |
| 1 | 658 | 393 | 65 | 0 | 0 | 117 |
| 1 | 659 | 393 | 65 | 0 | 0 | 92 |
| 1 | 660 | 393 | 65 | 0 | 0 | 106 |
| 1 | 661 | 393 | 65 | 0 | 0 | 82 |
| 1 | 662 | 393 | 65 | 0 | 0 | 98 |
| 1 | 663 | 393 | 65 | 0 | 0 | 138 |
| 1 | 664 | 393 | 65 | 0 | 0 | 86 |
| 1 | 665 | 393 | 65 | 0 | 0 | 131 |
| 1 | 666 | 393 | 65 | 0 | 0 | 155 |
| 1 | 667 | 393 | 65 | 0 | 0 | 116 |
| 1 | 668 | 393 | 65 | 0 | 0 | 161 |
| 1 | 669 | 393 | 65 | 0 | 0 | 120 |
| 1 | 670 | 393 | 65 | 0 | 0 | 126 |
| 1 | 671 | 393 | 65 | 0 | 0 | 92 |
| 1 | 672 | 393 | 65 | 0 | 0 | 94 |
| 1 | 673 | 393 | 65 | 0 | 0 | 112 |
| 1 | 674 | 393 | 65 | 0 | 0 | 97 |
| 1 | 675 | 393 | 65 | 0 | 0 | 161 |
| 1 | 676 | 393 | 65 | 0 | 0 | 110 |
| 1 | 677 | 393 | 65 | 0 | 0 | 150 |
| 1 | 678 | 393 | 61 | 0.516 | 0.258 | 108 |
| 1 | 679 | 393 | 61 | 0 | 0 | 95 |
| 1 | 680 | 393 | 56 | 0.645 | 0.323 | 118 |
| 1 | 681 | 393 | 56 | 0 | 0 | 141 |
| 1 | 682 | 393 | 56 | 0 | 0 | 159 |
| 1 | 683 | 393 | 56 | 0 | 0 | 107 |
| 1 | 684 | 393 | 56 | 0 | 0 | 176 |
| 1 | 685 | 393 | 56 | 0 | 0 | 108 |
| 1 | 686 | 393 | 56 | 0 | 0 | 94 |
| 1 | 687 | 393 | 56 | 0 | 0 | 91 |
| 1 | 688 | 393 | 56 | 0 | 0 | 109 |
| 1 | 689 | 393 | 56 | 0 | 0 | 96 |
| 1 | 690 | 393 | 56 | 0 | 0 | 100 |
| 1 | 691 | 393 | 56 | 0 | 0 | 104 |
| 1 | 692 | 393 | 56 | 0 | 0 | 98 |
| 1 | 693 | 393 | 56 | 0 | 0 | 96 |
| 1 | 694 | 393 | 56 | 0 | 0 | 109 |
| 1 | 695 | 393 | 56 | 0 | 0 | 115 |
| 1 | 696 | 393 | 54 | 0.258 | 0.129 | 125 |
| 1 | 697 | 393 | 54 | 0 | 0 | 125 |
| 1 | 698 | 393 | 54 | 0 | 0 | 104 |
| 1 | 699 | 393 | 48 | 0.774 | 0.387 | 134 |
| 1 | 700 | 393 | 48 | 0 | 0 | 161 |
| 1 | 701 | 393 | 48 | 0 | 0 | 109 |
| 1 | 702 | 393 | 48 | 0 | 0 | 103 |
| 1 | 703 | 393 | 48 | 0 | 0 | 144 |

| | | | | | | |
|---|---|---|---|---|---|---|
| 1 | 704 | 393 | 48 | 0 | 0 | 95 |
| 1 | 705 | 393 | 48 | 0 | 0 | 98 |
| 1 | 706 | 393 | 48 | 0 | 0 | 90 |
| 1 | 707 | 393 | 47 | 0.129 | 0.065 | 117 |
| 1 | 708 | 393 | 47 | 0 | 0 | 98 |
| 1 | 709 | 393 | 43 | 0.516 | 0.258 | 166 |
| 1 | 710 | 393 | 43 | 0 | 0 | 174 |
| 1 | 711 | 393 | 43 | 0 | 0 | 111 |
| 1 | 712 | 393 | 41 | 0.258 | 0.129 | 124 |
| 1 | 713 | 393 | 41 | 0 | 0 | 120 |
| 1 | 714 | 393 | 41 | 0 | 0 | 127 |
| 1 | 715 | 393 | 41 | 0 | 0 | 102 |
| 1 | 716 | 393 | 41 | 0 | 0 | 133 |
| 1 | 717 | 393 | 41 | 0 | 0 | 122 |
| 1 | 718 | 393 | 41 | 0 | 0 | 144 |
| 1 | 719 | 393 | 41 | 0 | 0 | 171 |
| 1 | 720 | 393 | 41 | 0 | 0 | 151 |
| 1 | 721 | 393 | 41 | 0 | 0 | 148 |
| 1 | 722 | 393 | 41 | 0 | 0 | 165 |
| 1 | 723 | 393 | 41 | 0 | 0 | 170 |
| 1 | 724 | 387 | 45 | 0.93 | 0.465 | 114 |
| 1 | 725 | 387 | 45 | 0 | 0 | 105 |
| 1 | 726 | 387 | 45 | 0 | 0 | 112 |
| 1 | 727 | 387 | 45 | 0 | 0 | 118 |
| 1 | 728 | 387 | 45 | 0 | 0 | 161 |
| 1 | 729 | 387 | 45 | 0 | 0 | 114 |
| 1 | 730 | 387 | 45 | 0 | 0 | 171 |
| 1 | 731 | 387 | 45 | 0 | 0 | 105 |
| 1 | 732 | 387 | 45 | 0 | 0 | 111 |
| 1 | 733 | 387 | 45 | 0 | 0 | 99 |
| 1 | 734 | 387 | 45 | 0 | 0 | 83 |
| 1 | 735 | 387 | 45 | 0 | 0 | 106 |
| 1 | 736 | 388 | 44 | 0.182 | 0.091 | 122 |
| 1 | 737 | 388 | 44 | 0 | 0 | 142 |
| 1 | 738 | 388 | 44 | 0 | 0 | 117 |
| 1 | 739 | 388 | 44 | 0 | 0 | 108 |
| 1 | 740 | 388 | 44 | 0 | 0 | 111 |
| 1 | 741 | 388 | 44 | 0 | 0 | 103 |
| 1 | 742 | 388 | 44 | 0 | 0 | 109 |
| 1 | 743 | 388 | 42 | 0.258 | 0.129 | 154 |
| 1 | 744 | 388 | 42 | 0 | 0 | 167 |
| 1 | 745 | 388 | 42 | 0 | 0 | 116 |
| 1 | 746 | 388 | 42 | 0 | 0 | 163 |
| 1 | 747 | 388 | 42 | 0 | 0 | 162 |
| 1 | 748 | 388 | 42 | 0 | 0 | 176 |
| 1 | 749 | 388 | 42 | 0 | 0 | 181 |
| 1 | 750 | 388 | 42 | 0 | 0 | 181 |

| | | | | | | |
|---|---|---|---|---|---|---|
| 1 | 751 | 388 | 42 | 0 | 0 | 159 |
| 1 | 752 | 388 | 42 | 0 | 0 | 166 |
| 1 | 753 | 388 | 42 | 0 | 0 | 174 |
| 1 | 754 | 388 | 42 | 0 | 0 | 165 |
| 1 | 755 | 388 | 42 | 0 | 0 | 181 |
| 1 | 756 | 388 | 42 | 0 | 0 | 164 |
| 1 | 757 | 388 | 42 | 0 | 0 | 173 |
| 1 | 758 | 388 | 42 | 0 | 0 | 168 |
| 1 | 759 | 388 | 42 | 0 | 0 | 165 |
| 1 | 760 | 388 | 42 | 0 | 0 | 165 |
| 1 | 761 | 387 | 42 | 0.129 | 0.065 | 145 |
| 1 | 762 | 387 | 42 | 0 | 0 | 122 |
| 1 | 763 | 387 | 42 | 0 | 0 | 108 |
| 1 | 764 | 387 | 42 | 0 | 0 | 126 |
| 1 | 765 | 387 | 42 | 0 | 0 | 139 |
| 1 | 766 | 387 | 42 | 0 | 0 | 141 |
| 1 | 767 | 387 | 42 | 0 | 0 | 125 |
| 1 | 768 | 387 | 42 | 0 | 0 | 136 |
| 1 | 769 | 387 | 42 | 0 | 0 | 88 |
| 1 | 770 | 387 | 42 | 0 | 0 | 91 |
| 1 | 771 | 387 | 42 | 0 | 0 | 128 |
| 1 | 772 | 387 | 42 | 0 | 0 | 149 |
| 1 | 773 | 387 | 42 | 0 | 0 | 169 |
| 1 | 774 | 387 | 42 | 0 | 0 | 149 |
| 1 | 775 | 387 | 42 | 0 | 0 | 128 |
| 1 | 776 | 387 | 42 | 0 | 0 | 126 |
| 1 | 777 | 387 | 42 | 0 | 0 | 100 |
| 1 | 778 | 387 | 42 | 0 | 0 | 172 |
| 1 | 779 | 387 | 42 | 0 | 0 | 131 |
| 1 | 780 | 387 | 42 | 0 | 0 | 130 |
| 1 | 781 | 387 | 42 | 0 | 0 | 117 |
| 1 | 782 | 387 | 42 | 0 | 0 | 125 |
| 1 | 783 | 387 | 42 | 0 | 0 | 164 |
| 1 | 784 | 387 | 42 | 0 | 0 | 166 |
| 1 | 785 | 387 | 42 | 0 | 0 | 158 |
| 1 | 786 | 387 | 42 | 0 | 0 | 154 |
| 1 | 787 | 387 | 42 | 0 | 0 | 139 |
| 1 | 788 | 387 | 42 | 0 | 0 | 115 |
| 1 | 789 | 387 | 42 | 0 | 0 | 153 |
| 1 | 790 | 387 | 42 | 0 | 0 | 123 |
| 1 | 791 | 387 | 42 | 0 | 0 | 109 |
| 1 | 792 | 387 | 42 | 0 | 0 | 144 |
| 1 | 793 | 387 | 42 | 0 | 0 | 140 |
| 1 | 794 | 387 | 42 | 0 | 0 | 129 |
| 1 | 795 | 387 | 42 | 0 | 0 | 133 |
| 1 | 796 | 387 | 42 | 0 | 0 | 116 |
| 1 | 797 | 387 | 42 | 0 | 0 | 141 |

| | | | | | | |
|---|---|---|---|---|---|---|
| 1 | 798 | 387 | 42 | 0 | 0 | 118 |
| 1 | 799 | 387 | 42 | 0 | 0 | 133 |
| 1 | 800 | 387 | 42 | 0 | 0 | 163 |
| 1 | 801 | 387 | 42 | 0 | 0 | 144 |
| 1 | 802 | 387 | 42 | 0 | 0 | 114 |
| 1 | 803 | 387 | 42 | 0 | 0 | 165 |
| 1 | 804 | 387 | 42 | 0 | 0 | 146 |
| 1 | 805 | 382 | 43 | 0.658 | 0.329 | 101 |
| 1 | 806 | 382 | 43 | 0 | 0 | 109 |
| 1 | 807 | 382 | 43 | 0 | 0 | 112 |
| 1 | 808 | 382 | 43 | 0 | 0 | 127 |
| 1 | 809 | 382 | 43 | 0 | 0 | 129 |
| 1 | 810 | 382 | 43 | 0 | 0 | 141 |
| 1 | 811 | 382 | 39 | 0.516 | 0.258 | 84 |
| 1 | 812 | 382 | 39 | 0 | 0 | 106 |
| 1 | 813 | 382 | 39 | 0 | 0 | 139 |
| 1 | 814 | 382 | 39 | 0 | 0 | 126 |
| 1 | 815 | 382 | 39 | 0 | 0 | 107 |
| 1 | 816 | 382 | 39 | 0 | 0 | 119 |
| 1 | 817 | 382 | 39 | 0 | 0 | 107 |
| 1 | 818 | 382 | 39 | 0 | 0 | 93 |
| 1 | 819 | 382 | 39 | 0 | 0 | 120 |
| 1 | 820 | 382 | 39 | 0 | 0 | 97 |
| 1 | 821 | 382 | 39 | 0 | 0 | 104 |
| 1 | 822 | 382 | 39 | 0 | 0 | 96 |
| 1 | 823 | 382 | 39 | 0 | 0 | 82 |
| 1 | 824 | 382 | 39 | 0 | 0 | 80 |
| 1 | 825 | 382 | 39 | 0 | 0 | 75 |
| 1 | 826 | 382 | 27 | 1.548 | 0.774 | 101 |
| 1 | 827 | 382 | 27 | 0 | 0 | 112 |
| 1 | 828 | 382 | 27 | 0 | 0 | 90 |
| 1 | 829 | 382 | 27 | 0 | 0 | 122 |
| 1 | 830 | 382 | 27 | 0 | 0 | 93 |
| 1 | 831 | 382 | 27 | 0 | 0 | 127 |
| 1 | 832 | 382 | 27 | 0 | 0 | 117 |
| 1 | 833 | 382 | 27 | 0 | 0 | 121 |
| 1 | 834 | 382 | 27 | 0 | 0 | 142 |
| 1 | 835 | 382 | 27 | 0 | 0 | 109 |
| 1 | 836 | 382 | 27 | 0 | 0 | 118 |
| 1 | 837 | 382 | 27 | 0 | 0 | 126 |
| 1 | 838 | 382 | 27 | 0 | 0 | 124 |
| 1 | 839 | 382 | 27 | 0 | 0 | 136 |
| 1 | 840 | 382 | 27 | 0 | 0 | 124 |
| 1 | 841 | 382 | 27 | 0 | 0 | 142 |
| 1 | 842 | 382 | 27 | 0 | 0 | 153 |
| 1 | 843 | 382 | 27 | 0 | 0 | 161 |
| 1 | 844 | 382 | 27 | 0 | 0 | 166 |

| | | | | | | |
|---|---|---|---|---|---|---|
| 1 | 845 | 382 | 27 | 0 | 0 | 171 |
| 1 | 846 | 382 | 27 | 0 | 0 | 171 |
| 1 | 847 | 382 | 27 | 0 | 0 | 179 |
| 1 | 848 | 382 | 27 | 0 | 0 | 180 |
| 1 | 849 | 382 | 27 | 0 | 0 | 182 |
| 1 | 850 | 382 | 27 | 0 | 0 | 187 |
| 1 | 851 | 387 | 31 | 0.826 | 0.413 | 189 |
| 1 | 852 | 387 | 32 | 0.129 | 0.065 | 147 |
| 1 | 853 | 387 | 32 | 0 | 0 | 140 |
| 1 | 854 | 388 | 33 | 0.182 | 0.091 | 182 |
| 1 | 855 | 388 | 33 | 0 | 0 | 140 |
| 1 | 856 | 388 | 33 | 0 | 0 | 132 |
| 1 | 857 | 388 | 33 | 0 | 0 | 132 |
| | | | **Total distance travelled (in µm)** | 98.78 | | |
| | | | **Average Velocity (µm/sec)** | | 0.057734 | |
