## Supplementary Data 2 for "Moving Yeasts: Resolving the Mystery"

**Table. 1** Tracking of Particle inside Yeast cell

| Track n° | Slice n° | X | Y | Distance (µm) | Velocity (µm per sec) | Pixel Value |
| --- | --- | --- | --- | --- | --- | --- |
| 1 | 1 | 82 | 123 | -1 | -1 | -16755968 |
| 1 | 2 | 113 | 118 | 4.051 | 2.025 | -16748800 |
| 1 | 3 | 117 | 111 | 1.04 | 0.52 | -16758528 |
| 1 | 4 | 111 | 82 | 3.82 | 1.91 | -16758016 |
| 1 | 5 | 98 | 79 | 1.721 | 0.861 | -16750080 |
| 1 | 6 | 122 | 60 | 3.949 | 1.974 | -16768768 |
| 1 | 7 | 101 | 68 | 2.899 | 1.449 | -16754688 |
| 1 | 8 | 83 | 90 | 3.667 | 1.833 | -16752128 |
| 1 | 9 | 117 | 101 | 4.61 | 2.305 | -16751872 |
| 1 | 10 | 136 | 61 | 5.713 | 2.856 | -16751360 |
| 1 | 11 | 183 | 43 | 6.492 | 3.246 | -16750848 |
| 1 | 12 | 181 | 56 | 1.697 | 0.848 | -16746496 |
| 1 | 13 | 160 | 55 | 2.712 | 1.356 | -16741120 |
| 1 | 14 | 162 | 57 | 0.365 | 0.182 | -16752384 |
|  |  |  | **Total distance travelled** | **41.736** |  |  |
|  |  |  | **Average Velocity** |  | **1.454643** |  |

**Table 2.** Tracking of bacteria A outside yeast cell as shown in figure 5 and Supplementary video 6

| Track n° | Slice n° | X | Y | Distance (µm) | Velocity (µm per sec) | Pixel Value |
| --- | --- | --- | --- | --- | --- | --- |
| 1 | 1 | 389 | 292 | -1 | -1 | -16750592 |
| 1 | 2 | 390 | 293 | 0.182 | 0.091 | -16751872 |
| 1 | 3 | 393 | 302 | 1.224 | 0.612 | -16774144 |
| 1 | 4 | 388 | 301 | 0.658 | 0.329 | -16772352 |
| 1 | 5 | 385 | 295 | 0.865 | 0.433 | -16749568 |
| 1 | 6 | 386 | 298 | 0.408 | 0.204 | -16749312 |
| 1 | 7 | 386 | 303 | 0.645 | 0.323 | -16770816 |
| 1 | 8 | 383 | 304 | 0.408 | 0.204 | -16759552 |
| 1 | 9 | 386 | 307 | 0.547 | 0.274 | -16774144 |
| 1 | 10 | 383 | 307 | 0.387 | 0.194 | -16772864 |
| 1 | 11 | 380 | 305 | 0.465 | 0.233 | -16754176 |
| 1 | 12 | 380 | 306 | 0.129 | 0.065 | -16759040 |
| 1 | 13 | 378 | 311 | 0.695 | 0.347 | -16773888 |
| 1 | 14 | 380 | 312 | 0.288 | 0.144 | -16773888 |
|  |  |  | **Total distance travelled** | **5.901** |  |  |
|  |  |  | **Average Velocity** |  | **0.175214** |  |

**Table 3.** Tracking of bacteria B outside yeast cell as shown in figure 5 and Supplementary video 6

| Track n° | Slice n° | X | Y | Distance (µm) | Velocity (µm per sec) | Pixel Value |
| --- | --- | --- | --- | --- | --- | --- |
| 1 | 1 | 379 | 280 | -1 | -1 | -16754688 |
| 1 | 2 | 380 | 283 | 0.408 | 0.204 | -16767744 |
| 1 | 3 | 377 | 284 | 0.408 | 0.204 | -16765440 |
| 1 | 4 | 376 | 285 | 0.182 | 0.091 | -16768512 |
| 1 | 5 | 378 | 286 | 0.288 | 0.144 | -16770304 |
| 1 | 6 | 376 | 284 | 0.365 | 0.182 | -16762368 |
| 1 | 7 | 374 | 286 | 0.365 | 0.182 | -16767232 |
| 1 | 8 | 372 | 288 | 0.365 | 0.182 | -16770304 |
| 1 | 9 | 372 | 286 | 0.258 | 0.129 | -16767232 |
| 1 | 10 | 372 | 289 | 0.387 | 0.194 | -16771328 |
| 1 | 11 | 372 | 288 | 0.129 | 0.065 | -16764672 |
| 1 | 12 | 372 | 290 | 0.258 | 0.129 | -16774656 |
| 1 | 13 | 371 | 292 | 0.288 | 0.144 | -16764160 |
| 1 | 14 | 373 | 292 | 0.258 | 0.129 | -16766464 |
|  |  |  | **Total distance travelled** | **2.959** |  |  |
|  |  |  | **Average Velocity** |  | **0.069929** |  |

**Table 4.** Tracking of bacteria C outside yeast cell as shown in figure 5 and Supplementary video 6

| Track n° | Slice n° | X | Y | Distance (µm) | Velocity (µm per sec) | Pixel Value |
| --- | --- | --- | --- | --- | --- | --- |
| 1 | 1 | 397 | 255 | -1 | -1 | -16764160 |
| 1 | 2 | 399 | 261 | 0.816 | 0.408 | -16773888 |
| 1 | 3 | 393 | 259 | 0.816 | 0.408 | -16770048 |
| 1 | 4 | 395 | 259 | 0.258 | 0.129 | -16772096 |
| 1 | 5 | 393 | 260 | 0.288 | 0.144 | -16771072 |
| 1 | 6 | 399 | 262 | 0.816 | 0.408 | -16774144 |
| 1 | 7 | 392 | 265 | 0.982 | 0.491 | -16773120 |
| 1 | 8 | 392 | 266 | 0.129 | 0.065 | -16773120 |
| 1 | 9 | 397 | 268 | 0.695 | 0.347 | -16773376 |
| 1 | 10 | 393 | 265 | 0.645 | 0.323 | -16770304 |
| 1 | 11 | 388 | 263 | 0.695 | 0.347 | -16768256 |
| 1 | 12 | 391 | 268 | 0.752 | 0.376 | -16768000 |
| 1 | 13 | 391 | 267 | 0.129 | 0.065 | -16765184 |
| 1 | 14 | 388 | 270 | 0.547 | 0.274 | -16771840 |
|  |  |  | **Total distance travelled** | **6.568** |  |  |
|  |  |  | **Average Velocity** |  | **0.198929** |  |

**Table 5.** Comparative of Displacement and velocities for particle in yeast cell and Bacteria outside Yeast cell

| **Spot** | **Total Distance travelled (µm)** | **Velocity (µm per sec)** |
| --- | --- | --- |
| Spot inside Yeast | 41.73 | 1.45 |
| A | 5.901 | 0.175 |
| B | 2.959 | 0.0699 |
| C | 6.568 | 0.198 |
