## Supplementary Data 3 for "Moving Yeasts: Resolving the Mystery"

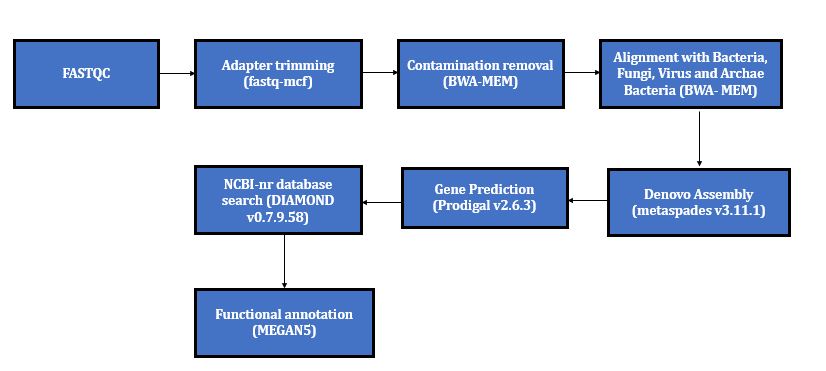

**Figure 1.** Bioinformatics workflow for metagenome analysis

**ORF Prediction**

**Table 1.** ORF length distribution

| **Total genes** | **genes>200bp** | **500bp>genes<1000bp** | **1000bp>genes** |
| --- | --- | --- | --- |
| 2637 | 13051 | 5192 | 4394 |

**Table 2.** Distribution of domain

| **Kingdom** | **Abundance percentage** |
| --- | --- |
| root;cellular organisms;Eukaryota; | 99.889 |
| root;cellular organisms;Bacteria; | 0.111 |

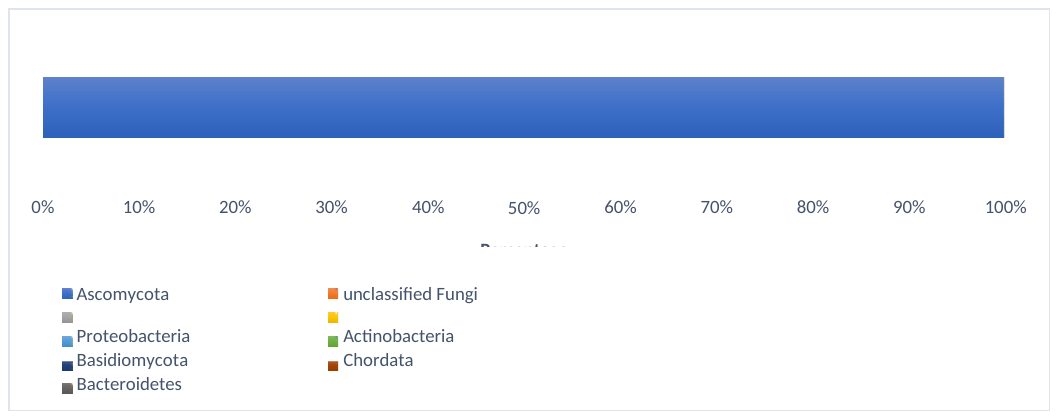

**Figure 2.** Distribution of Phylum

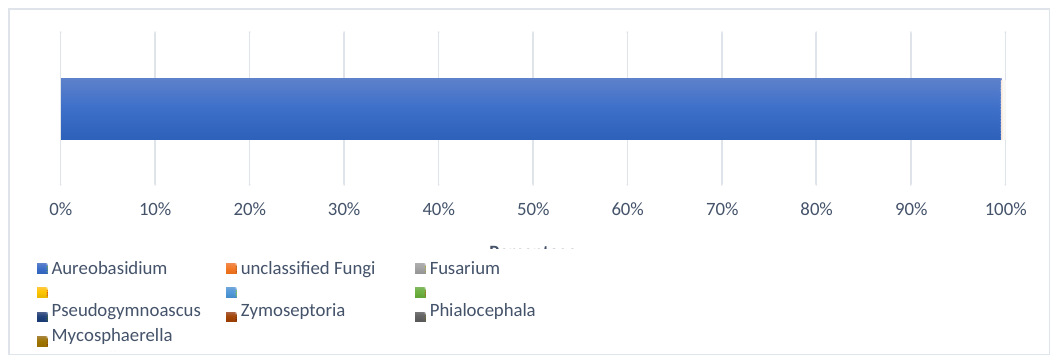

**Figure 3.** Distribution of Genus

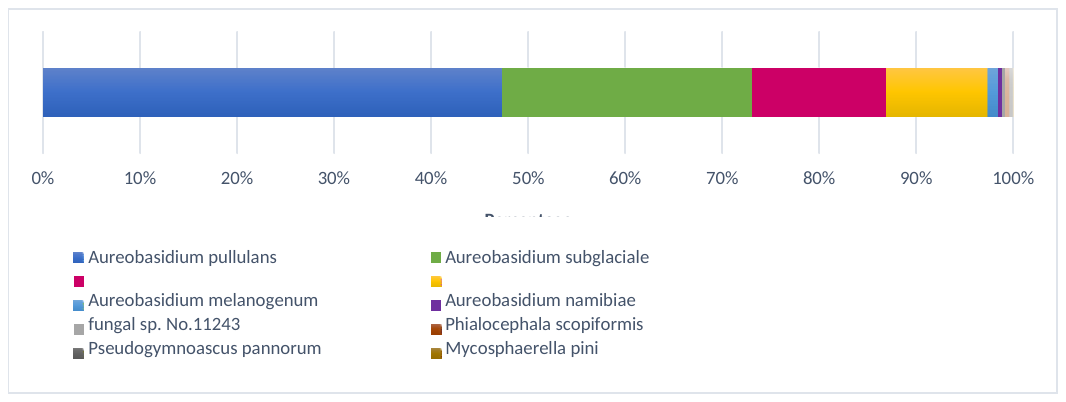

**Figure 4.** Distribution of species
