## Supplementary Data 4 for "Moving Yeasts: Resolving the Mystery"

| species | species_perc |
|---|---|
| root;cellular organisms;Eukaryota;Opisthokonta;Fungi;Dikarya;Ascomycota;saccharomyceta;Pezizomycotina;leotiomyceta;dothideomyceta;Dothideomycetes;Dothideomycetidae;Dothideales;Aureobasidiaceae;Aureobasidium;Aureobasidium pullulans; | 45.287 |
| root;cellular organisms;Eukaryota;Opisthokonta;Fungi;Dikarya;Ascomycota;saccharomyceta;Pezizomycotina;leotiomyceta;dothideomyceta;Dothideomycetes;Dothideomycetidae;Dothideales;Aureobasidiaceae;Aureobasidium;Aureobasidium subglaciale; | 24.594 |
| root;cellular organisms;Eukaryota;Opisthokonta;Fungi;Dikarya;Ascomycota;saccharomyceta;Pezizomycotina;leotiomyceta;dothideomyceta;Dothideomycetes;Dothideomycetidae;Dothideales;Aureobasidiaceae;Aureobasidium;Aureobasidium melanogenum; | 13.218 |
| root;cellular organisms;Eukaryota;Opisthokonta;Fungi;Dikarya;Ascomycota;saccharomyceta;Pezizomycotina;leotiomyceta;dothideomyceta;Dothideomycetes;Dothideomycetidae;Dothideales;Aureobasidiaceae;Aureobasidium;Aureobasidium namibiae; | 9.913 |
| root;cellular organisms;Eukaryota;Opisthokonta;Fungi;unclassified Fungi;fungal sp. No.11243; | 1.083 |
| root;cellular organisms;Eukaryota;Opisthokonta;Fungi;Dikarya;Ascomycota;saccharomyceta;Pezizomycotina;leotiomyceta;sordariomyceta;Leotiomycetes;Helotiales;mitosporic Helotiales;Phialocephala <Helotiales>;Phialocephala scopiformis; | 0.379 |
| root;cellular organisms;Eukaryota;Opisthokonta;Fungi;Dikarya;Ascomycota;saccharomyceta;Pezizomycotina;leotiomyceta;sordariomyceta;Leotiomycetes;Leotiomycetes incertae sedis;Pseudeurotiaceae;Pseudogymnoascus;Pseudogymnoascus pannorum; | 0.379 |
| root;cellular organisms;Eukaryota;Opisthokonta;Fungi;Dikarya;Ascomycota;saccharomyceta;Pezizomycotina;leotiomyceta;dothideomyceta;Dothideomycetes;Dothideomycetidae;Capnodiales;Mycosphaerellaceae;Mycosphaerella;Mycosphaerella pini; | 0.325 |
| root;cellular organisms;Eukaryota;Opisthokonta;Fungi;Dikarya;Ascomycota;saccharomyceta;Pezizomycotina;leotiomyceta;dothideomyceta;Dothideomycetes;Dothideomycetes incertae sedis;Botryosphaeriales;Botryosphaeriaceae;Macrophomina;Macrophomina phaseolina; | 0.271 |
| root;cellular organisms;Eukaryota;Opisthokonta;Fungi;Dikarya;Ascomycota;saccharomyceta;Pezizomycotina;leotiomyceta;dothideomyceta;Dothideomycetes;Dothideomycetidae;Capnodiales;Mycosphaerellaceae;Zymoseptoria;Zymoseptoria brevis; | 0.163 |
| root;cellular organisms;Eukaryota;Opisthokonta;Fungi;Dikarya;Ascomycota;saccharomyceta;Pezizomycotina;leotiomyceta;dothideomyceta;Dothideomycetes;Dothideomycetidae;Capnodiales;Mycosphaerellaceae;Zymoseptoria;Zymoseptoria tritici; | 0.163 |
| root;cellular organisms;Eukaryota;Opisthokonta;Fungi;Dikarya;Ascomycota;saccharomyceta;Pezizomycotina;leotiomyceta;dothideomyceta;Dothideomycetes;Dothideomycetidae;Capnodiales;Teratosphaeriaceae;Baudoinia;Baudoinia compniacensis; | 0.163 |
| root;cellular organisms;Eukaryota;Opisthokonta;Fungi;Dikarya;Ascomycota;saccharomyceta;Pezizomycotina;leotiomyceta;dothideomyceta;Dothideomycetes;Pleosporomycetidae;Pleosporales;Pleosporineae;Pleosporaceae;Stemphylium;Stemphylium lycopersici; | 0.163 |
| root;cellular organisms;Eukaryota;Opisthokonta;Fungi;Dikarya;Ascomycota;saccharomyceta;Pezizomycotina;leotiomyceta;Eurotiomycetes;Chaetothyriomycetidae;Chaetothyriales;Herpotrichiellaceae;Coniosporium;Coniosporium apollinis; | 0.163 |
| root;cellular organisms;Eukaryota;Opisthokonta;Fungi;Dikarya;Ascomycota;saccharomyceta;Pezizomycotina;leotiomyceta;sordariomyceta;Sordariomycetes;Hypocreomycetidae;Hypocreales;Clavicipitaceae;Torrubiella <Clavicipitaceae>;Torrubiella hemipterigena; | 0.163 |
| root;cellular organisms;Eukaryota;Opisthokonta;Fungi;Dikarya;Ascomycota;saccharomyceta;Pezizomycotina;leotiomyceta;sordariomyceta;Sordariomycetes;Hypocreomycetidae;Hypocreales;Nectriaceae;Fusarium;Fusarium solani species complex;Nectria haematococca; | 0.163 |
| root;cellular organisms;Eukaryota;Opisthokonta;Fungi;Dikarya;Ascomycota;saccharomyceta;Pezizomycotina;leotiomyceta;sordariomyceta;Sordariomycetes;Sordariomycetidae;Magnaporthales;Magnaporthaceae;Magnaporthe;Magnaporthe oryzae; | 0.163 |
| root;cellular organisms;Eukaryota;Opisthokonta;Fungi;Dikarya;Ascomycota;saccharomyceta;Pezizomycotina;leotiomyceta;Eurotiomycetes;Chaetothyriomycetidae;Chaetothyriales;Herpotrichiellaceae;Exophiala; | 0.108 |
| root;cellular organisms;Eukaryota;Opisthokonta;Fungi;Dikarya;Ascomycota;saccharomyceta;Pezizomycotina;leotiomyceta;Eurotiomycetes;Eurotiomycetidae;Eurotiales;Aspergillaceae;Penicillium;Penicillium camemberti; | 0.108 |
| root;cellular organisms;Eukaryota;Opisthokonta;Fungi;Dikarya;Ascomycota;saccharomyceta;Pezizomycotina;leotiomyceta;sordariomyceta;Sordariomycetes;Hypocreomycetidae;Hypocreales;Hypocreaceae;Trichoderma;Trichoderma gamsii; | 0.108 |
| root;cellular organisms;Eukaryota;Opisthokonta;Fungi;Dikarya;Ascomycota;saccharomyceta;Pezizomycotina;leotiomyceta;sordariomyceta;Sordariomycetes;Hypocreomycetidae;Hypocreales;Nectriaceae;Fusarium;Fusarium fujikuroi species complex;Fusarium fujikuroi; | 0.108 |
| root;cellular organisms;Eukaryota;Opisthokonta;Fungi;Dikarya;Ascomycota;saccharomyceta;Pezizomycotina;leotiomyceta;sordariomyceta;Sordariomycetes;Sordariomycetidae;Ophiostomatales;Ophiostomataceae;Grosmannia;Grosmannia clavigera; | 0.108 |
| root;cellular organisms;Bacteria;Actinobacteria <phylum>;Actinobacteria;Actinobacteridae;Actinomycetales;Micrococcineae;Microbacteriaceae;Agrococcus; | 0.054 |
| root;cellular organisms;Bacteria;Actinobacteria <phylum>;Actinobacteria;Actinobacteridae;Actinomycetales;Micrococcineae;Microbacteriaceae;Microbacterium; | 0.054 |
| root;cellular organisms;Bacteria;Actinobacteria <phylum>;Actinobacteria;Actinobacteridae;Actinomycetales;Micrococcineae;Micrococcaceae;Micrococcus;Micrococcus luteus; | 0.054 |
| root;cellular organisms;Bacteria;Actinobacteria <phylum>;Actinobacteria;Actinobacteridae;Actinomycetales;Propionibacterineae;Propionibacteriaceae;Propionibacterium; | 0.054 |
| root;cellular organisms;Bacteria;Actinobacteria <phylum>;Actinobacteria;Actinobacteridae;Actinomycetales;Pseudonocardineae;Pseudonocardiaceae; | 0.054 |
| root;cellular organisms;Bacteria;Bacteroidetes/Chlorobi group;Bacteroidetes; | 0.054 |
| root;cellular organisms;Bacteria;Deinococcus-Thermus;Deinococci;Deinococcales;Deinococcaceae;Deinococcus; | 0.054 |
| root;cellular organisms;Bacteria;Proteobacteria;Alphaproteobacteria;Rhizobiales;Bradyrhizobiaceae;Bradyrhizobium;Bradyrhizobium japonicum; | 0.054 |
| root;cellular organisms;Bacteria;Proteobacteria;Betaproteobacteria;Burkholderiales;Burkholderiaceae;Ralstonia; | 0.054 |
| root;cellular organisms;Bacteria;Proteobacteria;Gammaproteobacteria;Enterobacteriales;Enterobacteriaceae; | 0.054 |
| root;cellular organisms;Bacteria;Proteobacteria;Gammaproteobacteria;Pseudomonadales;Moraxellaceae;Acinetobacter;Acinetobacter junii; | 0.054 |
| root;cellular organisms;Bacteria;Proteobacteria;Gammaproteobacteria;Pseudomonadales;Pseudomonadaceae;Pseudomonas; | 0.054 |
| root;cellular organisms;Bacteria;Proteobacteria;Gammaproteobacteria;Xanthomonadales;Xanthomonadaceae; | 0.054 |
| root;cellular organisms;Eukaryota;Alveolata;Apicomplexa;Aconoidasida;Haemosporida;Plasmodium;Plasmodium (Plasmodium);Plasmodium inui; | 0.054 |
| root;cellular organisms;Eukaryota;Opisthokonta;Fungi;Dikarya;Ascomycota;saccharomyceta;Pezizomycotina;leotiomyceta;dothideomyceta;Dothideomycetes;Dothideomycetes incertae sedis;Botryosphaeriales;Botryosphaeriaceae;Diplodia;Diplodia seriata; | 0.054 |
| root;cellular organisms;Eukaryota;Opisthokonta;Fungi;Dikarya;Ascomycota;saccharomyceta;Pezizomycotina;leotiomyceta;dothideomyceta;Dothideomycetes;Pleosporomycetidae;Pleosporales;Pleosporineae;Leptosphaeriaceae;Leptosphaeria;Leptosphaeria maculans complex;Leptosphaeria maculans; | 0.054 |
| root;cellular organisms;Eukaryota;Opisthokonta;Fungi;Dikarya;Ascomycota;saccharomyceta;Pezizomycotina;leotiomyceta;Eurotiomycetes;Chaetothyriomycetidae;Chaetothyriales;Cyphellophoraceae;Cyphellophora;Cyphellophora europaea; | 0.054 |
| root;cellular organisms;Eukaryota;Opisthokonta;Fungi;Dikarya;Ascomycota;saccharomyceta;Pezizomycotina;leotiomyceta;Eurotiomycetes;Chaetothyriomycetidae;Chaetothyriales;Herpotrichiellaceae;Capronia;Capronia coronata; | 0.054 |
| root;cellular organisms;Eukaryota;Opisthokonta;Fungi;Dikarya;Ascomycota;saccharomyceta;Pezizomycotina;leotiomyceta;Eurotiomycetes;Chaetothyriomycetidae;Chaetothyriales;Herpotrichiellaceae;Cladophialophora;Cladophialophora psammophila; | 0.054 |
| root;cellular organisms;Eukaryota;Opisthokonta;Fungi;Dikarya;Ascomycota;saccharomyceta;Pezizomycotina;leotiomyceta;Eurotiomycetes;Chaetothyriomycetidae;Chaetothyriales;Herpotrichiellaceae;Rhinocladiella;Rhinocladiella mackenziei; | 0.054 |
| root;cellular organisms;Eukaryota;Opisthokonta;Fungi;Dikarya;Ascomycota;saccharomyceta;Pezizomycotina;leotiomyceta;Eurotiomycetes;Eurotiomycetidae;Eurotiales;Aspergillaceae;Aspergillus;Aspergillus calidoustus; | 0.054 |
| root;cellular organisms;Eukaryota;Opisthokonta;Fungi;Dikarya;Ascomycota;saccharomyceta;Pezizomycotina;leotiomyceta;Eurotiomycetes;Eurotiomycetidae;Eurotiales;Aspergillaceae;Aspergillus;Aspergillus fumigatus; | 0.054 |
| root;cellular organisms;Eukaryota;Opisthokonta;Fungi;Dikarya;Ascomycota;saccharomyceta;Pezizomycotina;leotiomyceta;Eurotiomycetes;Eurotiomycetidae;Eurotiales;Aspergillaceae;Aspergillus;Aspergillus terreus; | 0.054 |
| root;cellular organisms;Eukaryota;Opisthokonta;Fungi;Dikarya;Ascomycota;saccharomyceta;Pezizomycotina;leotiomyceta;Eurotiomycetes;Eurotiomycetidae;Eurotiales;Aspergillaceae;Penicillium;Penicillium brasilianum; | 0.054 |
| root;cellular organisms;Eukaryota;Opisthokonta;Fungi;Dikarya;Ascomycota;saccharomyceta;Pezizomycotina;leotiomyceta;Eurotiomycetes;Eurotiomycetidae;Eurotiales;Aspergillaceae;Penicillium;Penicillium chrysogenum complex;Penicillium rubens; | 0.054 |
| root;cellular organisms;Eukaryota;Opisthokonta;Fungi;Dikarya;Ascomycota;saccharomyceta;Pezizomycotina;leotiomyceta;Eurotiomycetes;Eurotiomycetidae;Eurotiales;Aspergillaceae;Penicillium;Penicillium expansum; | 0.054 |
| root;cellular organisms;Eukaryota;Opisthokonta;Fungi;Dikarya;Ascomycota;saccharomyceta;Pezizomycotina;leotiomyceta;Eurotiomycetes;Eurotiomycetidae;Eurotiales;Trichocomaceae;Rasamsonia;Rasamsonia emersonii; | 0.054 |
| root;cellular organisms;Eukaryota;Opisthokonta;Fungi;Dikarya;Ascomycota;saccharomyceta;Pezizomycotina;leotiomyceta;Eurotiomycetes;Eurotiomycetidae;Eurotiales;Trichocomaceae;Talaromyces;Talaromyces cellulolyticus; | 0.054 |

| | |
|---|---|
| root;cellular organisms;Eukaryota;Opisthokonta;Fungi;Dikarya;Ascomycota;saccharomyceta;Pezizomycotina;leotiomyceta;Eurotiomycetes;Eurotiomycetidae;Eurotiales;Trichocomaceae;Talaromyces;Talaromyces islandicus; | 0.054 |
| root;cellular organisms;Eukaryota;Opisthokonta;Fungi;Dikarya;Ascomycota;saccharomyceta;Pezizomycotina;leotiomyceta;Eurotiomycetes;Eurotiomycetidae;Eurotiales;Trichocomaceae;Talaromyces;Talaromyces stipitatus; | 0.054 |
| root;cellular organisms;Eukaryota;Opisthokonta;Fungi;Dikarya;Ascomycota;saccharomyceta;Pezizomycotina;leotiomyceta;Eurotiomycetes;Eurotiomycetidae;Onygenales;Arthrodermataceae;Microsporum;Microsporum gypseum; | 0.054 |
| root;cellular organisms;Eukaryota;Opisthokonta;Fungi;Dikarya;Ascomycota;saccharomyceta;Pezizomycotina;leotiomyceta;sordariomyceta;Leotiomycetes;Helotiales;Sclerotiniaceae;Botrytis;Botrytis cinerea; | 0.054 |
| root;cellular organisms;Eukaryota;Opisthokonta;Fungi;Dikarya;Ascomycota;saccharomyceta;Pezizomycotina;leotiomyceta;sordariomyceta;Leotiomycetes;Helotiales;Sclerotiniaceae;Sclerotinia;Sclerotinia sclerotiorum; | 0.054 |
| root;cellular organisms;Eukaryota;Opisthokonta;Fungi;Dikarya;Ascomycota;saccharomyceta;Pezizomycotina;leotiomyceta;sordariomyceta;Leotiomycetes;Leotiomycetes incertae sedis;Myxotrichaceae;mitosporic Myxotrichaceae;Oidiodendron;Oidiodendron maius; | 0.054 |
| root;cellular organisms;Eukaryota;Opisthokonta;Fungi;Dikarya;Ascomycota;saccharomyceta;Pezizomycotina;leotiomyceta;sordariomyceta;Sordariomycetes;Hypocreomycetidae;Glomerellales;Glomerellaceae;Colletotrichum;Colletotrichum fioriniae; | 0.054 |
| root;cellular organisms;Eukaryota;Opisthokonta;Fungi;Dikarya;Ascomycota;saccharomyceta;Pezizomycotina;leotiomyceta;sordariomyceta;Sordariomycetes;Hypocreomycetidae;Glomerellales;Glomerellaceae;Colletotrichum;Colletotrichum gloeosporioides; | 0.054 |
| root;cellular organisms;Eukaryota;Opisthokonta;Fungi;Dikarya;Ascomycota;saccharomyceta;Pezizomycotina;leotiomyceta;sordariomyceta;Sordariomycetes;Hypocreomycetidae;Glomerellales;Glomerellaceae;Colletotrichum;Colletotrichum higginsianum; | 0.054 |
| root;cellular organisms;Eukaryota;Opisthokonta;Fungi;Dikarya;Ascomycota;saccharomyceta;Pezizomycotina;leotiomyceta;sordariomyceta;Sordariomycetes;Hypocreomycetidae;Hypocreales;Clavicipitaceae;Metacordyceps;Metacordyceps taii; | 0.054 |
| root;cellular organisms;Eukaryota;Opisthokonta;Fungi;Dikarya;Ascomycota;saccharomyceta;Pezizomycotina;leotiomyceta;sordariomyceta;Sordariomycetes;Hypocreomycetidae;Hypocreales;Cordycipitaceae;Beauveria;Beauveria bassiana; | 0.054 |
| root;cellular organisms;Eukaryota;Opisthokonta;Fungi;Dikarya;Ascomycota;saccharomyceta;Pezizomycotina;leotiomyceta;sordariomyceta;Sordariomycetes;Hypocreomycetidae;Hypocreales;Hypocreaceae;Trichoderma;Trichoderma reesei; | 0.054 |
| root;cellular organisms;Eukaryota;Opisthokonta;Fungi;Dikarya;Ascomycota;saccharomyceta;Pezizomycotina;leotiomyceta;sordariomyceta;Sordariomycetes;Hypocreomycetidae;Hypocreales;Nectriaceae;Fusarium;Fusarium oxysporum species complex;Fusarium oxysporum; | 0.054 |
| root;cellular organisms;Eukaryota;Opisthokonta;Fungi;Dikarya;Ascomycota;saccharomyceta;Pezizomycotina;leotiomyceta;sordariomyceta;Sordariomycetes;Hypocreomycetidae;Hypocreales;Nectriaceae;Fusarium;Fusarium sambucinum species complex;Fusarium langsethiae; | 0.054 |
| root;cellular organisms;Eukaryota;Opisthokonta;Fungi;Dikarya;Ascomycota;saccharomyceta;Pezizomycotina;leotiomyceta;sordariomyceta;Sordariomycetes;Hypocreomycetidae;Hypocreales;Ophiocordycipitaceae;Ophiocordyceps;Ophiocordyceps unilateralis; | 0.054 |
| root;cellular organisms;Eukaryota;Opisthokonta;Fungi;Dikarya;Ascomycota;saccharomyceta;Pezizomycotina;leotiomyceta;sordariomyceta;Sordariomycetes;Sordariomycetidae;Sordariales;Lasiosphaeriaceae;Podospora; | 0.054 |
| root;cellular organisms;Eukaryota;Opisthokonta;Fungi;Dikarya;Ascomycota;saccharomyceta;Pezizomycotina;leotiomyceta;sordariomyceta;Sordariomycetes;Xylariomycetidae;Xylariales;Amphisphaeriaceae;Pestalotiopsis;Pestalotiopsis fici; | 0.054 |
| root;cellular organisms;Eukaryota;Opisthokonta;Fungi;Dikarya;Basidiomycota;Agaricomycotina;Agaricomycetes;Agaricomycetidae;Agaricales;Marasmiaceae;mitosporic Marasmiaceae;Moniliophthora;Moniliophthora roreri; | 0.054 |
| root;cellular organisms;Eukaryota;Opisthokonta;Fungi;Dikarya;Basidiomycota;Agaricomycotina;Tremellomycetes;Tremellales;mitosporic Tremellales;Trichosporon;Trichosporon oleaginosus; | 0.054 |
| root;cellular organisms;Eukaryota;Opisthokonta;Metazoa;Eumetazoa;Bilateria;Deuterostomia;Chordata;Craniata;Vertebrata;Gnathostomata;Teleostomi;Euteleostomi;Actinopterygii;Actinopteri;Neopterygii;Teleostei;Osteoglossocephalai;Clupeocephala;Euteleosteomorpha;Neoteleostei;Eurypterygia;Ctenosquamata;Acanthomorphata;Euacanthomorph | 0.054 |
| root;cellular organisms;Eukaryota;Opisthokonta;Metazoa;Eumetazoa;Bilateria;Deuterostomia;Chordata;Craniata;Vertebrata;Gnathostomata;Teleostomi;Euteleostomi;Sarcopterygii;Dipnotetrapodomorpha;Tetrapoda;Amniota;Mammalia;Theria;Eutheria;Boreoeutheria;Euarchontoglires;Glires;Rodentia;Sciurognathi;Sciuridae;Xerinae;Marmotini;Ictidom | 0.054 |
| root;cellular organisms;Eukaryota;Opisthokonta;Metazoa;Eumetazoa;Bilateria;Protostomia;Ecdysozoa;Panarthropoda;Arthropoda;Mandibulata;Pancrustacea;Hexapoda;Insecta;Dicondylia;Pterygota;Neoptera;Endopterygota;Diptera;Brachycera;Muscomorpha;Eremoneura;Cyclorrhapha;Schizophora;Acalyptratae;Ephydroidea;Drosophilidae;Drosophilin | 0.054 |
