## Supplementary Videos with captions for "Moving Yeasts: Resolving the Mystery"

Captions and Link (restricted access) for **Supplementary Videos**

Supplementary Video 1 (<https://youtu.be/y9lMPImsG-M>)

Displacement of Yeast (*A.tremulum)*. **Left Frame** indicating the presence of two yeasts wherein the blue arrow shows non-moving yeast while the black arrow indicates moving yeast. **Right frame** shows the tracking of moving yeast.

Supplementary Video 2 (<https://youtu.be/IzDrj9QYelY>)

Displacement of Yeast (*A.tremulum)*. **Left frame** indicating the presence of many yeasts in the microscopic field wherein many are moving from one place to another. Some are trembling at their place. The red arrow indicates bacteria outside the yeast cells. The red box indicates bacteria present inside the yeast cell. The Blue arrow indicates slightly moving yeast while the black arrow indicates moving yeast. **Right frame** shows the tracking of moving yeast.

Supplementary Video 3a (<https://youtu.be/Vhiswrr6FwU>)

Trembling behavior of *A.tremulum* because of the Bacterial struggle in crossing the EPS

Supplementary Video 3b (<https://youtu.be/kiEQ-6Uokzk>)

Displacement of yeast (*A.tremulum*) and release of bacteria. A white-colored box in the video indicates the yeast undergoing displacement. Released bacteria from yeast have been indicated by white arrow. The black arrow (at 4:19 minutes) indicates protrusion formation due to bacterial release.

Supplementary Video 4 (<https://youtu.be/YTwDIHOyAIc>)

Phase-contrast view of *A. tremulum*. The video shows trembling yeast cells. Some of them also contain lipid droplets. Some of lipid droplets are present outside of yeast. Small moving bacteria can also be observed outside yeast cells.

Supplementary Video 5 (<https://youtu.be/5SlLCi7-nkM>)

Trembling movement of *A.tremulum* stained with DAPI showing intracellular bacteria

Supplementary Video 6 (<https://youtu.be/pRjWP3BqXoY>)

Confocal Microscopic view of *A.tremulum*. The video shows presence of lipid droplets and bacteria inside yeast cell. The vigourously moving lipid droplets and slowly moving bacteria inside yeast can be easily observed. Bottom frame indicates the tracking of lipid droplet movemennnt inside yeast and bacterial movement (at right corner) outside yeast.

Supplementary Video 7 (<https://youtu.be/XUmozFFq8KM>)

Confocal Microscopic view of *A.tremulum*. The video shows presence of lipid droplets inside yeast cell.

Supplementary Video 8 (<https://youtu.be/U4c9dSKgLPY>)

Comparative Video showing movement of yeast (grown on PDA) with lipid (**right frame**) and without lipid droplets (**left frame**). Lipid droplet containing yeast are taken on 14^th^ day of incubation while yeast without lipid droplet are taken on 3^rd^ day of incubation.
